# Connectome-seq: High-throughput Mapping of Neuronal Connectivity at Single-Synapse Resolution via Barcode Sequencing

**DOI:** 10.1101/2025.02.13.638129

**Authors:** Danping Chen, Alina Isakova, Zhou Wan, Mark J. Wagner, Yunming Wu, Boxuan Simen Zhao

## Abstract

Understanding neuronal connectivity at single-cell resolution remains a fundamental challenge in neuroscience, with current methods particularly limited in mapping long-distance circuits and preserving cell type information. Here we present Connectome-seq, a high-throughput method that combines engineered synaptic proteins, RNA barcoding, and parallel single-nucleus and single-synaptosome sequencing to map neuronal connectivity at single-synapse resolution. This AAV-based approach enables simultaneous capture of both synaptic connections and molecular identities of connected neurons. We validated this approach in the mouse pontocerebellar circuit, identifying both established projections and potentially novel synaptic partnerships. Through integrated analysis of connectivity and gene expression, we identified molecular markers enriched in connected neurons, suggesting potential associated factors of circuit organization. By enabling systematic mapping of neuronal connectivity across brain regions with single-cell precision and gene expression information, Connectome-seq provides a scalable platform for comprehensive circuit analysis across different experimental conditions and biological states. This advance in connectivity mapping methodology opens new possibilities for understanding circuit organization in complex mammalian brains.

## Introduction

Understanding synaptic wiring diagrams of the brain, known as the connectomes, is a fundamental goal in neuroscience. This complex network of neuronal connections underlies all brain functions, from simple reflexes to complex cognitive processes. Breakthroughs in connectomics have provided unprecedented insights into brain organization. An early breakthrough was the complete reconstruction of the *C. elegans* connectome^1^, representing the first comprehensive map of a nervous system with its approximately 300 neurons and their synaptic connections. Recent achievements include the complete reconstruction of the adult *Drosophila* brain connectome, which mapped approximately 140,000 neurons and over 50 million connections, representing the largest connectome obtained to date^2–5^. The *Drosophila* connectome already has a profound effect on neuroscience, enabling researchers to trace neural circuits, generate hypotheses about their function, and develop circuit models rooted in actual connectivity. This resource exemplifies the transformative potential of comprehensive connectome data.

Serial-section electron microscopy (ssEM) has been the foundational method enabling these comprehensive connectome reconstructions in invertebrates. Adaptations of ssEM techniques for the more complex vertebrate nervous system have also driven major progress in mapping neuronal connectivity at single-cell resolution in the vertebrate brain. Early and highly influential work using ssEM in the retina has provided critical insights into the neural circuitry underlying specific functions, such as direction selectivity in retinal ganglion cells^6–8^. More recently, dense reconstructions of various brain regions have revealed fundamental principles of circuit organization. For example, study in the medial entorhinal cortex demonstrated precise axonal synapse sorting that supports temporal processing of information^9^. Dense reconstructions of adult mouse somatosensory cortex uncovered novel distance-dependent axonal targeting principles of inhibitory and excitatory connectivity^10^, while developmental studies in the same region revealed how inhibitory synaptic specificity emerges through distinct postnatal trajectories^11^. Comprehensive analysis of mouse visual cortex demonstrated structured functional connectivity rules governing inter-areal projections^12^, subsequent work showed widespread like-to-like connectivity principles across cortical layers and areas^13^, and detailed studies of thick-tufted layer 5 pyramidal cells revealed cell-type specific organization of proximal and extended connectivity patterns^14^. Cerebellar circuit mapping revealed how non-random redundant connectivity motifs help balance pattern separation with noise resilience^15^. While these studies have provided unprecedented insights into local circuit architecture, they remain fundamentally limited to volumes of a few hundred micrometers due to the technical challenges of imaging and reconstructing larger tissue volumes. Moreover, while invertebrate circuits like those in *C. elegans* and *Drosophila* exhibit a higher degree of stereotypy compared to mammals, it is now recognized that these circuits also possess individual variability^16,17^. Mammalian circuits exhibit a greater magnitude of individual variation in connectivity at the single-cell level. This biological variability means that studying mammalian circuits requires methods that are both practical and scalable enough to characterize connectivity in individual subjects, especially when aiming to correlate circuit structure with neural dynamics or behavior.

Meeting this challenge requires new approaches that can achieve both high throughput mapping and single-cell resolution in larger mammalian brains. Various techniques exist for mapping mammalian circuit connectivity at non-single-cell scales^18,19^ or single cells at low throughput^20–22^. Light microscopy-based methods like mGRASP^23^ and SynView^24^ enable visualization of synaptic contacts between specific neuronal populations, while genetically modified viral tracers allow circuit component identification without requiring direct synaptic visualization. However, these approaches face important limitations - light microscopy often lacks the resolution to definitively identify synapses, viral tracing can be constrained by tropism and potential toxicity, and most importantly, these methods are inherently designed to examine a specific subset of connections based on genetic markers or viral targeting, making comprehensive, unbiased circuit mapping challenging.

Recent advances in sequencing-based approaches have transformed our ability to map neural connectivity at scale. MAPseq pioneered this revolution by using barcoded Sindbis virus to trace thousands of neuronal projections simultaneously^25^, followed by BARseq which preserved spatial context through in situ sequencing^26–29^, and BRICseq which enabled multiplexed tracing from multiple sites^30^. Building on these advances, researchers developed approaches using barcoded rabies virus to map synaptic connectivity. Notable developments include RABID-seq’s demonstration of the concept in glial cells^31^, SBARRO’s proof-of-principle in controlled sparse neuronal cultures^32^, and recent work that revealed cell type-specific connectivity patterns in cortical circuits^33,34^. However, these methods face significant technical limitations: Sindbis virus-based approaches cannot reveal post-synaptic partners and exhibit severe toxicity due to high-level viral payload expression, typically killing cells within days; whereas rabies-based methods struggle with barcode mixing in densely connected neurons and viral toxicity that may compromise circuit integrity.

To address these challenges, we developed Connectome-seq, an approach that attempts to overcome key limitations of existing methods while offering unique advantages for circuit mapping. Leveraging minimally toxic adeno-associated viruses (AAV) vectors and combining molecular engineering with high-throughput sequencing, Connectome-seq enables unbiased mapping of neuronal connections at single-synapse resolution across large brain volumes. This approach simultaneously captures neuronal identity, gene expression, and synaptic connectivity without the need for pre-selection of specific cell types or limitations based on cellular density.

We demonstrated the utility of Connectome-seq in the mouse pontocerebellar circuit, a universally conserved circuit in mammals that exemplifies the challenges of mapping long-distance connections in the mammalian brain. Connectome-seq identified thousands of connections across this circuit, including both known projections and potentially novel synaptic partnerships. The high-throughput, unbiased nature of our method enables comprehensive analysis of connectivity patterns while preserving critical information about cellular identity and molecular characteristics. This combination of features makes Connectome-seq particularly well-suited for systematic circuit mapping across different experimental conditions and biological states.

## Results

The Connectome-seq workflow consists of three key steps: (1) molecular engineering of synaptic barcoding constructs and viral delivery of transgenes into target brain regions, (2) development of a parallel single-nucleus and single-synaptosome sequencing protocol, and (3) computational reconstruction of the connectome. Here, we present the development process and detailed results in each step, highlighting the advances that enable high-throughput synaptic mapping, and the biological insights gained from applying this method to map the mouse pontocerebellar circuit.

### 1. Molecular engineering to translate a synapse into a pair of RNA barcodes

Connectome-seq relies on the labeling of synaptic connections using engineered protein-RNA complexes to translate a synapse into a pair of unique RNA barcodes. The intended outcome is that one barcode will be associated with the pre-synaptic neuron and the other with the post-synaptic neuron, allowing for later profiling via high-throughput sequencing (Fig. 1A). Two protein anchors, termed SynBar (synaptic barcoding), were designed: PreSynBar to target presynaptic terminals and PostSynBar to target postsynaptic sites (Fig. 1B and Extended Data Fig. 1A). PreSynBar consists of a modified neurexin 1β protein fused to the large fragment of split-GFP (GFP1-10) and an RNA-binding domain, λN22 peptide, similar to other reported designs^25,35^. PostSynBar is based on neuroligin 1, also fused to the complementary small fragment of split-GFP (GFP11) and the same RNA-binding domain. These trans-membrane protein constructs are designed to interact at synapses, reconstituting GFP and bringing together their associated RNA barcodes to anchor at the synaptic membranes. The protein scaffolds were largely adopted from the SynView constructs^24^, where split-GFP fragments are embedded inside the synaptic proteins rather than at their N-terminals. This design choice aims to ensure GFP reconstitution occurs specifically upon proper synaptic protein interaction. An epitope tag (V5 for PreSynBar and HA for PostSynBar) was added to the extracellular domain of each construct to visualize tagged synapses via antibody staining.

**Figure 1.**
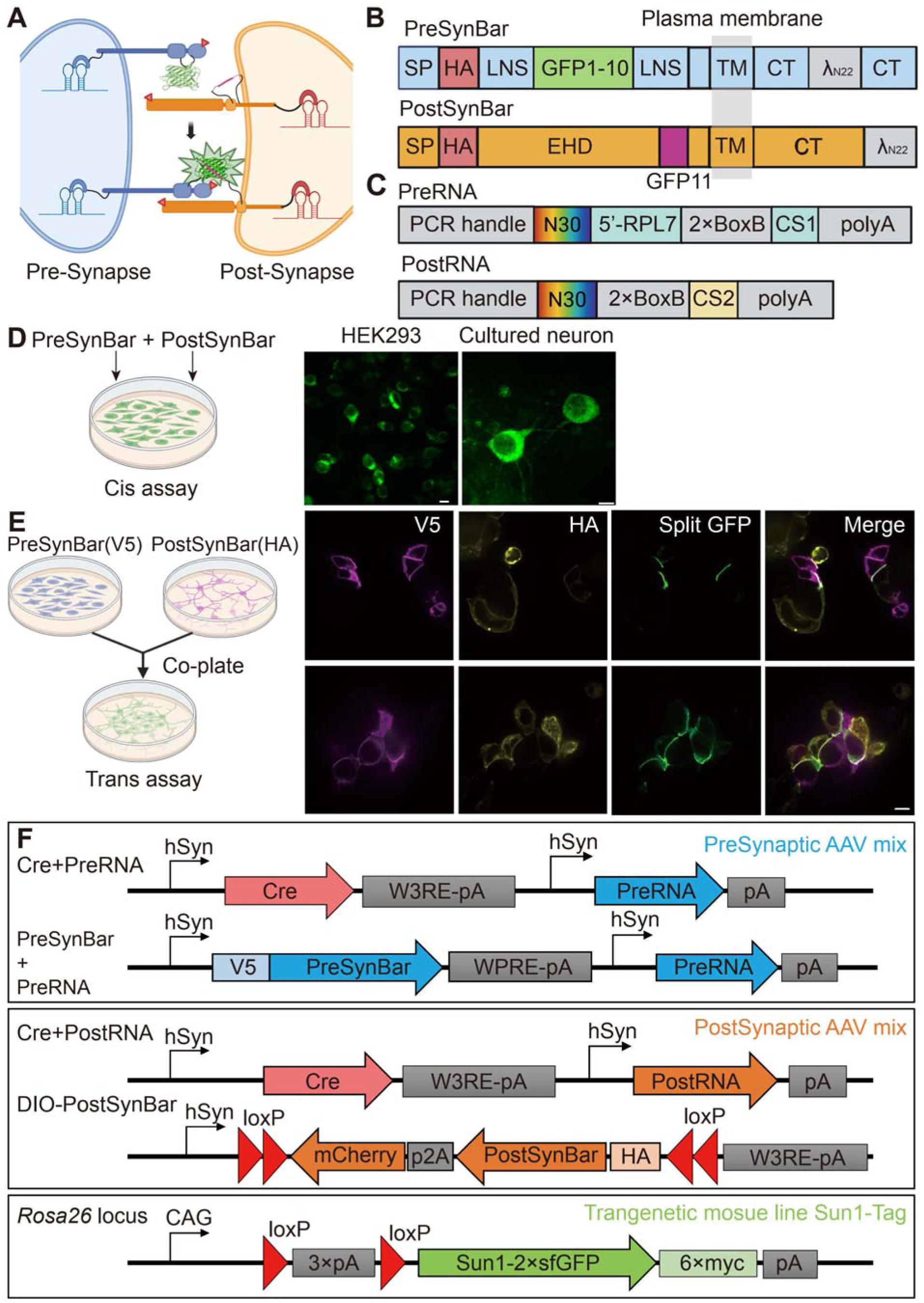
Design and validation of SynBar constructs, viral vectors, and the transgenic mouse line used in Connectome-seq. (A) Schematic illustration of the SynBar system at synapses. PreSynBar and PostSynBar proteins interact to form split-GFP and bring RNA barcodes together. (B) Schematic representation of PreSynBar and PostSynBar proteins, showing the neurexin/neuroligin base, split-GFP components, and RNA-binding domains. GFP1–10 and GFP11 denote the split-GFP moieties containing β-strands 1–10 and 11, respectively. SP, Signal peptide; EHD, Esterase homology domain; LNS, laminin/neurexin/sex hormone binding domain; TM, transmembrane domain; CT, cytoplasmic tail. (C) Design of PreRNA and PostRNA RNA. Key functional elements include PCR handles, N30 random region for unique cell of origin identification, BoxB sites for λN22 binding, and specific capture sequences (CS1/2) for 10X library construction. RPL7 5 UTR in PreRNA enhances distal transport. (D) Cis validation of SynBar interaction in HEK293 cells and rat cortical neurons showing reconstituted split GFP signals (green) inside cells expressing both PreSynBar and PostSynBar (cis-interaction). Scale bars: 10 μm. (E) Trans validation showing specific GFP reconstitution (green) at contact surface between HEK293 cells expressing complementary PreSynBar (magenta, via V5 staining) and PostSynBar (yellow, via HA staining) constructs. Two sets of representative images shown. Scale bar: 10 μm. (F) Design of viral constructs and transgenic mouse line used in Connectome-seq. Top: Presynaptic AAV mix containing two viruses - a Cre-PreRNA virus and a PreSynBar-PreRNA virus, both driven by two hSyn promoter cassettes. Middle: Postsynaptic AAV mix containing Cre-PostRNA virus and Cre-dependent (DIO) PostSynBar virus. Bottom: Sun1-Tag transgenic mouse line with Cre-dependent Sun1-2×sfGFP fusion protein inserted at the *Rosa26* locus.

An important design consideration for PreSynBar was the positioning of the λN22 RNA-binding domain. While PostSynBar tolerated the addition of λN22 at its extreme C-terminus, PreSynBar required internal placement within its C-terminal region. In trans-interaction assays, PreSynBar with λN22 at the extreme C-terminus failed to show proper interaction with PostSynBar, while PreSynBar with internal C-terminal λN22 showed evidence of trans-interaction (Extended Data Fig. 1B). This finding is consistent with previous studies showing the sensitivity of neurexin’s C-terminal region to modifications, suggesting the importance of preserving its native structure for proper membrane targeting.

Two corresponding RNA barcodes, named PreRNA and PostRNA, were designed with the aim of efficient expression, synaptic localization, and detection by sequencing (Fig. 1C). Each barcode contains several domains designed to perform specific functions: PCR handles for amplification, a 30-nucleotide random region (N30) with sufficient complexity to yield unique sequences for each neuron, two copies of BoxB sequence that can be tightly bound by the λN22 peptide on SynBar proteins, and a capture sequence (CS1 or CS2) matching the 10x Genomics feature barcoding technology. We used large-scale Golden Gate assembly to insert the random N30 region into the plasmid to create a highly complex plasmid library that was later packaged into AAV as a complex viral library. To ensure efficient expression and transport, the barcodes include a poly-adenylated region for RNA Polymerase II-driven transcription, which appeared to improve nuclear export and synaptic abundance compared to Pol III-driven expression in our experiments (Extended Data Fig. 1C). To facilitate the necessary long-distance transport of PreRNA to distal presynaptic terminals, we explored redesigning the PreRNA barcodes to naturally localize to neurites. Drawing upon previous research identifying specific 5 and 3 untranslated regions (UTRs) that enhance mRNA localization^36,37^, we tested several UTR modifications for their ability to promote PreRNA abundance in distal regions. Specifically, the addition of the 5 UTR of RPL7 led to the greatest increase in the abundance of PreRNA in both cytosolic and synaptosomal fractions of the postsynaptic region among all designs tested in our experiments, and was therefore incorporated into the final PreRNA design (Extended Data Fig. 1D).

To assess the stability of BoxB-λN22 interaction in our system, we performed RNA immunoprecipitation (RIP) assays using anti-GFP antibodies. The RIP experiments showed >500-fold enrichment of barcodes when using λN22-tagged SynBar proteins compared to untagged controls (SynView), suggesting that this protein-RNA interaction is sufficiently stable to withstand biochemical isolation (Extended Data Fig. 1E).

We further assessed the interaction of PreSynBar and PostSynBar through an established cultured-cell-based strategy used for other synaptic labeling approaches^24^. In cis validation experiments (where both constructs are expressed in the same cell), we observed strong reconstituted split-GFP signals when PreSynBar and PostSynBar were co-expressed within individual HEK 293 cells and cultured neurons (Fig. 1D). For trans validation (where the constructs are expressed in separate cells that are then co-cultured), we examined cells expressing each construct separately: when PreSynBar-expressing cells (detected by V5 staining in cyan) were co-plated with PostSynBar-expressing cells (detected by HA staining in magenta), we observed specific split-GFP reconstitution (green) at points of contact between the two cell populations, in both HEK 293 cells (Fig. 1E) and HEK 293-neuron mixed cultures (Extended Data Fig. 1F). The merged images showed the colocalization of all three signals at these cellular junctions, consistent with protein-protein interaction and GFP reconstitution both within single cells (cis) and between adjacent cells (trans).

### 2. Development of Parallel Nuclear and Synaptosome Isolation Strategy

A key feature of Connectome-seq is the ability to simultaneously capture both cellular identity and synaptic connectivity information. This is achieved through parallel profiling of neuronal nuclei and synaptosomes from the same tissue, where co-existing barcodes in these two structures link each cell to a synapse (Fig. 2A). Nuclei contain both cellular transcriptomes and copies of either PreRNA or PostRNA, allowing identification of cell types and their associated barcodes. During gentle brain tissue homogenization, the cell membranes connecting pre-synaptic terminals to axons and post-synaptic sites to dendrites are physically separated. The exposed membrane surfaces at both the pre- and post-synaptic compartments then reseal, forming structures called synaptosomes^38,39^. Since PreSynBar and PostSynBar proteins are connected across the synaptic cleft, tightly bound to the associated RNA barcodes, these protein-RNA complexes should remain intact as long as the synaptic interface is preserved. This allows synaptosomes to maintain the paired PreRNA and PostRNA information from connected neurons, providing direct evidence of synaptic connections. By matching barcodes between nuclei and synaptosomes, we can link the identity of connected neurons and map their connectivity patterns. To directly visualize barcode transport, we performed PreRNA FISH experiments in cultured neurons and observed clear PreRNA barcode signals both in transit along neuronal processes and at synaptic terminals colocalizing with PreSynBar proteins (Extended Data Fig. 1G). This result demonstrates efficient transport of RNA barcodes into synapse as we designed.

**Figure 2.**
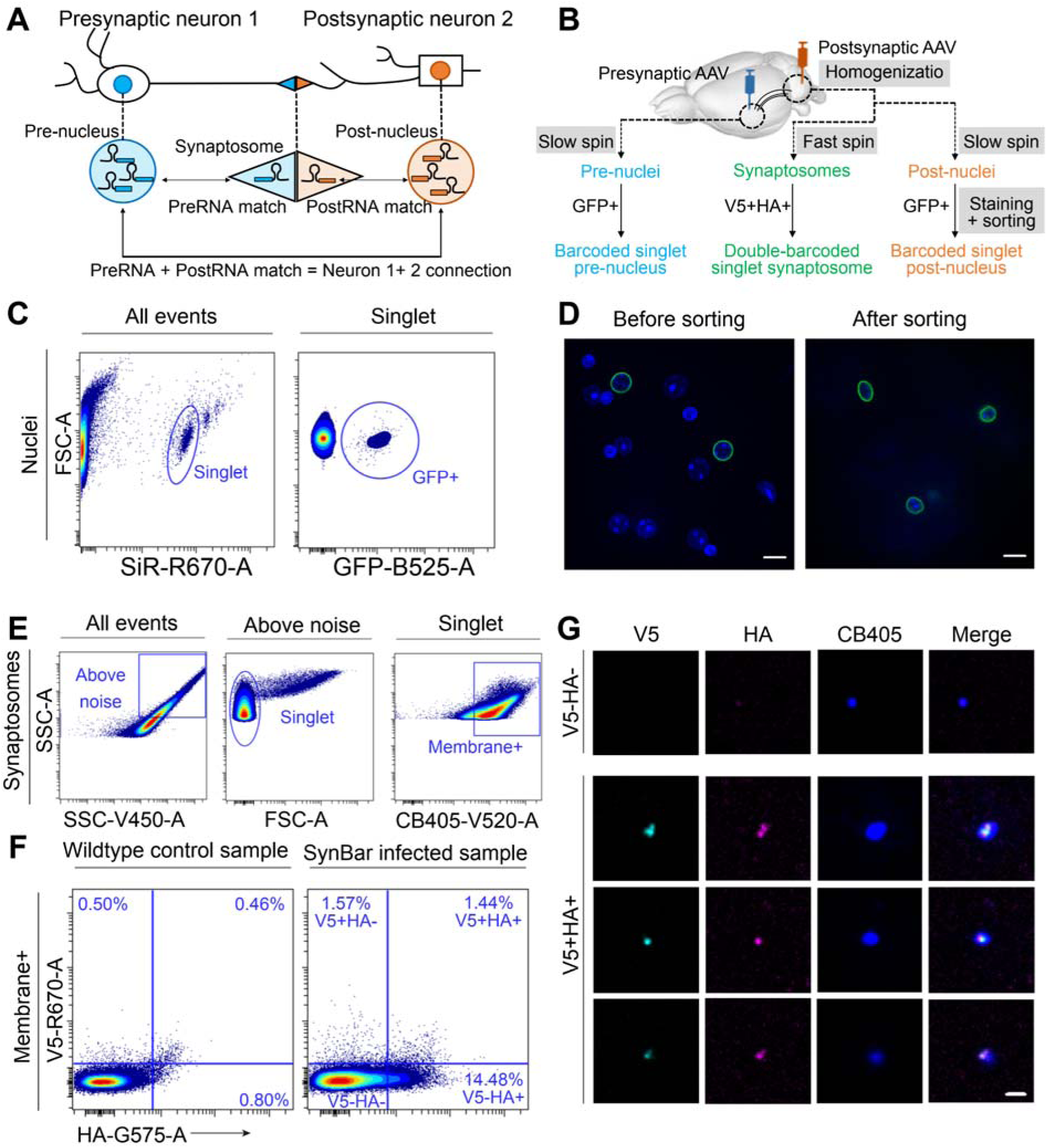
Nuclear and Synaptosome Isolation Strategy. (A) Schematic of the barcode matching strategy. Pre-nucleus contains PreRNA and post-nucleus contains PostRNA from their respective neurons. When both PreRNA and PostRNA sequences from nuclei match those found in a single synaptosome, this indicates a synaptic connection between the two neurons. (B) Workflow for parallel isolation of nuclei and synaptosomes. Following AAV injection into pre- and postsynaptic regions, brain tissue is processed through differential centrifugation. Slow spins yield crude pre- and post-nuclei, while a fast spin of the supernatant of the slow spin produces the synaptosome fraction. Nuclei and synaptosomes are enriched by FACS based on GFP signal (nuclei) or V5+HA antibody staining (synaptosomes). (C) Nuclear isolation strategy. Left: Initial FACS plot showing all events, with gating on SiR-DNA signal identifying singlet nuclei. Right: Gating on singlet nuclei to select GFP+ population. (D) Validation of nuclear sorting. Confocal images showing pre-sort mixture of GFP+ and GFP-nuclei (left) and post-sort enrichment of GFP+ singlet nuclei (right). Scale bars: 10 μm. (E) Sequential gating strategy for synaptosome isolation. Left to right: Selection of events above noise threshold, selection of singlet population based on scattering properties, and identification of membrane-intact singlet synaptosomes based on Cellbrite 405 (CB405) signal. (F) Synaptosome enrichment by antibody staining. FACS plots comparing wildtype control (left) versus SynBar-infected samples (right), showing enriched V5+/HA+ double-positive population in infected samples. (G) Imaging validation of sorted synaptosomes. Confocal images of sorted populations showing V5-HA-double-negative and V5+HA+ double-positive synaptosomes. Blue is from universal membrane stain with CB405. Cyan is PreSynBar signal from staining with anti-V5 antibody. Magenta is PostSynBar signal from staining with anti-HA antibody. Scale bars: 2 μm.

To implement this strategy, we designed AAV constructs carrying Cre recombinase and SynBar components (Fig. 1F). The Cre-expressing virus induces the expression of a Cre-dependent nuclear-localized fluorescent reporter (Sun1-Tag)^40^, labeling nuclei of infected neurons with membrane-anchored GFP to enable their isolation. An important consideration in our virus design was to ensure PreSynBar expression and targeting. Initial versions of PreSynBar virus showed inconsistent protein localization to neuronal processes. We discovered that including two transcriptional cassettes – one driving PreSynBar protein and another expressing PreRNA – improved targeting efficiency (Extended Data Fig. 2A). This two-cassette design consistently achieved better axonal localization compared to single-cassette versions.

We next developed a protocol to purify and enrich both nuclei and synaptosomes from the same brain regions (Fig. 2B): Using gentle homogenization in a sucrose-based buffer (SET buffer), we performed sequential centrifugation to fractionate the homogenate into crude nuclei and synaptosomes and used flow cytometry for fluorescence-based selection to enrich the population containing barcodes.

For nuclear isolation, we used Sun1-Tag GFP signal for specific enrichment. A key consideration was the choice of DNA stain – while violet dye like DAPI and Hoechst are commonly used for nuclear sorting, their spectral overlap with GFP confounds the selection of GFP-positive events. We therefore used SiR-DNA, a far-red DNA stain, to enable clear separation of DNA content signal from GFP fluorescence. Our optimized sorting strategy employed sequential gating (Fig. 2C): First, we selected events based on SiR-DNA signal intensity to enrich for nuclei containing approximately single-genome DNA content, which helped discriminate singlet nuclei from aggregates and debris. Within this singlet population, we then identified GFP-positive nuclei derived from infected neurons. This approach consistently yielded high-quality nuclear preparations, with GFP expression and nuclear singlet status confirmed by confocal microscopy post collection (Fig. 2D). RNA quality is also critical for single-nucleus RNA sequencing. To preserve RNA integrity, we minimized post-sorting handling of nuclei by sorting GFP-positive singlet nuclei directly into a collection buffer and immediately loading the sample onto the 10x Genomics microfluidic chip. This streamlined approach ensured consistent, high-quality nuclear RNA for efficient cell typing.

The next critical component of Connectome-seq is the ability to isolate and enrich double-barcoded synaptosomes. We found that crude synaptosomes (P2 fraction) provided a suitable starting material for our flow cytometry-based enrichment workflow. This approach effectively excluded non-synaptosomal fractions and debris while preserving intact synaptosomes, achieving similar goals to traditional sucrose gradient purification but with reduced processing time. We developed a systematic sequential gating strategy using flow cytometry to enrich synaptosomes based on their size, internal complexity, membrane integrity, and the presence of PreSynBar and PostSynBar-associated fluorescent signals (Fig. 2E). This flow cytometry-based approach proved crucial for reliable synaptosome detection and sorting.

Our initial plan to identify double-barcoded synaptosomes relied on flow cytometry detection of reconstituted split-GFP signal from interacting PreSynBar and PostSynBar proteins. However, we discovered that the native fluorescent signal fell below the detection sensitivity of standard flow cytometers. This limitation led us to explore several alternative approaches. First, we used axon- and dendrite-targeted fluorescent proteins (PreTag-GFP and PostTag-RFP, Extended Data Fig. 2B, C) for synaptosome labeling. This approach provided bright fluorescent signals suitable for flow cytometry and enabled the enrichment of connected synaptosomes via co-existing green and red fluorescence (Extended Data Fig. 2D). Confocal imaging of sorted double-positive synaptosomes revealed that most structures were true singlets containing both fluorescent proteins, verifying the robustness of our sequential gating strategy (Extended Data Fig. 2E). However, subsequent analysis of ambient barcode RNA showed high levels of contaminating PreRNA and PostRNA among sorted structures, specifically, a high level of PreRNA in postsynaptic nuclei (Extended Data Fig. 2F), indicating that fluorescent proteins not strictly localized to synaptic terminals cannot be used as synaptic barcoding complexes. This finding highlighted the importance of using synapse-specific protein anchors like our SynBar system.

We next explored immunostaining approaches for SynBar detection. We first fluorophore-conjugated a commercially available antibody specific only to reconstituted GFP but not the large split GFP fragment. This custom-conjugated antibody effectively detected GFP+ synaptosomes by flow cytometry (Extended Data Fig. 3A), and the enriched population was confirmed as true singlet by confocal imaging (Extended Data Fig. 3B), but batch variation made it impractical for routine use. We therefore engineered V5 and HA epitope tags into the extracellular domains of PreSynBar and PostSynBar proteins (Fig. 2B), enabling detection with commercially available anti-V5-AlexaFluor 647 and anti-HA-PE antibodies. A critical optimization revealed that traditional sucrose-based SET buffer severely reduced antibody staining efficiency; switching to a PBS-based protocol provided optimal balance between staining efficiency and synaptosome preservation (Extended Data Fig. 3C).

Our final optimized sorting protocol employed sequential gating based on stringent criteria (Fig. 2E, F): First, we established the noise floor threshold using a buffer-only control sample. Second, we identified singlet synaptosomes based on their characteristic vertical distribution in the scatter plot, consistent with previous literature reports^38,41^. Third, we selected membrane-intact particles by gating on the top ∼90% of CellBrite 405 signal intensity to exclude dimly stained outliers. Finally, we identified double-positive (V5+HA+) synaptosomes using gates calibrated against wildtype control samples stained in parallel - specifically, we set thresholds that yielded less than 0.5% false positives in the V5 channel and less than 1% false positives in the HA channel. Confocal imaging of sorted positive populations verified their singlet morphology and fluorescent signals (Fig. 2G), with quantitative analysis showing ∼90% HA signal retention in HA+ sorted populations and ∼65% V5 signal retention in V5+ sorted populations (Extended Data Fig. 3D). Furthermore, analysis of double-positive structures revealed a higher proportion in V5+ versus HA+ sorted populations (Extended Data Fig. 3E), consistent with the relative frequencies observed in flow cytometry data (Fig. 2F). Using this approach, we could routinely isolate approximately 20,000 double-positive synaptosomes per hour. They were similarly sorted into a small amount of collection buffer in a pre-wet PCR strip tube and directly subjected to 10x Genomics workflow like sorted GFP+ nuclei.

Meanwhile, we also assessed the potential contamination of RNA barcodes from nuclei into synaptosome fractions during biochemical fractionation, especially for PostRNA due to the proximity of cerebellar nuclei and synapses. We designed an experiment to compare barcode contents of synaptosomes prepared from cerebellum samples injected with PostSynBar and PostRNA either together or separately (Extended Data Fig. 3F). After sorting HA+ synaptosomes and quantify PostRNA contents via qPCR, we observed significantly higher level of PostRNA in the co-injected group compared to the negative control post-injection mixed group (Extended Data Fig. 3G). These data indicate that contamination of RNA barcodes from nuclei into synaptosomes is minimal under our experimental conditions.

### 3. Development of parallel single-nucleus and single-synaptosome sequencing

To capture both cellular identity and synaptic connectivity information, we developed parallel single-nucleus and single-synaptosome sequencing. This method combines single-nucleus RNA sequencing with single-synaptosome barcode sequencing to link neuronal transcriptomes with their synaptic connections through an optimized library construction protocol. 10x Genomics Single Cell 3 protocol with featuring barcoding was modified to enable simultaneous capture of transcriptomes and barcode RNAs. Within each loaded microfluidics-generated oil emulsion droplet, single nucleus or synaptosome was encapsulated with a single gel bead containing three types of reverse transcription oligos: poly(dT) for mRNAs, and two specific capture sequences for PreRNA and PostRNA, respectively (Fig. 3A). This design enables concurrent reverse transcription of cellular transcripts and barcode RNAs within the same droplet without potential competition.

**Figure 3.**
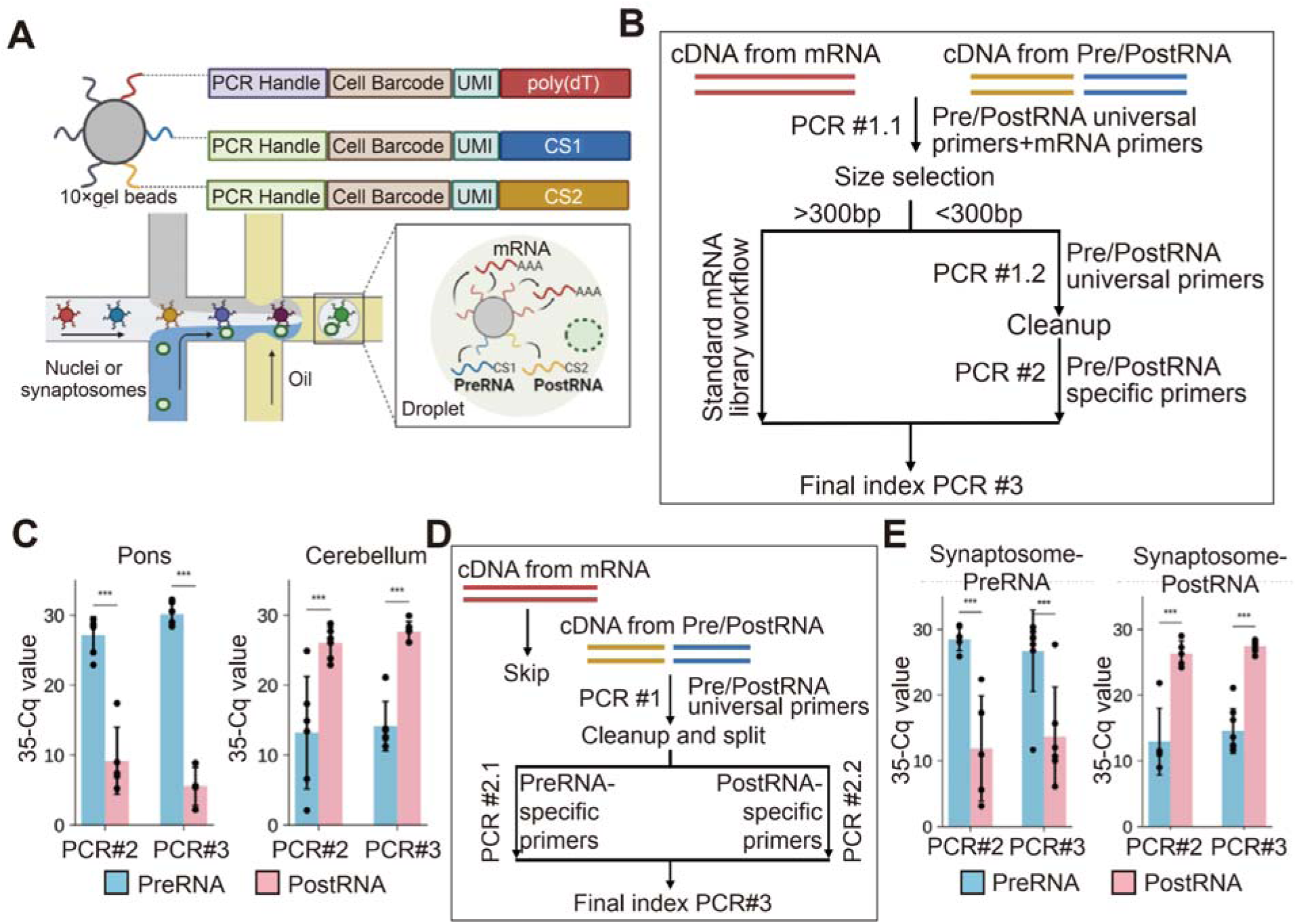
Development of parallel single-nucleus and single-synaptosome sequencing. (A) Adapting 10x Genomics gel bead design for simultaneous capture of transcriptomes and barcodes. Top: Structure of three capture oligonucleotides on gel beads - poly(dT) for mRNA (red), Capture Sequence 1 (CS1) for PreRNA (blue), and Capture Sequence 2 (CS2) for PostRNA (orange). Bottom: Schematic of microfluidic co-encapsulation showing nuclei/synaptosomes with gel beads in oil emulsion droplets. Inset shows molecular components within a droplet. (B) Nuclear library construction workflow. The protocol separates mRNA-derived cDNA (red) from Pre/PostRNA-derived cDNA (blue/orange) through size selection after initial PCR#1.1. Large fragments (>300bp) undergo standard mRNA library preparation, while small fragments (<300bp) are processed through PreRNA- or PostRNA-specific amplification steps. PCR#2 uses barcode-specific primers, followed by final indexing PCR#3. (C) qPCR analysis showing selective amplification of PreRNA and PostRNA barcodes from nuclei samples from pons and cerebellum. Bar plots show Taqman qPCR values (35-Cq) for PreRNA (blue) and PostRNA (pink) after PCR#2 and PCR#3. Higher values indicate greater abundance. PreRNA barcodes showed consistently higher abundance in pons samples whereas PostRNA barcodes showed consistently higher abundance in cerebellum samples, demonstrating specificity of the amplification strategy. Error bars represent standard deviation; individual data points shown as black dots. *** p < 0.001, paired t-test. (D) Synaptosome library construction workflow optimized for barcode recovery. mRNA capture is omitted, and both PreRNA and PostRNA cDNA undergo direct amplification with universal primers (PCR#1). The protocol then splits into separate reactions for PreRNA- and PostRNA-specific amplification (PCR#2.1 and PCR#2.2) before final indexing. (E) qPCR validation of Pre- and PostRNA barcode enrichment in synaptosome samples. Bar plots show Taqman qPCR values (35-Cq) comparing PreRNA-enriched (blue) and PostRNA-enriched (pink) samples after PCR#2 and PCR#3. Synaptosome-PreRNA samples show specific enrichment of PreRNA barcodes, while Synaptosome-PostRNA samples show specific enrichment of PostRNA barcodes, confirming successful library construction of each type of barcode. Error bars represent standard deviation; individual replicates shown as black dots. *** p < 0.001, paired t-test.

For nuclear libraries, we developed a custom library construction protocol optimized for both transcriptome and barcode capture. Our workflow involves: Initial pre-amplification (PreAmp1) targeting both mRNA and barcodes; size-based separation of barcode and mRNA libraries using optimized sample-to-purification-beads ratios; and separate amplification of transcriptome and barcode components (Fig. 3B). Taqman qPCR quantification of PreRNA and PostRNA abundance at both PCR#2 and #3 verified that expected barcode types were enriched in each nuclei sample (Fig. 3C). This final protocol achieved high-quality nuclear transcriptomes and efficient barcode co-detection, with close to 99% of nuclei containing expected barcodes (Data shown in the next section).

For synaptosome libraries, our initial attempts to capture both mRNA and barcodes were hampered by the dominance of mitochondrial RNA, which reduced barcode detection efficiency while not offering additional insights into transcriptome heterogeneity among synaptosomes (Extended Data Fig. 4A-D). We therefore chose to omit mRNA capture from synaptosomes and developed a library construction protocol for synaptosome-specific barcode recovery with five key modifications to the standard 10x Genomics workflow (Fig. 3D): First, we omitted template-switching oligonucleotides (TSO) during reverse transcription, as we already knew the complete sequences of PreRNA and PostRNA. Second, we used barcode-specific primers with short extension times during pre-amplification (PCR#1) to selectively amplify short barcodes rather than long mRNAs. Third, we separated PreRNA and PostRNA into two reactions during second amplification (PCR#2) using specific primer pairs for each. Fourth, we optimized all bead purification steps for the smaller size of barcode molecules. Fifth, we noted that standard primers for barcode amplification tend to misalign and favor the high abundance target when the difference in abundance was large between PreRNA and PostRNA, resulting in under-amplification of the less abundant target. Hence, we utilized locked nuclei acid (LNA) modified primers, replacing a few regular adenosines with LNA-modified ones at specific positions within the primer annealing region, which achieved more balanced amplification of both PreRNA and PostRNA (Extended Data Fig. 4E-H). Similarly, we implemented Taqman qPCR quantification of PreRNA and PostRNA abundance at each step, which verified that this final protocol achieved high specificity and minimal cross-contamination between barcode types (Fig. 3E).

For a typical parallel single-nucleus and single-synaptosome sequencing experiment, we processed approximately 20,000 to 30,000 nuclei from each brain region and around 30,000 to 140,000 double-positive synaptosomes. Our optimized protocol achieved robust detection of both cellular transcriptomes as well as nuclear and synaptic barcodes, enabling comprehensive mapping of neural circuits with cellular gene expression profile and single cell identity.

### 4. Application of Connectome-seq to the pontocerebellar circuit

To demonstrate the capability of Connectome-seq in mapping neuronal connectivity in vivo, we applied our method to map the mouse pontocerebellar circuit. This circuit was chosen for its well-characterized anatomy and the presence of diverse cell types, providing an excellent testbed for our technology.

We injected AAV vectors encoding PreSynBar and PreRNA constructs bilaterally into the pons, and PostSynBar and PostRNA constructs unilaterally into the cerebellar cortex of adult mice (Fig. 4A). The injections were performed using titers and volumes to achieve widespread expression while minimizing potential overexpression artifacts. After a two-week expression period, we observed robust and specific expression of PreSynBar (detected by anti-V5 antibody) in pontine neurons (Fig. 4B) and their axonal projections to the cerebellum (Fig. 4C), as well as strong expression of PostSynBar (detected by anti-HA antibody) in cerebellar neurons (Fig. 4D). The specificity of PreSynBar localization was evidenced by the colocalization with VGLUT1 and VGLUT2 markers expressed in the presynaptic mossy fiber terminals projected from pons (Extended Data Fig. 5A). Barcoded synapses were detected via abundant specific split GFP signals at the interface between PreSynBar- and PostSynBar-expressing neurons (Fig. 4E). RNA in situ hybridization showed colocalization of PreRNA and PostRNA with SynBar proteins at mossy fiber terminals, confirming efficient barcode transport and localization at synapses *in vivo* (Fig. 4F, Extended Data Fig. 5B-D). Notably, PreRNA showed a clustered distribution pattern consistent with its localization in presynaptic glomeruli structures, while PostRNA exhibited a more dispersed pattern throughout postsynaptic neurons with partial enrichment at PostSynBar-positive synapses.

**Figure 4.**
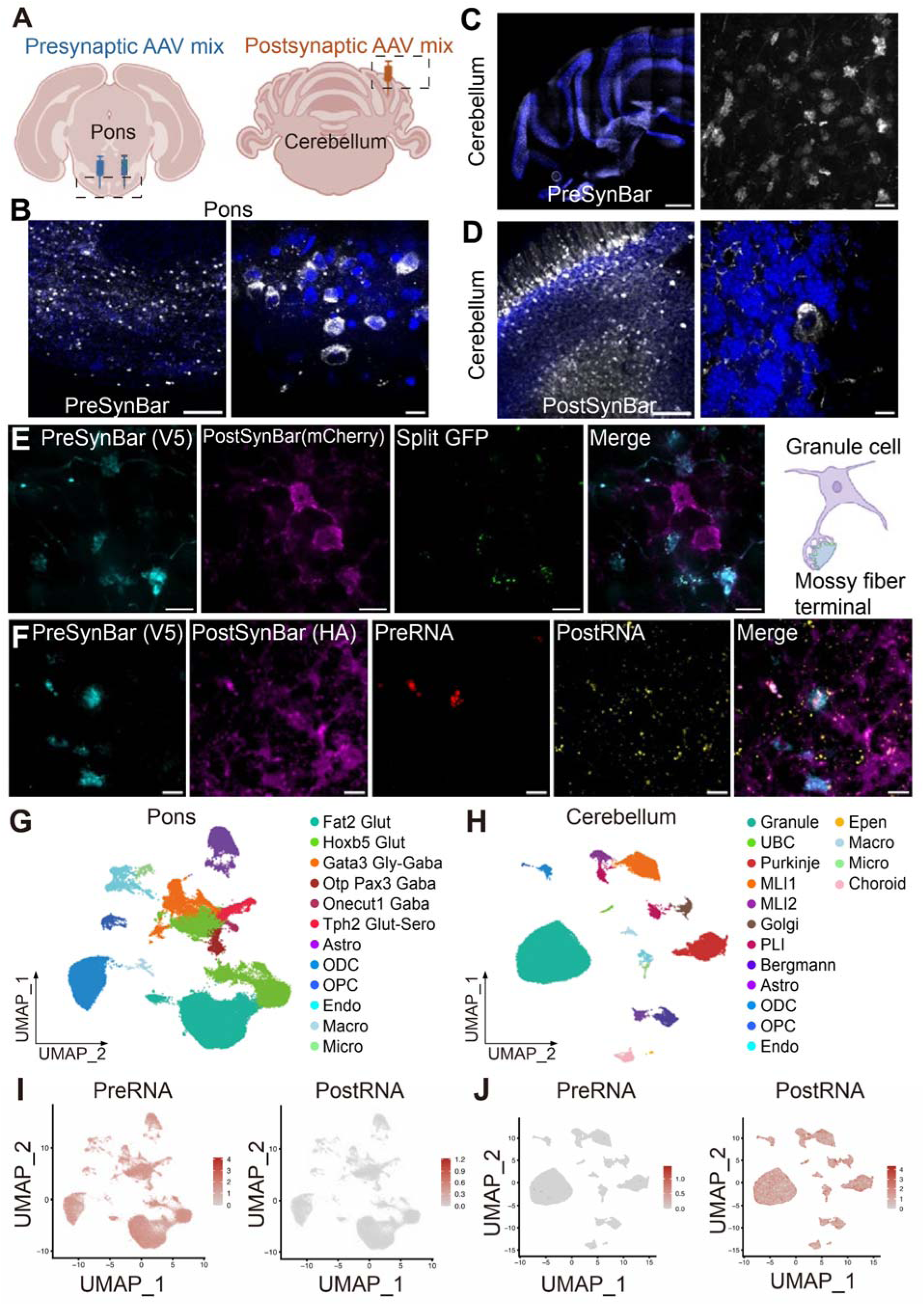
*In vivo* application and validation of Connectome-seq in the mouse pontocerebellar circuit. (A) Schematic showing bilateral injection of presynaptic AAV mix (blue) into the pons and unilateral injection of postsynaptic AAV mix (orange) into the cerebellum. (B) Confocal images of pons neurons expressing PreSynBar. Left: low magnification showing widespread expression of PreSynBar in pons neurons. Right: high magnification showing cellular expression pattern. Blue: DAPI; White: PreSynBar signal stained with anti-V5 antibody. Scale bars: left panel 100 μm, right panel 10 μm. (C) PreSynBar projection to the cerebellum. Left: low magnification showing cerebellar lobules with PreSynBar-positive mossy fiber terminals from pons (white, anti-V5 antibody staining) in the granule cell layer outlined by strong DAPI signal (blue). Right: high magnification showing individual mossy fiber terminals expressing PreSynBar. Scale bar: left panel 100 μm, right panel 10 μm. (D) PostSynBar expression across different cerebellar cell types. Left: low magnification view showing distinct PostSynBar expression patterns in molecular layer interneurons, Purkinje cells, and granule cells based on their characteristic locations relative to the granule cell layer. Right: high magnification showing cellular morphology details with dense granule cell dendrites and one Purkinje cell soma. Blue: DAPI; White: PostSynBar signal stained with anti-HA antibody. Scale bar: left panel 100 μm, right panel 10 μm. (E) Split-GFP reconstitution at synaptic contacts. From left to right: PreSynBar-expressing mossy fiber terminals (cyan), PostSynBar-expressing granule cells (magenta), split GFP signals marking connected synapses (green), and merged image showing colocalization. Scale bar: 10 μm. (F) RNA fluorescence in situ hybridization (FISH) validating synaptic barcode localization in the cerebellum. From left to right: PreSynBar protein (cyan), PostSynBar protein (magenta), PreRNA RNA (red), PostRNA RNA (yellow), and merged image showing colocalization. Scale bar: 10 μm. (G) UMAP visualization of major cell populations in the pons, showing distinct clusters of neuronal populations with different neurotransmitter expression, including glutamatergic (Fat2 Glut and Hoxb5 Glut), GABAergic (Gata3 Gly-Gaba, Onecut1 Gaba, Otp Pax3 Gaba), serotonergic (Tph2 Glut-Sero), as well as various glial cell types including astrocytes (Astro), oligodendrocytes (ODC), oligodendrocyte precursor cells (OPC), endothelial cells (Endo), macrophage (Macro), and microglia (Micro). (H) UMAP visualization of major cerebellar cell populations, highlighting granule cells, Purkinje cells, Golgi cells, unipolar brush cells (UBC), molecular layer interneurons (MLI1 and MLI2), Purkinje cell layer interneurons (PLI), and various non-neuronal cell types including Bergmann glia, astrocytes (Astro), oligodendrocytes (ODC), oligodendrocyte precursor cells (OPC), endothelial cells (Endo), ependymal cell (Epen), macrophage (Macro), microglia (Micro), and choroid plexus cells (Choroid). (I) UMAP visualization showing PreRNA expression distribution across all pontine cell populations. PostRNA has virtually no expression as expected. Scale bars indicate centered log-ratio (CLR)-normalized relative barcode expression levels (per cell). (J) UMAP visualization showing PostRNA expression distribution across all cerebellar cell populations. PreRNA has virtually no expression as expected. Scale bars indicate CLR-normalized relative barcode expression levels (per cell).

To capture the full diversity of cell types and connectivity in both regions while minimizing individual differences, we performed Connectome-seq on six biological replicates. For each replicate, we isolated and sequenced all GFP-positive singlet nuclei from both infected regions in pons and cerebellum, and tens of thousands synaptosomes from the same cerebellar region. For each sequencing experiment, we aimed to sort and sequence all GFP-positive nuclei from the dissected brain region to ensure capturing the nuclear barcode pool as completely as possible. For synaptosomes, we stained approximately one-third of the total synaptosomes isolated from each cerebellum. Under our sorting conditions, it took about 1.5 hours to collect sufficient double-positive synaptosomes (∼30,000) for 10X sequencing, which typically consumed one-third of the stained sample - effectively sampling about 10% of the total synaptosome preparation. While our flow cytometry does not intentionally select synaptosomes based on size or fluorescence intensity, the overall approach may still be subject to technical biases, including differential AAV transduction efficiency across cell types, varying nuclear RNA recovery rates that depend on neuronal size, and size- or shape-dependent differences in synaptosome purification yields. These potential sources of bias should be considered when interpreting the observed distribution of synapse types.

Single-nucleus RNA sequencing of the sorted GFP-positive nuclei revealed a diverse array of cell types in both the pons and cerebellum, with their proportions closely matching the expected distributions based on previous studies. In total, we recovered 180,468 and 129,192 nuclei profiles from pons and cerebellum with a median transcript capture of 1,852 and 1,346 unique molecular identifiers per profile respectively across all six individual brain samples. Using established cell type markers from published dataset, we performed comprehensive cell typing in each brain region using gene expression data from snRNA-seq.

Through unbiased clustering analysis (Extended Data Fig. 6A, B) followed by quality control to remove low-quality clusters (Extended Data Fig. 6C, D), we identified several distinct neuronal and non-neuronal populations in the pons and cerebellum. Initial cell type annotation was performed using canonical marker genes (Extended Data Fig. 6E, F) and further refined using the Allen Institute’s MapMyCell annotation tool^42^ (Extended Data Fig. 6G, H). Following quality control, we identified 109,269 and 78,358 high-quality nuclei profiles from pons and cerebellum (Fig. 5G, H), with cell type identities validated using marker gene expression across merged clusters (Extended Data Fig. 6I, J). In the pons, neuronal populations comprised approximately 77% of total cells, with glutamatergic neurons being the most abundant (73,483 cells, including 36,212 Fat2 Glut and 37,271 Hoxb5 Glut populations), followed by GABAergic neurons (8,389 cells, including 1,673 Otp Pax3 Gaba, 6,098 Gata3 Gly-Gaba, and 618 Onecut1 Gaba populations) and serotonergic neurons (2,318 cells). These populations were classified based on their molecular markers, including *Slc17a6* (VGLUT2) and *Slc17a7*(VGLUT1) for glutamatergic neurons, *Gad1*, *Gad2*, and *Slc32a1* (Vgat) for GABAergic neurons, and *Slc6a4* and *Tph2* for serotonergic neurons. While our injection targeted pontine nuclei which are predominantly glutamatergic, some viruses may inherently spread to adjacent brainstem structures. The GABAergic neurons detected likely originate from neighboring regions such as the caudal pontine reticular nucleus, where GABAergic neurons are well-characterized and involved in startle responses and anxiety like behavior^43,44^.The non-neuronal cells included oligodendrocytes (ODC, 12,739 cells), astrocytes (6,353 cells), oligodendrocyte precursor cells (OPC, 1,561 cells), and smaller populations of endothelial cells (3,444 cells), microglia (519 cells), and macrophages (463 cells). In the cerebellum, neurons constituted approximately 91% of all cells, with granule cells being the predominant population (49,385 cells, ∼63% of total cells). The remaining neuronal populations included molecular layer interneurons (MLI1:7,190 cells; MLI2: 1,610 cells), Purkinje cells (9,163 cells), Purkinje layer interneurons (PLI, 2,085 cells), Golgi cells (1,423 cells), and unipolar brush cells (UBC, 298 cells). Among non-neuronal cells, we identified Bergmann glia (2,023 cells), oligodendrocytes (897 cells), astrocytes (1,042 cells), ependymal cells (91 cells), choroid plexus cells (1,543 cells), and smaller populations of endothelial cells (584 cells), microglia (398 cells), and macrophages (453 cells). Cell type-specific marker genes were selected based on previously published cerebellar atlas data^45^. Notably, granule cells, which typically constitute ∼99% of cerebellar neurons, represent only 63% in our dataset—an underrepresentation also observed in other single-cell studies^42,45^. This widespread technical limitation likely reflects granule cells’ small size and low RNA content, making them particularly susceptible to exclusion during quality filtering, as evidenced by the substantial reduction in cerebellar nuclei counts after quality control (from 129,192 to 78,358).

**Figure 5.**
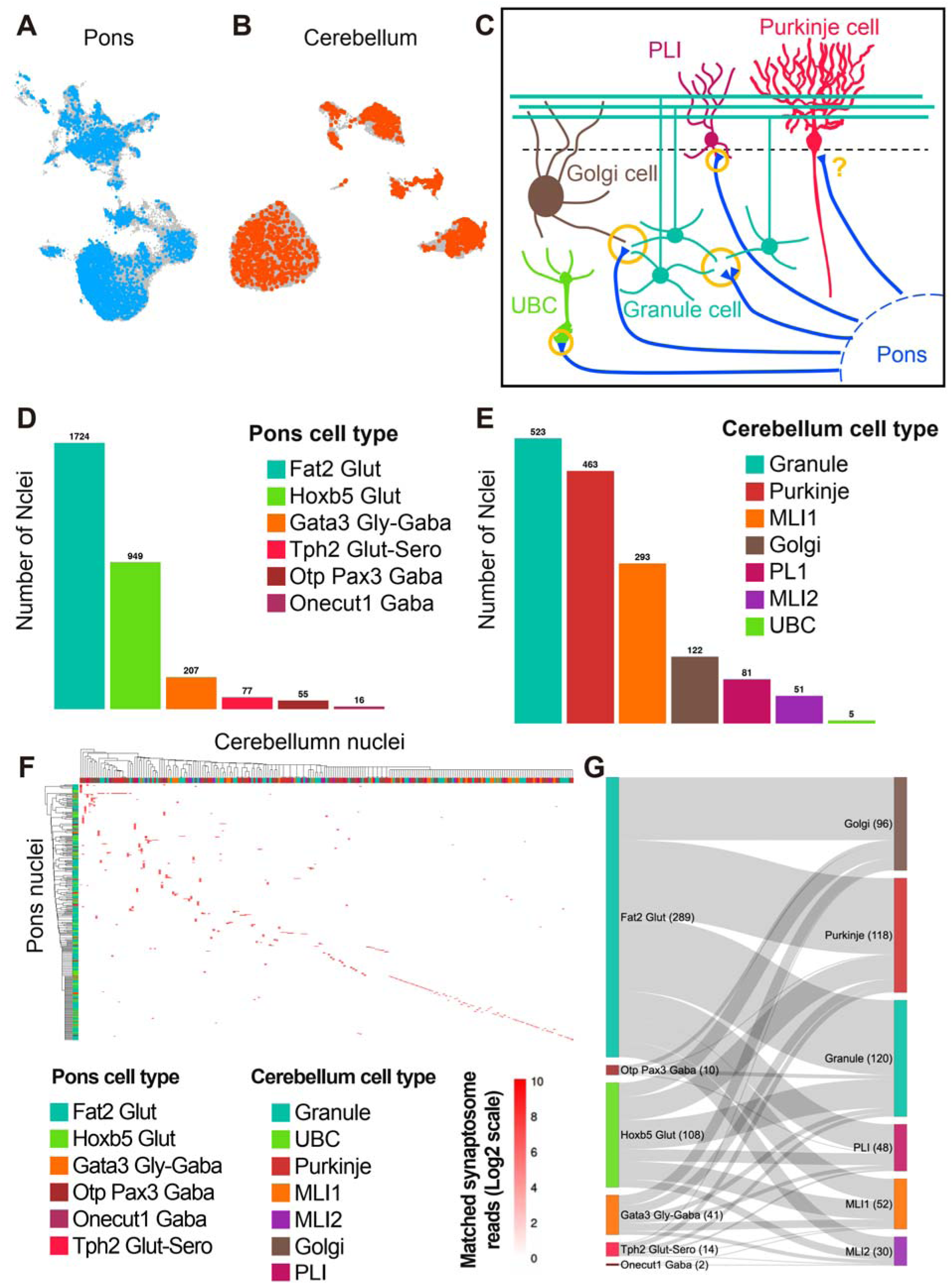
Single-cell resolution mapping of the pontocerebellar circuit. (A) UMAP visualization of pons nuclei colored by matched status (blue = synaptosomally matched nuclei, gray = unmatched nuclei). (B) UMAP visualization of cerebellar nuclei colored by matched status (orange = synaptosomally matched nuclei, gray = unmatched nuclei). (C) Schematic illustration of the canonical pontocerebellar circuit architecture. Major cell types are shown with their characteristic morphologies and relative positions. Major cellular targets of pontine axons (blue lines) include granule cells (teal) and Golgi cells (brown), as well as other targets including UBCs (unipolar brush cells, green) and PLI (Purkinje layer interneurons, magenta). Purkinje cell (red) is shown with its characteristic dendritic arbors, while its status as a direct pontine target is under investigation. (D) Quantification of matched cell type distribution in the pons (n=3,028 nuclei). Cell types include Fat2 Glut (n=1,724), Hoxb5 Glut (n=949), Gata3 Gly-GABA (n=207), Tph2 Glut-Sero (n=77), Otp Pax3 Gaba (n=55), and Onecut1 Gaba (n=16). (E) Quantification of matched cell type distribution in the cerebellum (n=1,538 nuclei). Distribution shows Granule cells (n=523), Purkinje cells (n=463), MLI1 (n=293), Golgi cells (n=122), PLI (n=81), MLI2 (n=51), and UBC (n=5). (F) Cell type-specific connectivity matrix between pons neurons (rows, n=327) and cerebellar neurons (columns, n=219). Color intensity indicates synaptosome Pre/PostRNA UMI counts (Log2 scale) supporting each neuron pair connection. Hierarchical clustering reveals organized patterns of connectivity. (G) Sankey diagram quantifying synaptosomal connections between major cell types. Width of flows represents the number of unique synaptosomes between each cell type pair. Numbers indicate total unique matched synaptosomes detected for each cell type: Fat2 Glut (289), Hoxb5 Glut (108), Gata3 Gly-GABA (41), Tph2 Glut-Sero (14), Otp Pax3 Gaba (2), and Onecut1 Gaba (2) from pons connecting to Golgi (96), Purkinje (118), Granule (120), PLI (48), MLI1 (52), and MLI2 (30) in cerebellum.

Importantly, we detected barcodes in a high percentage of the sequenced nuclei, with 98.6% of pontine nuclei containing PreRNA and 98.7% of cerebellar nuclei containing PostRNA (Fig. 4I, J). This high efficiency of mRNA and barcode co-detection was crucial for comprehensive connectivity mapping, as they serve as the source pool of all synaptic barcodes to match with. Overall, these results confirmed that our modified library construction strategy to allow both mRNA and barcode capture generated sufficient data to accurately determine barcoded cell types, and that our viral strategy recapitulated the natural distribution of different cell types in each targeted region as reported by other unbiased single cell sequencing experiments.

To assess the diversity of our captured barcode pool, we first analyzed total barcode diversity from sequencing libraries generated from AAV genomes (Extended Data Fig. 7A). We detected ∼29 million distinct PreRNA barcodes from ∼100 million reads (two libraries of ∼50 million each) and ∼17 million distinct PostRNA barcodes from ∼50 million reads. At injected AAV amount typical for our experiments (5 billion particles in pons, 1.4 billion in cerebellum), the projected diversity reached ∼61 million distinct sequences for PreRNA and ∼55 million for PostRNA. Comparing AAV, nucleus, and synaptosome libraries revealed a progressive decline in preserved diversity: nucleus libraries retained 15.3% of PreRNA sequences (4.4 million) and 19.4% of PostRNA sequences (3.3 million), while synaptosome libraries retained only 1.7% of PreRNA sequences (0.5 million) and 7.6% of PostRNA sequences (1.3 million). Notably, in synaptosomes the relative order of PreRNA and PostRNA diversity was reversed (virus and nucleus libraries total distinct barcode count: PreRNA > PostRNA; synaptosomes library total distinct barcode count: PreRNA < PostRNA), which may reflect the more demanding axonal transport of PreRNA compared to dendritic transport of PostRNA. Distribution analysis showed reduced overall diversity but higher maximum counts and heavier tails from virus to nucleus to synaptosome libraries (Extended Data Fig. 7B), likely caused by increasingly more PCR cycles needed during library preparation. Per-neuron and per-synaptosome comparisons confirmed the stronger reduction of barcode diversity in synaptosomes (Extended Data Fig. 7C). Joint density analysis indicated that most synaptosomes contained only modest numbers of PreRNA and PostRNA UMIs (Extended Data Fig. 7D), and read depth analysis showed most UMIs were supported by just 1–10 reads (Extended Data Fig. 7E–F). These consistent patterns across PreRNA and PostRNA suggest technical variation, rather than compartment-specific bias, dominated read depth. Overall, the presence of extremely high-count barcodes and unusually barcode-rich nuclei/synaptosomes suggest issues in PCR bias and ambient RNA contamination, underscoring the importance of the stringent quality-control and filtering steps we applied throughout our analysis.

### 5. Reconstruction of the pontocerebellar connectome

With the efficient barcoding of nuclei and synaptosomes in the pontocerebellar circuit using Connectome-seq constructs, we next focused on reconstructing the connectome from our sequencing data. We developed a computational pipeline to extract, error-correct, and match barcodes from both nuclear and synaptosomal sequencing data (Extended Data Fig. 8A). The pipeline implements multi-level error correction, including CB-UMI–based filtering and network-based 30-mer correction, followed by stringent barcode matching criteria. We first collapsed barcodes from each dataset by allowing up to 1 mismatch among all detected unique N30 sequences. Next, we performed inter-sample matching by allowing up to 5 mismatches between the barcodes from nuclei and synaptosomes to account for PCR and sequencing introduced errors.

To determine this 5-mismatch threshold as appropriate, we compared the pairwise Hamming distance distribution of our experimental barcode libraries against the theoretical distribution expected from our random 30-mer barcode design (Extended Data Fig. 8B). Virus libraries closely aligned with the theoretical curve beginning at Hamming distance 2, while nucleus libraries only began to align after 6 mismatches, indicating that errors introduced during library preparation caused nucleus barcodes to deviate from the original design. We selected 5 mismatches as our threshold because this captures most mutation-derived matches while maintaining reasonable false positive control. To further minimize false positives, we implemented a biological constraint that no single synaptosome can originate from multiple neurons—synaptosomes mapping to multiple nuclei within the maximum allowed Hamming distance were excluded from analysis. This combination of mismatch tolerance and biological constraint provided optimal balance between sensitivity and specificity.

Next, we collapsed barcodes within each dataset by testing 0–10 mismatches across all unique N30 sequences with matching rates steadily increasing and plateauing at Hamming distance 5 (Extended Data Fig. 8C). Our analysis revealed high barcode diversity in the synaptosomal fraction, with 81,452 unique PreRNA and 147,164 unique PostRNA sequences detected from ∼600k barcode-containing synaptosomes (Extended Data Fig. 8D). At the 5-mismatch threshold, 32,269 (39.6%) PreRNA were uniquely matched to their source in the presynaptic nuclear pool, while 62,205 (42.3%) PostRNA were uniquely matched to the postsynaptic nuclear source. Despite technical factors affecting barcode recovery and matching rates, the matched barcodes provided sufficient coverage to reconstruct major circuit features, as demonstrated by the cell-type specific connectivity patterns described below.

When analyzing PreRNA and PostRNA matches separately, we found that 3,028 pons nuclei and 1,538 cerebellar nuclei were matched with corresponding synaptosomes (Fig. 5A, B). To assess the reliability of these matches, we first evaluated their consistency with known pontocerebellar circuit architecture (Fig. 5C). The canonical circuit consists of mossy fibers from pontine neurons that project through the middle cerebellar peduncle to target three main cell types in the cerebellar cortex: granule cells, Golgi cells, and unipolar brush cells (UBC)^46^. Recent studies have revealed additional connectivity patterns, such as mossy fiber inputs onto candelabrum cells, a type of Purkinje cell layer interneurons (PLI)^47^. Transient mossy fiber inputs to Purkinje cells during development were also long known to exist^48^. Given that our molecular approach has the potential to uncover novel connectivity patterns, where unexpected connections may represent genuine biological findings rather than technical artifacts, we focused our false discovery rate (FDR) analysis on the pontine side, where glutamatergic neurons are well-established as the classical projection neurons. When excluding all GABAergic neurons, the FDR was 9.2% (278/3,028). Using more stringent criteria where only glutamatergic neurons (Fat2 Glut and Hoxb5 Glut) are considered true positives (excluding both GABAergic and serotonergic neurons), the estimated FDR increases to 11.7% (355/3,028). It is worth noting that serotonergic neurons in the pons, particularly from raphe nuclei, may also project to the cerebellum^49^, though definitive molecular evidence for this connection remains to be established.

We next analyzed two key metrics of the connected populations: (1) the relative abundance of each cell type within the matched population (percentage of matched cells) and (2) the connection rate for each cell type (percentage of total cells of that type that were matched to synaptosomes. In the pons (Fig. 5D), among the 3,028 matched neurons, glutamatergic neurons were the predominant population, with Fat2 Glut (1,724 cells, 56.9% of matched; 4.76% match rate) and Hoxb5 Glut cells (949 cells, 31.3% of matched; 2.55% match rate) together accounting for 88.2% of all matched neurons. The remaining populations included Gata3 Gly-GABA (207 cells, 6.8% of matched; 3.39% match rate), Tph2 Glut-Sero (77 cells, 2.5% of matched; 3.32% match rate), Otp Pax3 Gaba (55 cells, 1.8% of matched; 3.29% match rate), and Onecut1 Gaba (16 cells, 0.53% of matched; 2.59% match rate). The high proportion and match rates of glutamatergic neurons align with their established role as the primary projection neurons in pontocerebellar circuits. In the cerebellum (Fig. 5E), the 1,538 matched neurons showed a diverse cellular composition reflecting the layered architecture of cerebellar cortex. Granule cells, despite their small size, formed the largest population (523 cells, 34.0% of matched; 1.06% match rate), followed by Purkinje cells (463 cells, 30.1% of matched; 5.05% match rate), MLI1 interneurons (293 cells, 19.1% of matched; 4.08% match rate), Golgi cells (122 cells, 7.93% of matched; 8.57% match rate), PLI (81 cells, 5.3% of matched; 3.88% match rate), MLI2 (51 cells, 3.3% of matched; 3.17% match rate), and UBC (5 cells, 0.3% of matched; 1.68% match rate). Total UMI counts of matched barcodes for each cell types also closely resembled the distribution of cell numbers (Extended Data Fig. 8E, F). The varying match rates likely reflect a combination of biological and technical factors, with the notably low match rate for granule cells potentially attributed to their small soma size (∼5-8 μm) compared to Purkinje cells (∼20-40 μm) and Golgi cells (∼20 μm), which would affect both viral infection efficiency and RNA capture in single-nucleus sequencing.

Same analysis of matched barcodes from synaptosomes showed similar distribution patterns matching the nuclei ones for both synaptosomes counts and barcode UMI counts (Extended Data Fig. 8G–J). We also quantified the average number of synaptosomes associated with each neuron organized by cell types (Extended Data Fig. 8K, H). In the pons, different cell types have similar average matched synaptosome counts per cell, whereas in the cerebellum, Golgi cells displayed a much higher average (14.3) count compared to granule cells (6.4), which is consistent with known differences in their synaptic connectivity (one Golgi cell contacting much more mossy fiber terminals than a granule cell).

To reconstruct the full connectome, we analyzed synaptosomes containing both PreRNA and PostRNA that matched with our filtered nuclear populations, allowing us to map connections between individual pons and cerebellar neurons. The resulting clustered connectivity matrix (Fig. 5F) revealed distinct connectivity patterns at single cell level organized by cell types, where 327 pons neurons formed connections with 219 cerebellar neurons. Each red box in the matrix represents synaptosome Pre/PostRNA UMI counts (Log2 scale) supporting specific neuron pairs, with a subset of (∼15.7%) neuron pairs supported by over 3 synaptosome barcode UMI count, suggesting potentially stronger connections. The hierarchical clustering reveals modules of connectivity dominated by glutamatergic inputs and more distributed outputs. Quantification of these connections by cell type (Fig. 5G) shows that among the 464 total unique synaptosomes, pontine glutamatergic neurons (Fat2 and Hoxb5 Glut) formed the majority (289 and 108 unique synaptosomes respectively). The remaining populations - Gata3 Gly-GABA, Tph2 Glut-Sero, Otp Pax3 Gaba, and Onecut1 Gaba - showed more modest connectivity (41, 14, 10, and 2 unique synaptosomes respectively). On the cerebellar side, granule cells received the highest number of unique synaptosomal connections (120), followed by Purkinje cells (118), Golgi (96), MLI1 (52), PLI (48), and MLI2 (30). The substantial numbers of Glut-to-granule cell and Glut-to-Golgi cell connections align with known feedforward excitatory and inhibitory pontocerebellar circuitry, while the similar number of Glut-to-Purkinje cell connections represents an unexpected finding that warrants further investigation. The organized, cell-type specific patterns observed in both the matrix and the quantified connections suggest these represent genuine circuit features rather than random noise, though some connections may still reflect technical artifacts given current understanding of pontocerebellar circuits.

### 6. Analysis and validation of a novel pons-to-Purkinje cell connectivity

Having established the technical framework of Connectome-seq and verified its ability to capture known connectivity patterns, we wanted to evaluate its potential in revealing new connections in well-established neural circuits. We decided to follow up on our intriguing discovery of the consistent connection between pontine neurons and Purkinje cells. The existence of such connectivity has been debated in the field, with previous studies only reporting its presence in the developing brain, not in adults^48,50^.

To investigate the molecular characteristics of these connected neurons, we isolated all the Purkinje cells from our cerebellar snRNA-seq data and further grouped them into seven subclusters (Fig. 6A). Intriguingly, Purkinje cells with pons connection identified by Connectome-seq (Cseq-positive) showed distinct clustering patterns, concentrating in subclusters 1, 2, 3, 4, 6, and 7, suggesting potential molecular programs associated with their connectivity (Fig. 6B). Quantification revealed the highest enrichment of connected cells in cluster 7 (42.3%), followed by cluster 6 (11.3%), with decreasing proportions in other clusters (Extended Data Fig. 9A). To identify connectivity-associated marker genes, we performed differential gene expression analysis between pons-connection Positive-cluster (1, 2, 3, 4, 6, 7) and Negative-cluster (0, 5) Purkinje cells. Our analysis revealed several highly enriched genes in connected neurons, including *Grid2ip*, *Cacna1g*, *Stac*, *Dlgap4*, *Dagla*, *Foxp4*, *Grik4, Zfp385c*, *Mir9-3hg*, and *Abr* (Fig. 6C, Extended Data Fig. 9B-E). The enriched genes fell into distinct functional categories: synaptic organization and plasticity (*Grid2ip*), ion channel / neuronal excitability (*Cacna1g*), vesicle trafficking and calcium channel modulation (*Stac*), synaptic scaffolding / postsynaptic density (*Dlgap4*), endocannabinoid signaling / lipid metabolism (*Dagla*), transcriptional regulation / neurodevelopment (*Foxp4*), glutamatergic synaptic transmission / excitatory neurotransmission (*Grik4*), RNA-binding / neuronal differentiation (*Zfp385c*), non-coding RNA / neurogenesis regulation (*Mir9-3hg*), and signal transduction / Rho GTPase regulation (*Abr*). Interestingly, *Grid2ip*/Delphilin, acts as a synaptic structural scaffold in Purkinje cells, maintaining proper synaptic connectivity and signal transmission strength. Delphilin colocalizes and interacts with the glutamate receptor δ2 subunit (GluRδ2) at synaptic sites, and is almost exclusively found at parallel-fiber–Purkinje-cell (PF-PC) synapses in contrast to climbing fiber (CF-PC) synapses in the molecular layer^51,52^. The co-enrichment of established connectivity regulators like *Grid2ip* alongside genes involved in diverse cellular processes suggests a complex molecular program integrating multiple pathways.

**Figure 6.**
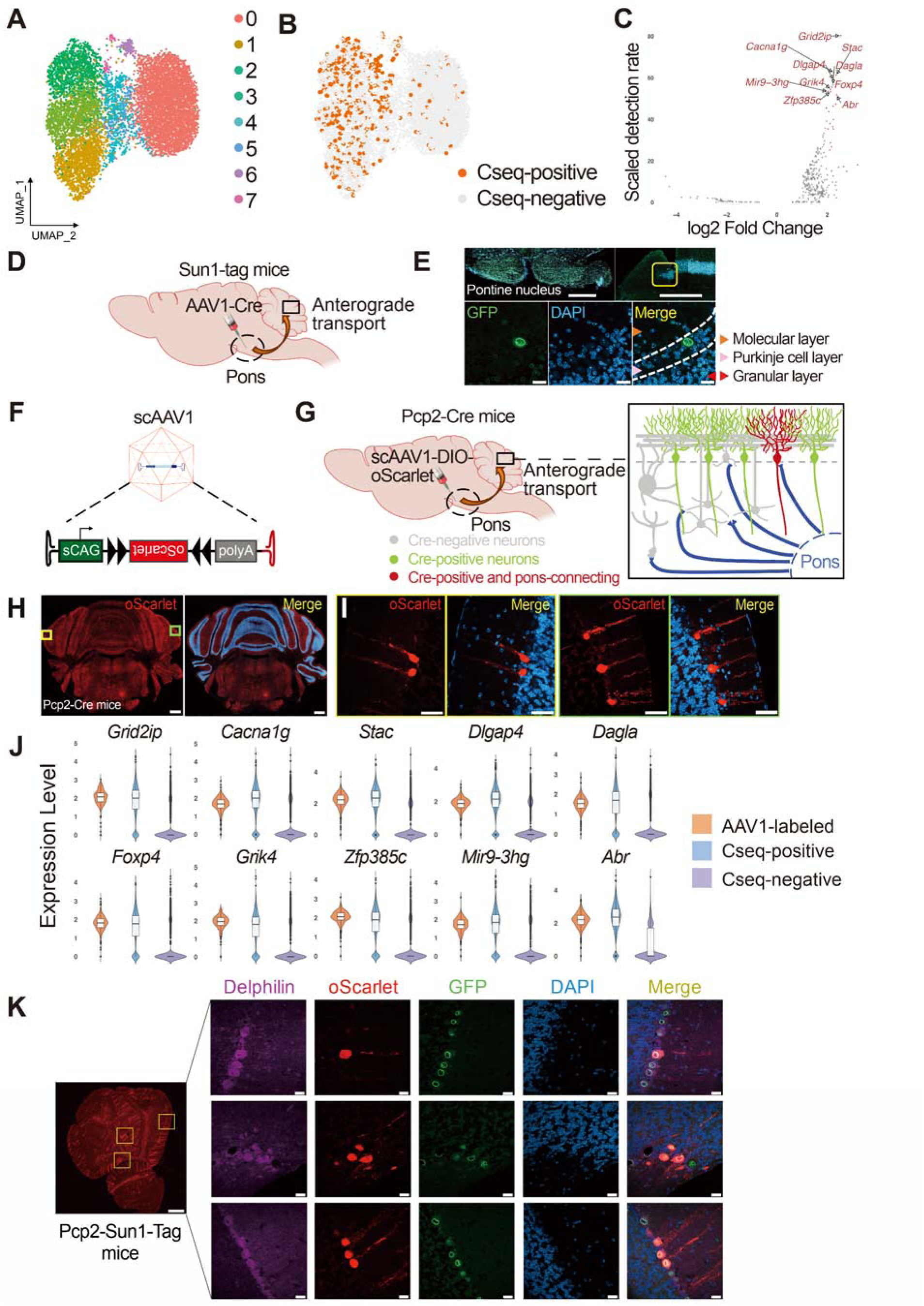
Discovery and validation of direct pons-to-Purkinje cell connectivity. (A) UMAP visualization of single-nucleus RNA sequencing data from cerebellar Purkinje cells, revealing 7 distinct molecular populations (clusters 0-7) based on transcriptional profiles. (B) UMAP plot of Purkinje cells identified through Connectome-seq as receiving direct pontine inputs (Cseq-positive, orange) or lacking pontine connections (Cseq-negative, gray), demonstrating non-random distribution of pontine-connected cells across molecular subtypes. (C) Volcano plot showing differential gene expression analysis between pons-connected and non-connected Purkinje cell clusters. Genes with an adjusted p-value corresponding to a -log10(adjusted p-value)>300 are shown. The x-axis represents the log2(fold change), and the y-axis represents the marker gene scaled detection rate (y=exp[10×(pct.1−pct.2)]/10, where pct.1 and pct.2 are the percentages of cells expressing the marker gene in positive clusters and negative clusters, respectively), emphasizing genes with large shifts in expression prevalence. Highlighted in red are marker genes with the highest detection differences, including *Grid2ip*, *Cacna1g*, *Stac*, *Dlgap4*, *Dagla*, *Foxp4*, *Grik4*, *Zfp385c*, *Mir9-3hg* and *Abr*. (D) Injection scheme of AAV1-Cre into pons of Sun1-Tag mouse and expected distribution of GFP-positive neurons. (E) Representative images of AAV1-labeled pontine nuclei (top left) and cerebellum (top right). Bottom panel reveals pons-connecting neurons including multiple granule cells, one Purkinje cell, and several other interneurons. GFP-expression (green), DAPI nuclear staining (blue), and merged images showing colocalization in reference to the anatomical layers indicated by arrows. Scale bar: top: 400 μm, bottom: 40 μm. (F) Design of self-complementary AAV1 (scAAV1) construct for anterograde trans-synaptic tracing. Vector contains a self-complementary AAV genome with short CAG promoter driving Cre-dependent oScarlet expression, flanked by inverted terminal repeats (ITRs). (G) Injection scheme of scAAV1-DIO-oScarlet into pons of Pcp2-Cre mouse and expected distribution of RFP-positive neurons. (H) Anterograde trans-synaptic tracing confirms direct pontine inputs to adult Purkinje cells. Following injection of Cre-dependent AAV1-oScarlet virus into the pons of adult Pcp2-Cre mice, labeled Purkinje cells (red) are visible across cerebellar lobules, validating persistent pons-to-Purkinje connectivity. Nuclei counterstained with DAPI (blue). Scale bar: 600 μm. (I) Representative high-magnification images of AAV1-labeled Purkinje cells. Left: oScarlet fluorescence (red) reveals detailed cellular morphology of pons-connected Purkinje cells. Right: Merged images with DAPI (blue) show characteristic positioning of Purkinje cell bodies and their elaborate dendritic trees extending into the molecular layer. Scale bar: 40 μm. (J) Expression levels of top marker genes in three different groups of Purkinje cells: AAV1-labeled (orange), Positive-cluster (blue), and Negative-cluster (purple), shown as violin plots with boxplots overlay. (K) Immunofluorescence validation of marker gene *Grid2ip*/Delphilin expression. Left: Overview of cerebellar parasagittal sections showing oScarlet-labeled pontine-connected Purkinje cells (red). Yellow boxes indicate regions magnified in right panels. Right panels (left to right): Delphilin immunostaining (magenta), pontine-connected Purkinje cells (oScarlet, red), GFP-labeled Purkinje cell signal (green), DAPI nuclear staining (blue), and merged image showing colocalization. Three representative cerebellar lobules are shown. Scale bars: 400 µm for left panel, 20 µm for right panel.

To validate this unexpected finding, we performed anterograde tracing with AAV1, which spreads trans-synaptically through a VAMP2-dependent synaptic vesicle release mechanism^53^. We chose this approach over conventional retrograde tracing because it allows selective visualization of postsynaptic targets without the confounding effects of axon collaterals or non-specific uptake that can occur with traditional tracers. We first injected AAV1-Cre virus into the pons of Sun1-Tag mice (Cre-dependent nuclear GFP), and observed robust GFP expression in the pons and cerebellum neurons including granule cells, Purkinje cells, and other interneurons (identified based on size and anatomical localization), which was consistent with our sequencing results (Fig. 6D, E). To further validate the connectivity of Purkinje cells, we developed a Cre-dependent AAV1-DIO-oScarlet virus (Fig. 6F), incorporating both a self-complementary AAV genome and a short CAG promoter to achieve high expression levels of oScarlet in the postsynaptic neurons driven by Cre. This design was based on previous work demonstrating that these two features enhanced protein expression from trans-synaptically trafficked viral particles^54^. We injected this Cre-dependent virus into the pons of adult Pcp2-Cre mice, which express Cre specifically in Purkinje cells (Fig. 6G). This approach revealed robust oScarlet expression in Purkinje cells, providing strong evidence for persistent pons-to-Purkinje cell connections in the adult brain (Fig. 6H, I). To comprehensively analyze the spatial distribution of pons-connecting Purkinje cells, we crossed Pcp2-Cre with Sun1-Tag reporter line to generate Pcp2-Sun1-Tag mice that express nuclear GFP in all Purkinje cell nuclei, and injected AAV1-DIO-oScarlet virus into the pons of these mice. Detailed imaging analysis across multiple parasagittal sections revealed a clear spatial pattern of pontine connectivity (Extended Data Fig. 10A). Quantification of the tracing ratio (oScarlet-positive/total GFP-positive Purkinje cells) showed that the fraction of pons-connected Purkinje cells decreased systematically from lateral to medial regions, reaching minimum levels in medial vermal areas (Extended Data Fig. 10B). This sagittal analysis confirmed the pattern observed in coronal sections (Fig. 6H), which showed high tracing ratios in lateral regions with sharp drops in the central region. The convergence of these two independent planes of analysis demonstrates a pronounced lateral-to-medial gradient where pontocerebellar projections preferentially target lateral hemispheres and posterior vermis (lobules VI-VIII), while showing minimal connections to anterior vermis (lobules I-V), consistent with established patterns of pontine input to the cerebellum^55^

To provide an independent validation of our connectivity-marker genes, we performed single-nucleus sequencing of Purkinje cells labeled by nuclear GFP from our AAV1-Cre injection experiment, and performed differential gene expression analysis comparing them with Positive-cluster and Negative-cluster Purkinje cells from our original Connectome-seq dataset. The result confirmed that all top 10 connectivity markers were highly enriched in AAV1-labeled Purkinje cells (Fig. 6J). Based on these findings, we selected five genes (*Grid2ip*/Delphilin, CACNA1G, STAC, DLGAP4, and ABR) for protein-level validation by immunofluorescence on cerebellar sections from the same AAV1-oScarlet-injected Pcp2-Sun1-Tag mice. All five proteins were expressed in a subset of Purkinje cells, and importantly, all oScarlet-positive Purkinje cells expressed these markers, confirming our transcriptomic findings at the protein level (Extended Data Fig. 10C). We then focused on *Grid2ip*/Delphilin, which showed the strongest enrichment, for detailed parasagittal imaging analysis. This revealed a similar lateral-to-medial gradient in Delphilin expression that mirrored pons-to-PC connectivity (Extended Data Fig. 10D). High-magnification imaging of representative parasagittal slices demonstrated robust Delphilin expression in pons-connected Purkinje cells (Fig. 6K and Extended Data Fig. 10E). RNA in situ hybridization data from the Allen Brain Atlas database further confirmed the presence of *Grid2ip* transcript in a subset of Purkinje cells, supporting *Grid2ip* as a potential molecular predictor of pons connectivity (Extended Data Fig. 10F). Taken together, these results demonstrated the ability of Connectome-seq to reveal novel circuit features and relevant molecular markers while highlighting the importance of molecular context in circuit organization.

## Discussion

Understanding neural circuit organization at single-cell resolution remains a fundamental challenge in neuroscience, with sequencing-based approaches emerging as a promising alternative to traditional microscopy methods^56^. In this study, we developed Connectome-seq, demonstrating its ability to map neuronal connectivity while simultaneously capturing molecular identities of connected neurons. Applied to the mouse pontocerebellar circuit, this approach validated both established connectivity patterns and suggested unexpected circuit features that warrant further investigation.

The observation of potential pons-to-Purkinje cell connectivity in our dataset raises intriguing questions about cerebellar circuit organization. Through molecular characterization of these connected neurons, we identified several markers enriched in pons-connected Purkinje cells, with *Grid2ip*/Delphilin emerging as a particularly strong predictor of connectivity. This interface is where Purkinje cells extend various cellular processes into the granule layer and where pontine mossy-fiber terminals arborize^57,58^. The co-localization at this site raises the possibility that mossy fibers make contacts with Purkinje cell processes at this interface. However, further validation through complementary approaches, particularly electrophysiological recordings, will be essential to confirm these connections and understand their functional significance. The spatial intermingling of pontine axons targeting different cerebellar cell types suggests a more complex organization of the pontocerebellar circuit than previously appreciated, raising intriguing questions about the developmental mechanisms establishing these precise connectivity patterns.

Connectome-seq offers several advantages over existing approaches to neural circuit mapping. While serial electron microscopy reconstruction has provided foundational insights into circuit architecture at synaptic resolution, as demonstrated by recent studies in the cerebellar cortex^15^, it faces inherent limitations in scalability and throughput for mapping long-distance circuits across multiple brain regions. Light microscopy approaches such as array tomography and mGRASP allow synaptic resolution imaging but are limited by the need for thin sectioning and reconstruction of large volumes. Among recent sequencing-based neural connectivity mapping technologies, SYNseq first attempted to map synaptic connectivity by crosslinking pre- and post-synaptic proteins across synapses and joining their associated RNA barcodes through droplet-based RT-PCR^35^. Connectome-seq builds directly upon this pioneering conceptual framework while addressing the critical technical limitations in barcode joining efficiency through two key innovations: (1) elimination of the challenging droplet-based RT-PCR joining step by implementing direct barcode matching through parallel single-nucleus and single-synaptosome sequencing, and (2) use of substantially shorter, optimized RNA barcodes that achieve better synaptic transport and amplification efficiency compared to the longer sequences used in SYNseq. Other successful sequencing-based methods have improved scalability, though each faces distinct caveats: Sindbis virus-based methods enable high-throughput projection mapping but cannot reveal post-synaptic partners, while rabies virus-based methods allow synaptic connectivity mapping but struggle with barcode mixing in densely connected regions. Both viruses also face severe toxicity issues that may compromise circuit integrity. The technical advances in Connectome-seq, combined with AAV-based delivery for improved biocompatibility, enabled successful mapping of the pontocerebellar circuit and validation of novel connectivity patterns, demonstrating the potential for sequencing-based connectivity mapping to become a practical tool for circuit analysis.

A potential limitation of Connectome-seq involves the induced expression of modified neurexin and neuroligin proteins, which are known synaptogenic molecules that can influence synapse formation and stability. Previous studies have demonstrated that neurexin/neuroligin expression levels can affect synaptic connectivity, with overexpression potentially inducing ectopic synapse formation^59^. However, several factors suggest this concern may be mitigated in our experimental design. First, our viral expression is transient (2-3 weeks), limiting the duration of potential synaptogenic effects compared to chronic overexpression paradigms. Second, recent work using similar neurexin/neuroligin scaffolds has successfully mapped physiological connectivity patterns without apparent artifacts^60^, suggesting that moderate overexpression levels may not significantly perturb endogenous circuit organization. Additionally, our validation experiments using AAV1 anterograde tracing independently confirmed the pons-to-Purkinje cell connectivity patterns identified through Connectome-seq, providing orthogonal evidence that our findings reflect genuine circuit features rather than overexpression artifacts. Nevertheless, future iterations of this method should incorporate controls to assess potential synaptogenic effects, such as comparing connectivity patterns before and after viral expression or using alternative protein scaffolds with reduced synaptogenic activity. Additionally, Connectome-seq currently faces structural limitations, most notably restriction to long-distance projection mapping due to the requirement for anatomically separated viral injection sites. To overcome such limitation, future work could employ complementary Cre-on/Cre-off expression systems that would achieve mutually exclusive PreSynBar and PostSynBar expression within single brain regions, potentially enabling local circuit analysis by targeting different neuronal populations based on their molecular identities.

Through systematic optimization of molecular constructs, isolation protocols, and computational strategies, we achieved matching between synaptosomal barcodes and their nuclear sources at rates of 39.6% for PreRNA and 42.3% for PostRNA. These rates reflect several technical challenges in barcode recovery and matching. Source nuclei can be lost during tissue processing, particularly during dissection, homogenization, and flow cytometry enrichment where nuclear aggregates and compromised nuclei are excluded. Single-cell sequencing dropout effects may prevent capture of low-abundance barcodes in both nuclear and synaptosomal samples. Additionally, sequence mismatches are introduced during PCR amplification, especially given the higher number of PCR cycles required for synaptosomal samples. The impact of PCR-induced errors is evidenced by progressively increased matching rates when allowing more mismatches in the matching criteria, though this must be balanced against the introduction of false positives. The relatively low read counts per synaptosome also make the method susceptible to contamination from ambient RNA, potentially generating spurious connectivity calls. For the current study, we focused our analysis and validation efforts on high-confidence connections supported by multiple synaptosomal matches, particularly the consistent pons-to-Purkinje cell connectivity. Improving signal-to-noise ratios through better synaptosome isolation protocols and more stringent computational filtering will be crucial for future applications requiring high-confidence connectivity mapping. Our analysis identified several potential sources of technical artifacts: the detection of GABAergic neurons in pons projecting to cerebellum (9.2% of matched populations) likely represent either misassignment of barcodes to false source nuclei contaminated by RNA from true source nuclei, or false-positive matches from our barcode collapsing and matching algorithm. Using known non-projecting cells as negative controls, we estimated false discovery rates between 9.2%-11.7%.

To estimate the method’s sensitivity, we calculated the theoretical synaptic populations accessible to our sequenced cells. Given our sampling of 13.0% of total glutamatergic pontine neurons (78,358 out of 603,082, according to published dataset^61^) and 0.25% of cerebellar granule cells (49,385 out of ∼20 million from one third of cerebellum), combined with the known ∼200 billion total mossy fiber (MF)→GrC synapses in the cerebellum^15^, we estimate approximately 12.5% × 0.25% × 200 billion = 62.0 million theoretical MF→GrC synapses could be formed by our sequenced population. Similarly, with 1,423 cerebellar Golgi cells sampled and approximately 228 MF synapses per Golgi cell^62^, we estimate 13.0% × 1,423 × 228 = 42,178 theoretical MF→Golgi synapses in our sampled population. Our detection of 120 MF→GrC and 96 MF→Golgi connections therefore represent drastically different sensitivities of 0.0002% and 0.228% respectively for these connection types. This difference in sensitivity likely reflects distinct cellular architectures - the larger soma size and different synaptic organization of Golgi cells may facilitate both viral infection and RNA capture. While current sampling rates are extremely sparse due to technical limitations, several aspects of the method could be substantially improved. The 23.6% rate of dual-RNA capture from synaptosomes suggests room for optimization in synaptosomal isolation and RNA preservation protocols. The drop from 39.6% PreRNA and 42.3% PostRNA matching rates to 12.0% final validated connections (327 + 219 nuclei in the final connectivity matrix over 3,028 + 1,538 nuclei from the single-side-matched populations) indicates significant potential for improving barcode design, library construction, and matching algorithms. Additionally, cell-type specific differences in detection sensitivity point to opportunities for optimizing viral delivery and gene expression strategies. As these technical aspects get further improved, the detection of highly validated connections like MF→GrC and MF→Golgi circuits, along with potentially novel MF→Purkinje connectivity patterns and molecular markers such as *Grid2ip* that correlate with specific connections, suggests Connectome-seq could become a powerful tool for high-throughput circuit mapping across diverse brain regions.

The development of Connectome-seq represents a notable advance in our ability to understand neural circuit organization across multiple scales. By enabling systematic mapping of long-distance circuits with molecular precision, this approach bridges a critical gap between local circuit mapping and systems-level connectivity. Beyond the pontocerebellar system studied here, future applications of Connectome-seq could reveal organizational principles of other long-range circuits, particularly when integrated with complementary approaches like spatial transcriptomics and electrophysiological validation. As the technical limitations are addressed and sensitivity improved, this molecular understanding of circuit-specific connectivity patterns will be essential for deciphering how neural networks process information across brain regions, how circuits are established during development, and how their architecture may be disrupted in neurological disorders.

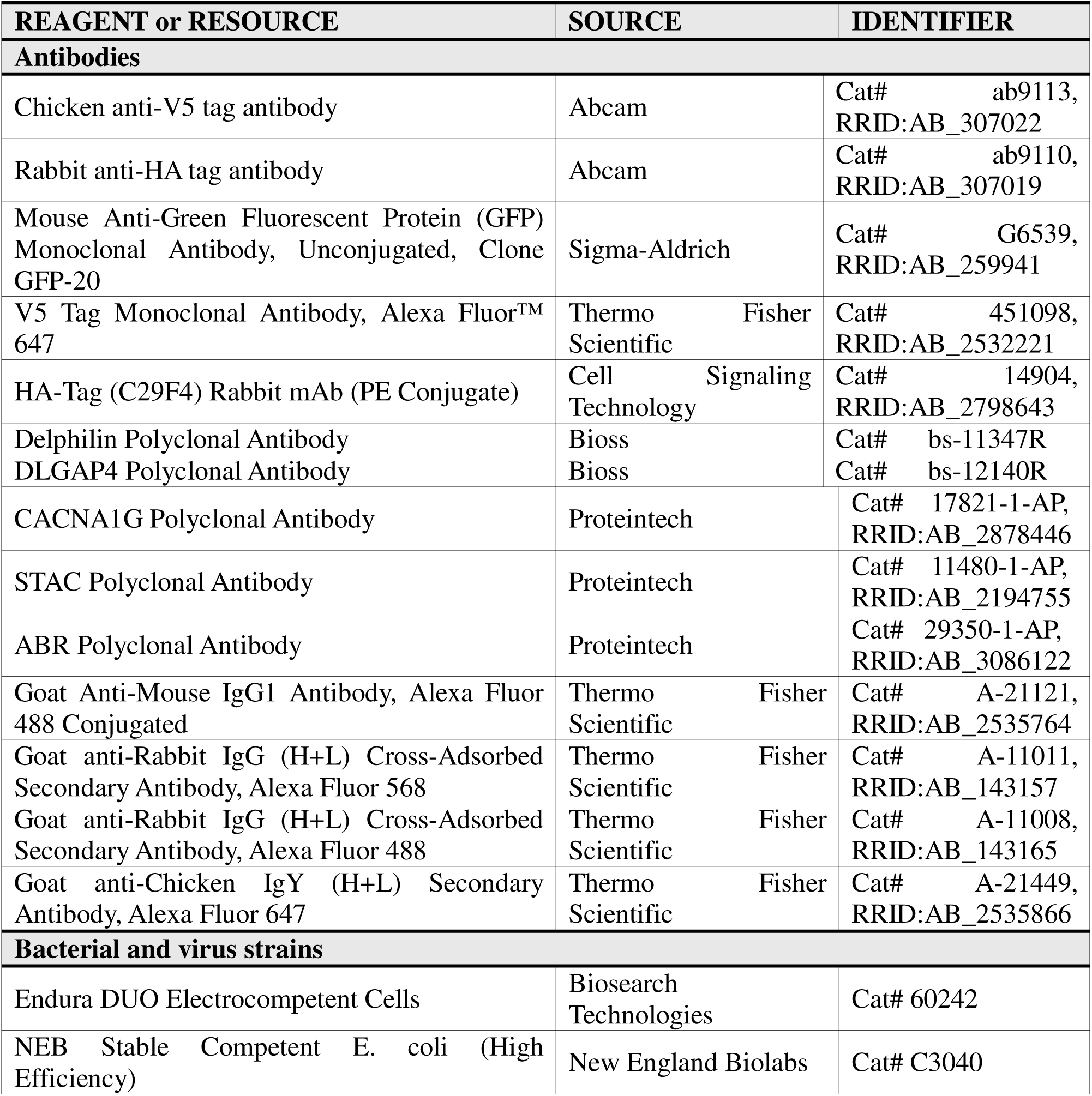
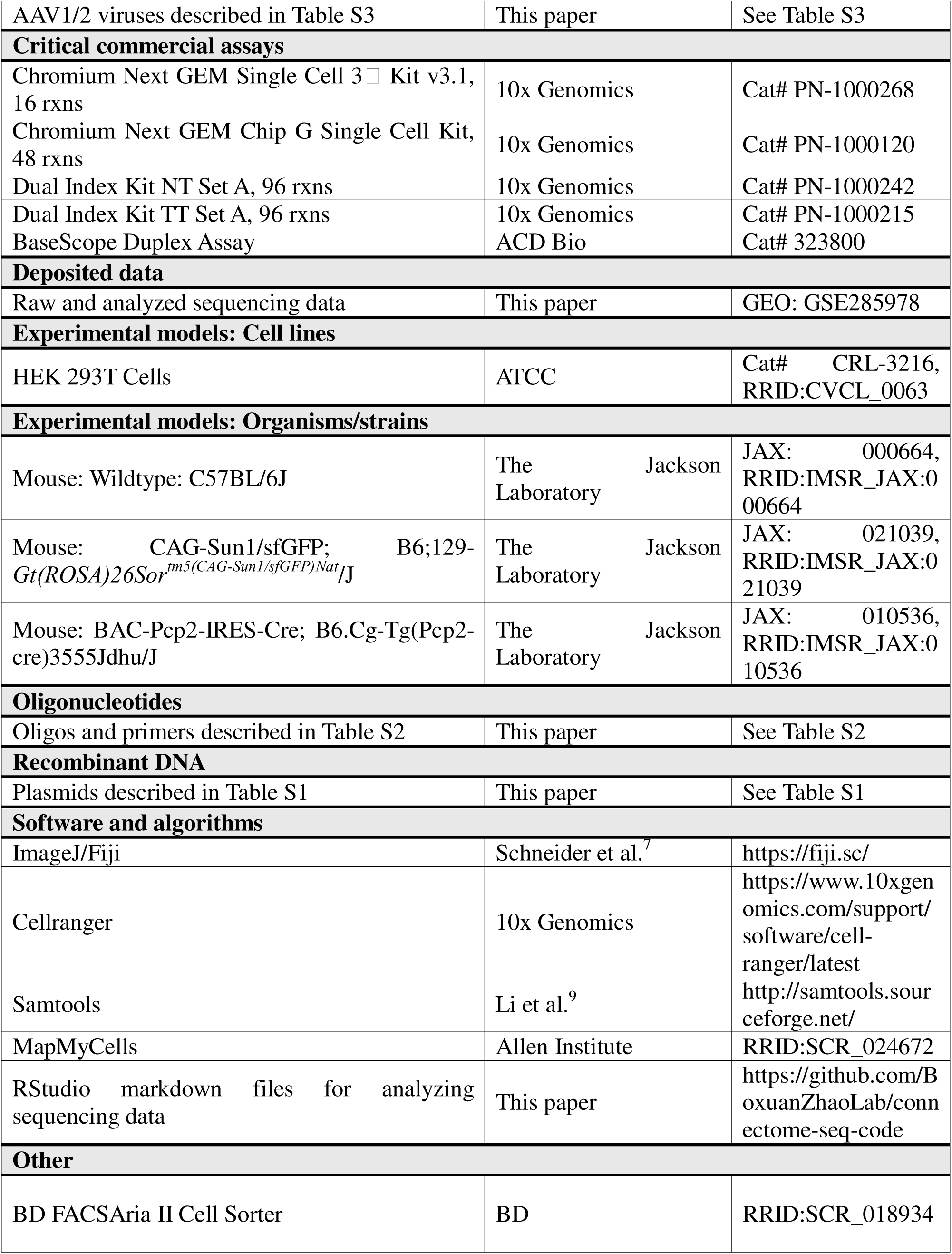

## Resource Availability

### Lead Contact

Further information and requests for resources and reagents should be directed to and will be fulfilled by the Lead Contact, Boxuan Zhao.

### Materials Availability

All unique reagents generated in this study are available from the lead contact.

### Data and Code Availability

Single-nucleus and single-synaptosome sequencing data reported in this paper were deposited into the Gene Expression Omnibus database under accession number GSE285978 and are available at the following URL: https://www.ncbi.nlm.nih.gov/geo/query/acc.cgi?acc=GSE285978. Code supporting the current study is available at https://www.github.com/BoxuanZhaoLab/connectome-seq-code and are available from the corresponding author on request.

## Experimental Model and Subject Details

### Cell Lines

HEK 293T cells (ATCC CRL-3216) were maintained in DMEM (Gibco) supplemented with 10% FBS (Gibco), penicillin-streptomycin (Corning, 5000 units/mL penicillin and 5000 μg/mL streptomycin) at 37°C, 5% CO2. This cell line has not been authenticated.

### Animals

All animal procedures were approved by Stanford University’s Institutional Animal Care and Use Committee (APLAC Protocol #: 14007) and University of Illinois Institutional Animal Care and Use Committee (IACUC Protocol #: 23169). For Connectome-seq experiments, 6-8 weeks old male/female Sun1-sfGFP-Myc (Jackson Laboratory, Stock #021039) transgenic mice were used. For imaging experiments, 6-8 weeks old male/female C57BL/6J (Jackson Laboratory, Stock #000664) mice were used. For AAV1 tracing experiments, 6-8 weeks old male/female Pcp2-Cre (Jackson Laboratory, Stock #010536) transgenic mice were used.

Mice were group-housed (2-5 per cage) on a 12h light/dark cycle with *ad libitum* access to food and water. Temperature was maintained at 20 ± 2°C and humidity at 50 ± 10%. Both male and female mice were used for experiments.

## Detailed Materials and Methods

### Molecular Biology and Viral Constructs

#### Plasmid Construction

Constructs were cloned into the pAAV viral vector (a gift from F. Zhang, MIT). Table S1 listed all plasmids used in this study. For all constructs, PCR fragments were amplified using Q5 High-Fidelity 2X Master Mix (NEB), and vectors were double digested with NEB restriction enzymes. Both fragments were purified with DNA gel and assembled using Gibson assembly. Assembled products were introduced into competent NEB Stable bacteria via heat shock transformation and correct clones were verified with Sanger sequencing.

#### Multiplexed Plasmid Library Construction

Barcoded RNA libraries were generated by converting single-stranded oligonucleotides containing random N30 sequences into double-stranded DNA through primer extension. Following exonuclease treatment to remove residual ssDNA, the purified dsDNA inserts were cloned into the vector backbone using Golden Gate Assembly. Assembly conditions were optimized using various vector:insert molar ratios (1:5 to 1:20). Large-scale library construction was performed using 3 μg vector DNA with the optimal ratio of insert in 150 μL reactions containing T4 DNA ligase buffer and Golden Gate Assembly enzyme mix. The ligated products were transformed into Endura electrocompetent cells (Lucigen), recovered, and expanded at 30°C. Library complexity was assessed through colony counting and Sanger sequencing of individual clones.

#### HEK 293T Cell Culture and Transfection

HEK 293T cells (ATCC) were maintained in High Glucose DMEM (Thermo) supplemented with 10% FBS (Corning), 1% (v/v) penicillin-streptomycin (Corning) at 37°C with 5% CO2. For transfections, cells were plated in 6-well plates at 5×10^5^ cells/well 24h before transfection. Cells at 70-80% confluency were transfected using polyethylenimine (PEI, 1 mg/mL in water, pH 7.3) at 3:1 PEI:DNA mass ratio. For each well, 2 μg total plasmid DNA was mixed with 6 μg PEI in 200 μL Opti-MEM (Thermo), incubated for 10 min at room temperature, then added dropwise to cells. Media was changed after 6 hours. For cis-assay, cells were co-transfected with 1:1 PreSynBar and PostSynBar plasmids. For trans-assay, cells were transfected with PreSynBar or PostSynBar separately, after 24 hours both populations of cells were lifted and replated into a separate pre-coated glass-bottom dish, with PostSynBar-transfected cells replated first and PreSynBar-transfected cells replated on top of them one hour later. The total density of cells was around 2/3 of the confluent amount of the new dish. Replated cells were incubated for another 24 hours followed by imaging.

#### HEK 293T Immunofluorescence Imaging

Cells to be imaged were plated on glass coverslips coated with 25 μg/mL poly-D-lysine in PBS for 1 h at 37°C. After transfection and replating, cells were fixed with 4% (v/v) paraformaldehyde in PBS for 15 min, washed two times with PBS, and permeabilized with ice-cold methanol at 4°C for 10 min or with 0.1% Triton X-100 in PBS at room temperature for 10 min. Cells were then washed two times with PBS and blocked for 1 h with 2% BSA (w/v) in PBS at room temperature. Samples were incubated with primary antibodies diluted 1:1000 in 1% BSA in PBS at 4°C overnight. We used anti-V5 antibody to detect PreSynBar, anti-HA antibody for PostSynBar, and anti-GFP antibody for split GFP. After primary antibody incubation, cells were washed three times in PBS and then incubated with secondary antibodies conjugated to Alexa Fluor-488/568/647 and DAPI diluted 1:1000 in 1% BSA in PBS for 90 min at room temperature. Cells were washed three times with PBS and mounted on glass slides and imaged by confocal fluorescence microscopy (Zeiss). Images were collected with Slidebook (Intelligent Imaging Innovations).

#### Concentrated AAV Production

Concentrated AAV virus was prepared for *in vivo* use as described previously^63^. Three 15-cm plates of HEK 293T cells with fewer than fifteen passages were transfected at 80% confluence. For each 15-cm plate, we combined 5.2 μg plasmid containing transgenes, 4.35 μg of AAV1 capsid plasmid, 4.35 μg of AAV2 capsid plasmid, and 10.4 μg pDF6 adenovirus helper plasmid with 130 μL PEI in 500 μL of serum-free DMEM for 10 min at room temperature. The media in the plates was then removed by aspiration and replaced with 20 mL of complete growth media plus the DNA mixture. HEK 293T cells were incubated for 48 h at 37 °C, supernatant was discarded, and the cell pellet was pooled together in PBS and collected by centrifugation at 800*g* at room temperature for 10 min. The pellet was resuspended in 20 mL of TBS solution (100 mM NaCl, 20 mM Tris, pH = 8.0), freshly made sodium deoxycholate solution (10% in H_2_O) was added to the resuspended cells to a final concentration of 0.5%, and benzonase nuclease (Sigma) was added to a final concentration of 50 U/mL. The mixture was shaken at 37 °C for 1 hour and cleared by centrifugation at 3,000*g* for 15 min. The supernatant was loaded using a peristaltic pump (Gilson MP4) at 1 mL/min flow rate onto a HiTrap heparin column (Cytiva) that was pre-equilibrated with 10 mL of TBS. The column was washed with 10 mL of TBS using the peristaltic pump, followed by washing with 1 mL of 200 mM NaCl, 20 mM Tris, pH = 8.0 and 1 mL of 300 mM NaCl, 20 mM Tris, pH = 8.0 using a 5 mL syringe. The virus was eluted using 5 mL syringes with 1.5 mL of 400 mM NaCl, 20 mM Tris, pH = 8.0; 3.0 mL of 450 mM NaCl, 20 mM Tris, pH = 8.0 and 1.5 mL of 500 mM NaCl, 20 mM Tris, pH = 8.0. The eluted virus was concentrated using Amicon Ultra 15 mL centrifugal units (100,000 MWCO, Millipore) at 2,000× g for 2 min, to a final volume of less than 500 μL. Then 1 mL sterile PBS was added to the filter unit and the column was centrifuged again for 1 min, followed by another wash with 500 μL PBS with 0.05% of PF-68 (Sigma) and centrifugation until the virus volume was ∼100 μL. The concentrated AAV virus was divided into 5 μL aliquots in PCS strips and flash frozen by liquid nitrogen followed by storage at −80 °C.

To titer the concentrated viruses, 2 μL of virus was incubated with 1 μL DNase I (NEB), 4 μL DNase I buffer, and 33 μL H_2_O at 37°C for 30 min followed by 75°C for 15 min. 5 μL of this reaction was then added to 14 μL H_2_O and 1 μL of Proteinase K (NEB) at 50°C for 30 min followed by 98°C for 10 min. qPCR reaction was prepared by adding 2 μL of this reaction to 5 μL of SYBR Green master mix (2x), 0.06 μL each of the forward and reverse primers (50 μM stock), and 2.88 μL of H_2_O. Primers were designed against the synapsin promoter and WPRE listed in Table S2. Standardized curves were generated using a purified linearized AAV transgene plasmid that contained synapsin promoter and WPRE at 0.05 ng, 0.1 ng, or 0.2 ng per μL. The titer of each viral sample was calculated in reference to the standard curve as fully described in ref ^63^.

#### AAV Genome Library Construction

Purified AAV (∼1×10^10 vg per sample) was treated with DNase I in a 40 µL reaction at 37 °C for 30 min and heat-inactivated at 75 °C for 15 min, followed by Proteinase K–based lysis with the Zymo Quick-DNA kit (Biological Fluids & Cells workflow) to degrade the capsid and release viral DNA. DNA was column-purified, eluted in 20 µL pre-warmed nuclease-free water (≥50 °C), quantified with 1 µL on a Qubit ssDNA High-Sensitivity assay, and the remainder kept on ice. To set the minimal unbiased amplification cycle number, 10 µL SYBR Green qPCR reactions were run with barcode-specific primers, and the preparative PCR cycle number was defined as the point of half-maximal fluorescence. Preparative PCR was performed in 50 µL with NEBNext Q5 Master Mix using the same barcode primers, and products were cleaned on Zymo DNA Clean & Concentrator (DCC) columns and eluted in 20 µL, with concentrations measured by NanoDrop. A second 10 µL SYBR Green qPCR with Illumina indexing primers (P5/P7) was used to determine cycle number corresponding to half of the maximum fluorescence for indexing, and final indexing PCRs were carried out in 25 µL with NEBNext Q5 Master Mix. Indexed products were cleaned on DCC columns, eluted in 20 µL nuclease-free water, quantified by NanoDrop, and submitted for QC and sequencing. Both PreRNA and PostRNA barcode libraries were sequenced at ∼50 million reads. Total projected diversity of the AAV library was calculated by *preseq* package^64^.

#### Primary Rat Cortical Neuron Culture and AAV Infection

Rat cortical neurons were isolated from E18 embryos following protocols approved by Stanford University’s IACUC. Culture plates and/or glass coverslips were coated with 0.001% (w/v) poly-l-ornithine (Sigma-Aldrich) in DPBS (Gibco) at room temperature for 2 hours, washed twice with DPBS, and subsequently coated with 5 μg/mL mouse laminin (Gibco) in DPBS at 37°C overnight. Cortical tissue was enzymatically digested in papain (Worthington) and DNase I (Roche) for 30 min at 37°C, then filtered through a 40 μm nylon cell strainer. Neurons were plated in 24-well plates at 1.25 × 10^6^ cells/well on glass coverslips or directly on the plate surface and maintained in complete neurobasal media (GIBCO) containing 2% B27 (Life Technologies), 1% Glutamax (Life Technologies), and 1% penicillin-streptomycin (VWR, 5 units/mL penicillin, 5 μg/mL streptomycin) at 37°C with 5% CO2. At DIV4, glial proliferation was suppressed by replacing 300 μL media with 500 μL complete neurobasal media supplemented with 10 μM FUDR (Sigma-Aldrich). Thereafter, 50% media changes with fresh complete neurobasal media were performed every 3 days. For viral transduction, neurons were infected at DIV7 with concentrated AAV at 8× 10^9^ viral genomes per well in 24-well plates. Media was changed 24 hours post-infection, and cells were maintained for at least 7 additional days to allow for robust transgene expression before experimental use.

#### Mouse stereotaxic surgeries

For stereotaxis surgeries, mice were anesthetized with 1.5%–2.0% isoflurane, and placed in a stereotaxic apparatus (Kopf Instruments) on a heating pad. The fur was removed from the scalp and a midline incision was made. We then drilled open a ∼0.3 mm diameter craniotomy over the injection sites. Using a glass pipette (30–50 μm tip diameter), we injected 500 nL of AAV at 100 nL/min to each injection site. The following coordinates (in mm relative to lambda unless otherwise noted) were used for viral injections: Bilateral pons: +0.2 A/P, ±0.5 M/L, −5.55 D/V; Right cerebellum (right vermis of cerebellar lobule V): +0.2 mm A/P anterior to the post-lambda fissure on the cranial midline. −3.0 M/L, −0.5 mm D/V below dura. We allowed the virus to incubate for 2-3 weeks before experiments. The virus titers used for each experiment are listed in Table S3.

#### Mouse histology and immunohistochemistry

Following virus injection and expression, mice were heavily anesthetized with isoflurane and then perfused with 20 mL of cold phosphate-buffered saline (PBS) followed by 20 mL of cold 4% paraformaldehyde (PFA) in PBS. The brain was extracted from the skull and incubated in PFA at 4°C overnight, and then transferred to 30% sucrose for another incubation at 4°C overnight. The brain was sliced on a vibratome (Leica) or a cryotome (Leica) in 100-μm sections and mounted on glass slides with coverslips for imaging. For immunohistochemistry of PreSynBar,

PostSynBar, or SEMA3C, slices were blocked and permeabilized in PBS with 0.3% Triton-X (PBST) and 5% normal goat serum (NGS) at room temperature for 2 hours. Slices were stained in primary antibodies in PBST+5% NGS at 4°C overnight, followed by three washes with PBST, and staining with secondary antibodies in PBST+5% NGS at room temperature for 90 min. Slices were washed three times in PBST and mounted on glass slides with mounting media containing DAPI for imaging. Imaging was performed using similar methods described for HEK 293T cells.

#### BaseScope In Situ Hybridization and Immunofluorescence

We combined the workflow of the standard BaseScope Duplex Detection kit (Advanced Cell Diagnostics) and the tech note workflow of RNAscope or BaseScope RED Assay combined with Immunohistochemistry - Integrated Co-Detection Workflow (ICW) to enable co-detection of both barcodes and SynBar proteins. Briefly, 10 μm Cryosections were fixed in 4% PFA for 1 hour at room temperature after brief heat treatment (100°C, 2 min). Primary antibodies were applied in co-detection diluent overnight at 4°C. Following PBST washes and post-fixation (4% PFA, 30 min), sections were treated with hydrogen peroxide for 5 min and BLOXALL Endogenous Blocking Solution (Vector Laboratories) for 10 min, followed by protease IV treatment for 30 min at 40°C. Probes (Human PPIB-C1 control probes for PreRNA, DapB-C2 control probes for PostRNA) were hybridized for 2 hours at 40°C followed by sequential amplification steps using the BaseScope Duplex Detection kit. All 12 amplification steps and both Fast Red development (2 min, room temperature) and Green development (40°C, 5 min) were used. All washing steps between amplifications used BaseScope wash buffer at room temperature. Co-detection blocking was carried out (40°C, 15 min), followed by secondary antibodies incubation overnight at 4°C. Sections were mounted and imaged immediately after final PBST washes.

For RNA-protein colocalization analysis using FIJI (ImageJ): Binary masks were generated for both RNA (FISH) and protein (immunofluorescence) channels using automatic Triangle thresholding for protein signals and either 40% of maximum intensity (PreRNA) or automatic Triangle thresholding (PostRNA) for RNA signals. For proximity analysis, RNA masks were dilated by 1 pixel using the Maximum filter. Direct overlap was calculated as the intersection of RNA and protein masks. Proximity overlap was determined as the intersection between dilated RNA masks and protein masks. The percentage of RNA in protein was calculated as (overlap area / total RNA area) × 100. For each condition, three independent samples were analyzed. Statistical significance was assessed using two-tailed t-tests.

#### Parallel Nuclear and Synaptosomal Isolation

Fresh brain tissue was extracted and processed immediately after perfusing the mice with 10 mL ice-cold PBS. Target brain regions (right 1/3 of the cerebellum or bilateral pons) were dissected with new blades and homogenized in 1 mL SET buffer (320 mM sucrose, 5 mM Tris-HCl pH 7.5, 1 mM EDTA, 1:100 diluted Protease Inhibitor Cocktail (Sigma), 0.2 U/μL RNase inhibitor (Lucigen)) on ice using a glass dounce homogenizer (15 strokes) followed by additional homogenization with another 0.4 mL fresh SET buffer (5 strokes). The homogenate was combined and centrifuged at 1,500g for 8 min at 4°C to separate the nuclear pellet (P1) and synaptosomal supernatant (S1).

For nuclear isolation, the P1 fraction was briefly rinsed with nuclear lysis buffer (10 mM Tris-HCl pH 7.4, 10 mM NaCl, 3 mM MgCl2, 0.1% NP-40, 1:100 diluted Protease Inhibitor Cocktail (Sigma), 0.2 U/μL RNase inhibitor (Lucigen)) and then resuspended in 600 μL fresh lysis buffer. Cerebellar samples were further homogenized with 30 strokes using a plastic pestle, while pontine samples were gently mixed. Following lysis, samples were diluted with an equal volume of lysis buffer and incubated on ice for 10 min. Nuclear suspensions were filtered and pelleted at 1,500g for 5 min. The nuclear pellet was washed once with 800 μL nuclear lysis buffer, then resuspended in 800 μL washing and staining buffer (1X PBS, 1% BSA, 0.2 U/μL RNase inhibitor (Lucigen)). After 3 min incubation on ice, nuclei were pelleted (1,500g, 5 min), resuspended in 600 μL washing and staining buffer containing SiR-DNA at 1:200 final dilution ratio, filtered, and sent for flow cytometry.

For synaptosome isolation, the S1 fraction was centrifuged at 12,600g for 16 min at 4°C to obtain the crude synaptosomal pellet (P2). The P2 fraction was gently resuspended in 600 μL washing and staining buffer, 200 μL crude synaptosomes were taken out and blocked by adding 2 μL Fc receptor blocker (1:100) for 5 min on ice. Synaptosomes were then stained with anti-V5-Alexa Fluor 647 (1:50) and anti-HA-PE (1:50) antibodies for 1 hour at 4°C by rotation incubation. After washing with 400 μL washing and staining buffer (6,000g, 5 min), samples were resuspended in 500 μL washing and staining buffer. 200 μL of the suspension was diluted in 800 μL washing and staining buffer containing 1:10,000 diluted CellBrite 405 membrane dye, filtered, and sent for flow cytometry.

#### Parallel Nuclear and Synaptosomal Flow Cytometry

Single nucleus sorting: Nuclei were sorted on a BD FACSAria II flow cytometer using SiR-DNA (1:200) for DNA content and Sun1-Tag GFP for infected neuron identification. Sequential gating was performed to first select singlet nuclei based on SiR-DNA intensity, followed by GFP-positive event selection. Sorted nuclei were collected directly into pre-wet PCR strip tubes containing 5 μL collection buffer (PBS with 1% BSA and 4U/μL Protector RNase Inhibitor (Roche)) for downstream processing.

Synaptosomes were sorted using a reconfigured BD FACSAria II flow cytometer. To optimize synaptosome detection, the standard optical configuration was modified: the bandpass filter was removed from the V450 detector to create a second side scatter channel (SSC), and the V525 detector was fitted with a 450/50 bandpass filter and a 410 longpass filter to enable violet fluorescence detection of CellBrite 405. Sequential gating was employed to enrich synaptosomes. First, events were gated based on dual side scatter measurements to identify events above the noise floor threshold, comparing with a buffer-only negative control sample. Second, singlet synaptosomes were selected based on their forward scatter (FSC) and side scatter (SSC) ratios, corresponding to the expected size and internal complexity of synaptosomes. Third, membrane integrity was assessed using CellBrite Steady 405 dye, with intact synaptosomes selected based on the top ∼90% intensity of the CellBrite 405 signal. Finally, double-positive synaptosomes, representing connected pre- and post-synaptic terminals, were identified based on V5 and HA fluorescence signals. Thresholds for V5 (<0.5% false positive) and HA (<1% false positive) were empirically determined using wildtype control samples to minimize the inclusion of non-specific events. Approximately 20,000 double-positive synaptosomes were collected per hour into 5 μL of collection buffer for downstream processing. The BD FACSAria II, with its cuvette-based flow cell enabling longer laser exposure time and gel-coupled light collection, along with its continuous spectral detection range, was chosen for its superior ability to consistently detect and isolate singlet synaptosome populations compared to other platforms with segmented spectral detection or plastic-based flow cells.

#### RT-qPCR of Barcode RNA from Sorted Synaptosomes

150,000 synaptosomes were sorted by flow cytometry and collected into pre-wetted centrifuge tubes containing 1 µL NxGen RNase inhibitor. Samples were centrifuged at 20,000 g for 25–30 min at 4 °C, leaving ∼5 µL residual volume after removing the supernatant. Pellets were lysed by adding 7.15 µL of freshly prepared lysis/primer mix (5 µL H_2_O, 0.15 µL 10% Triton X-100, 0.5 µL 50 µM Oligo-dT primer, 0.5 µL 50 µM barcode RT primer, 1 µL 10 mM dNTP), incubated at 65 °C for 5 min, briefly spun, and placed on ice for 1 min. cDNA synthesis was performed using LunaScript RT SuperMix (5×, NEB). TaqMan qPCR reactions were then run in 10 µL volumes using TaqMan Fast Advanced Master Mix (2×, ThermoFisher) with barcode-specific forward and reverse primers, probe, cDNA template, and nuclease-free water.

#### Single-Nucleus and Single-Synaptosome Sequencing Libraries Construction and Sequencing

Libraries were generated using the Chromium Next GEM Single Cell 3 GEM, Library & Gel Bead Kit v3.1, Chromium Next GEM Chip G, and 10x Chromium Controller (10x Genomics) according to manufacturer’s protocol with modifications. For each experiment, approximately 20,000-30,000 individually sorted GFP-positive nuclei or double-positive synaptosomes were loaded into each channel of the chip. For nuclear libraries, cDNA was amplified, size-selected using SPRIselect beads to separate transcriptome and barcode components, and independently constructed into mRNA libraries and barcode libraries. Synaptosome libraries were constructed using barcode-specific amplification without template switching oligonucleotides. The resulting libraries were quality-controlled using Bioanalyzer High Sensitivity DNA Analysis (Agilent), quantified by Qubit 4 Fluorometer (ThermoFisher), and pooled for sequencing on an Illumina NovaSeq X platform.

#### AAV1-labeled Anterograde Tracing Single-Nucleus RNA Sequencing

For validation of connectivity-associated marker genes, we performed single-nucleus RNA sequencing on cerebellar neurons infected by anterograde-transported AAV1. Adult Sun1-Tag mice (6-8 weeks) were stereotaxically injected with 500 nL of AAV1-hSyn-Cre virus bilaterally into the pons using coordinates: +0.2 A/P, ±0.5 M/L, −5.55 D/V relative to lambda. This injection strategy leverages AAV1’s trans-synaptic transport capability to deliver Cre recombinase to pontine-connected cerebellar neurons, thereby activating Sun1-Tag expression specifically in pons-receiving cells.

After 3 weeks expression, mice were perfused with ice-cold PBS and cerebellar tissue was rapidly dissected and processed for nuclear isolation as described in the main methods. GFP-positive nuclei (representing pons-connected neurons) were sorted by flow cytometry using the same protocol described for Connectome-seq experiments. Sorted nuclei were immediately loaded onto 10x Genomics Chromium Single Cell 3 platform and processed according to manufacturer’s protocols. Libraries were sequenced on Illumina NovaSeq platform and analyzed using Cell Ranger followed by standard single-cell analysis pipelines in R/Seurat to determine cell type identities of AAV1-labeled neurons.

### Computational Analysis

#### Raw Data Processing and Barcode Extraction

Raw sequencing data was processed using Cell Ranger (v7.1.0) with custom genome references including SynBar and Cre genes. Nuclear libraries were processed using force-cells parameter of actual loaded amount, while synaptosomal libraries used expect-cells parameter of actual loaded amount. Barcode sequences were extracted from filtered nuclear and unfiltered synaptosomal BAM files using custom Python scripts. Sequences were collapsed using Hamming distance 1 within samples to account for sequencing errors.

#### Single Nucleus RNA-seq Analysis

Initial data processing was performed for each sample independently. Raw data was loaded using Read10X, including both gene expression and synapse barcodes (SynBarpost, SynBarpre). Seurat objects were created with gene expression data as the default assay, while synapse barcodes were stored as separate assay. Quality control metrics were calculated including the percentage of mitochondrial genes (pattern “^mt-”), number of features, and total counts per cell. Cells were filtered to retain those with >100 and <8000 features and <5% mitochondrial reads. Data normalization was performed using LogNormalize with a scale factor of 10,000, followed by identification of 2,000 variable features using variance-stabilizing transformation (VST) method. After scaling all genes, principal component analysis was performed.

For multi-sample analysis, the six independent samples were integrated using Harmony to remove batch effects. For cerebellum data, after Harmony integration, clustering was performed using the standard Seurat pipeline^65^, with gene expression data normalized using LogNormalize with scale factor 10,000, followed by identification of 2,000 variable features using variance-stabilizing transformation. After data scaling, principal component analysis was performed. UMAP embedding was carried out at multiple resolutions (0.2-0.8) with 0.4 selected as optimal resolution based on cluster separation and biological relevance. Pons nuclei data were processed with a custom pipeline modified from a published single nucleus sequencing pipeline^66^. Briefly, variable genes were identified using a binomial model that compared expected versus observed expression frequencies across cells, selecting genes where the observed proportion was at least 5% less than the expected proportion based on bulk expression. Expression data was square root transformed, and a k-nearest neighbor graph was constructed using cosine distance on the first 30 harmony-corrected principal components. Clustering was performed using the Louvain algorithm across multiple resolution parameters (0.6-2.0), with the optimal resolution (1.0) selected based on cluster size criteria (minimum 500 cells) and cluster separation.

For both datasets, small clusters (<10 cells) were merged with their most transcriptionally similar neighbors based on correlation of average expression profiles. Clusters with median gene counts below 1,000 were filtered out (clusters 0, 2, 37 and 38 for pons; clusters 0, 19, 22 and 23 for cerebellum). Cell types were initially annotated using canonical marker genes, including neurotransmitter genes for pons described in ref^67^ and published cell type markers for cerebellum described in ref^45^. Cluster identities were further annotated using Allen Institute’s MapMyCell annotation tool, with each cluster assigned to the cell type showing the highest annotation counts. For the pons nuclei data, cluster annotations were manually refined by examining UMAP coordinates. Specifically, 248 cells originally assigned to cluster 27 and positioned below a UMAP Y-axis threshold were reassigned to cluster 10 to correct for over aggregation. Initial subclusters were merged into major cell types based on shared marker expression patterns and consistent MapMyCell annotations.

### Connectivity Analysis

Nuclear and synaptosomal barcode sequences from neurons were matched using a stepwise filtering approach. Initial pre-filtering removed ambiguous nuclear barcodes where the highest count was not at least 5 times (for pons) or 2 times (for cerebellum) greater than the second highest count, and synaptosomes associated with more than 100 unique barcode sequences. Matching between nuclear and synaptosomal barcodes was performed iteratively, allowing 0-10 hamming distance mismatches. Potential artifacts were removed through multiple filtering steps: nuclear and synaptosomal barcode reads with identical 10x Cell Barcodes were eliminated to avoid misassignment, barcodes associated with more than 100 synapses were removed to control for ambient RNA, and for cases with one synaptosome matching to multiple nuclei, only the pair with the highest synaptic count followed by highest nuclear count was retained. The filtered matches were used to construct cell-type grouped connectivity matrices based on shared barcodes between pre- and post-synaptic populations.

### Differential Expression Analysis

Purkinje cell data was extracted from cerebellum nuclei data based on cell type annotation and scVIIntegration^68,69^ was used to remove batch effect. Purkinje cell connectivity-associated marker gene expression was analyzed using Seurat’s FindMarkers function to compare pons-connection positive clusters and negative clusters. Analysis parameters required genes to be expressed in at least 25% of cells in either population with a minimum log fold-change threshold of 0.25. Statistical significance was determined by using adjusted p-values to account for multiple comparisons. Top differentially expressed genes were identified for each comparison and visualized using normalized expression data across conditions.

### Statistical Analysis

Statistical analyses were performed with at least two biological replicates, with most key experiments using three or more replicates. Specific statistical tests included: Unpaired two-tailed Student’s t-test for comparing expression levels between two conditions (e.g., RNA Polymerase II vs III-driven expression); One-way ANOVA followed by Dunnett’s multiple comparisons test for comparing multiple conditions to a control (e.g., different RNA targeting sequences); Paired t-test for comparing high and low abundance samples in PCR analysis; Hierarchical clustering using Ward’s method (ward.D2) for connectivity analysis; For differential gene expression analysis, statistical significance was assessed using adjusted p-values to control for multiple comparisons. Data are presented as mean ± SEM unless otherwise noted in figure legend. Statistical significance was typically indicated as: *p < 0.05, **p < 0.01, ***p < 0.001; ns, not significant. No statistical methods were used to pre-determine sample size or fit with assumptions of statistical tests. No subjects with successful library construction were excluded from the study. Nuclei not passing quality control metrics defined in “Single Nucleus RNA-seq Analysis” were omitted from the bioinformatic analysis.

## Acknowledgments

We are deeply grateful to Drs. Alice Y. Ting and Liqun Luo (Stanford University) for their foundational contributions to this work, including the initial conception of the method, guidance throughout the experimental development, and critical feedback on the manuscript. Their insights were instrumental in shaping both the technical approach and the scientific rigor of this study. We thank Dr. Steve R. Quake (Stanford University) for providing guidance to the project and critical sequencing resources. We thank Dr. Etienne Herzog (The University of Bordeaux) and Dr. Meredith Weglarz (Stanford University - Shared FACS Facility) for assistance with protocol optimization for single synaptosome sorting experiments. Nuclei and synaptosome flow cytometry sorting and analysis for this project was done on an instrument in the Stanford Shared FACS Facility (RRID: SCR_017788) obtained using NIH S10 Shared Instrument Grant S10RR025518-01. This research was conducted with support from the Neuro-omics Initiative grant from Wu Tsai Neurosciences Institute of Stanford University. B.S.Z is funded by an Elsa U. Pardee Foundation Research Grant and an Edward Mallinckrodt Jr. Foundation Grant.

## Contributions

B.S.Z. conceived and designed the study, performed all experiments with D.C. and A.I., analyzed the data with A.I., Z.W., and wrote the manuscript. B.S.Z. and A.I. designed the sequencing workflow, performed sequencing experiments, and built the bioinformatic pipeline with help from Z.W. and Y.W.. D.C. performed validation experiments on cell type specific connectivity. A.I. and Z.W. helped with data analysis and visualization. M.J.W. helped design the AAV injection scheme and perform stereotaxic injections. Y.W. helped with Basescope RNA imaging experiments. All authors reviewed the manuscript and provided critical feedback.

---

**Extended Data Figure 1:**
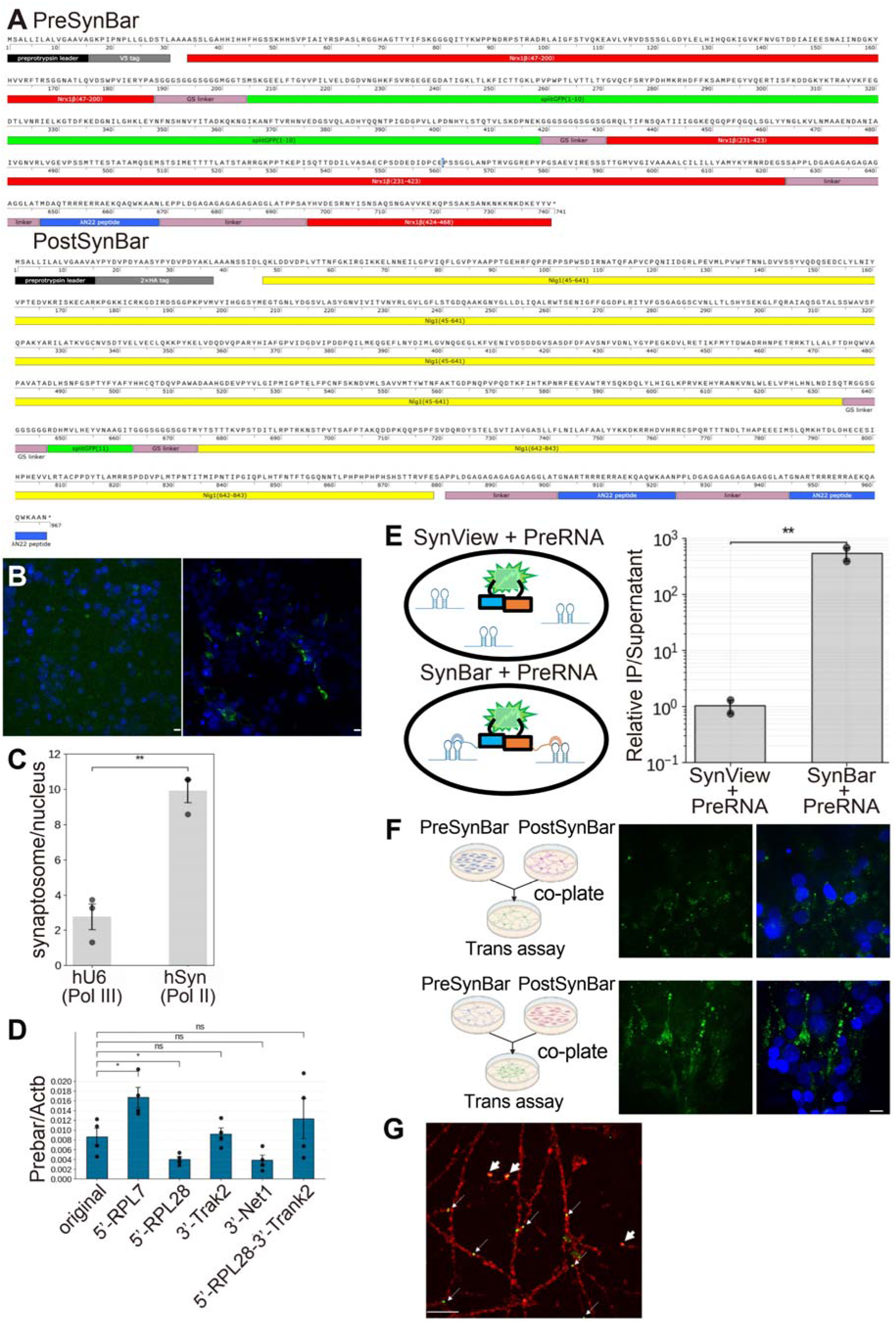
Design and Optimization of SynBar Protein-RNA Complex. (A) Protein domain organization of PreSynBar and PostSynBar constructs. PreSynBar contains modified neurexin 1β (red) with embedded split-GFP(1-10) (green), V5 tag (grey), and λN22 RNA-binding domain (blue). PostSynBar contains modified neuroligin 1 (yellow) with embedded split-GFP(11) (green), HA tag (grey), and λN22 domain (blue). Numbers indicate amino acid positions. (B) Trans-interaction assay in HEK 293 cells demonstrating the importance of λN22 positioning in PreSynBar. Left: PreSynBar with C-terminal λN22 failed to show proper interaction with PostSynBar, so no split GFP signal was observed. Right: PreSynBar with internal C-terminal λN22 showed robust trans-synaptic interaction with PostSynBar evidenced by clear split GFP signals. Scale bars: 10 μm. Blue: DAPI; Green: reconstituted split GFP signal. (C) Comparison of barcode expression levels under different promoters. RNA Polymerase II-driven expression (hSyn promoter) shows higher barcode abundance in the synaptosomal fraction (normalized to nuclear fraction) compared to Polymerase III-driven expression (hU6 promoter). Data are represented as mean ± SEM from n = 3 biological replicates. Statistical significance was determined using unpaired two-tailed Student’s t-test (**p < 0.01). (D) Optimization of PreRNA localization using various targeting sequences. After co-injecting AAVs expressing PreSynBar and different PreRNA designs into presynaptic brain regions, the abundance of PreRNA in postsynaptic synaptosomal fraction were measured by qPCR (normalized to housekeeping gene Actb) for five different barcode designs: Original unmodified PreRNA, RPL7 5-UTR, RPL28 5-UTR, Trak2 3-UTR, Net1 3-UTR, and combined RPL28 5-UTR + Trak2 3-UTRs. RPL7 5-UTR shows highest enrichment. Data are represented as mean ± SEM from n = 4 biological replicates. Statistical significance was determined using one-way ANOVA followed by Dunnett’s multiple comparisons test comparing all conditions to Original (*p < 0.05; ns, not significant). (E) RNA immunoprecipitation (RIP) demonstrating stable λN22-BoxB RNA interaction. Left: Schematic showing experimental design – HEK 293 cells were co-transfected with either PreSynBar+PostSynBar or PreSynView+PostSynView (control lacking λN22 domains) along with PreRNA containing BoxB sequences. Right: Quantification of PreRNA enrichment in anti-GFP immunoprecipitates versus supernatant, shown as log10 ratio. SynBar shows >500-fold higher RNA enrichment compared to SynView control, confirming stable and specific BoxB-λN22 interaction. Data are represented as mean ± SEM from n = 2 biological replicates. Statistical significance was determined using unpaired two-tailed Student’s t-test on log-transformed values (**p < 0.01). (F) Trans-interaction assay validating SynBar interaction between different cell types. Left: Experimental schematic showing separate plating and co-culture conditions: HEK 293 cells expressing PreSynBar with neurons expressing PostSynBar (top), and neurons expressing PreSynBar with HEK 293 cells expressing PostSynBar (bottom). Right: Representative immunofluorescence images showing specific GFP reconstitution (green) at contact points between the different cell types in both configurations. Scale bar: 10 μm. Blue: DAPI nuclear staining. (G) Confocal microscopy image of cultured neurons with PreRNA FISH. PreRNA barcode signals (green) colocalize with PreSynBar signals (red). Thin arrows indicate PreRNA barcodes in transit along neuronal processes, while thick arrows highlight PreRNA barcode signals localized at synaptic terminals. Scale bars: 10 μm.

**Extended Data Figure 2:**
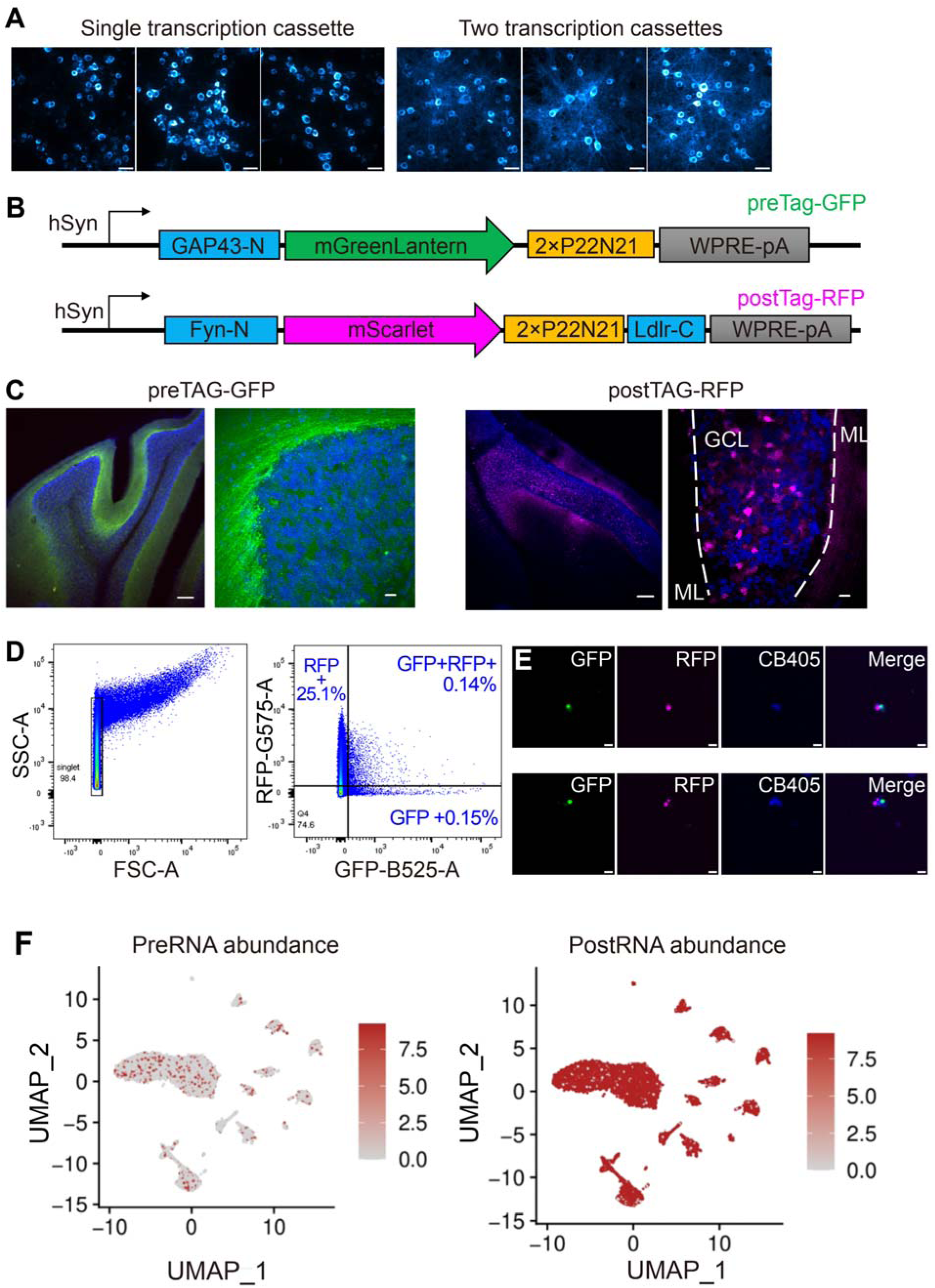
AAV Construct Optimization, and Alternative Fluorescent Protein Approach. (A) Confocal microscopy images illustrating the optimization of PreSynBar expression and localization in cultured neurons. Upper: Representative image of neurons infected with an AAV carrying a single transcriptional cassette driving the expression of V5-tagged PreSynBar. The protein shows predominantly somatic localization with limited distribution into neuronal processes. Lower: Representative image of neurons infected with an AAV carrying two transcriptional cassettes: one driving V5-tagged PreSynBar expression and another driving PreRNA expression. This design demonstrates more extensive localization of PreSynBar throughout neuronal processes. Note the comparable expression levels of PreSynBar in the soma between the two conditions. Pseudo color: V5 staining. Scale bars: 10 μm. (B) Design of preTag-GFP and postTag-RFP constructs. PreTag-GFP contains GAP43-N targeting sequence, mGreenLantern, and RNA-binding domain (2×P22N21). PostTag-RFP contains Fyn-N and Ltdr-C targeting sequences, oScarlet, and RNA-binding domain. Both constructs are driven by hSyn promoter. (C) In vivo expression pattern of fluorescent protein constructs. Left: PreTag-GFP shows strong axonal localization in cerebellar molecular layer (ML). Right: PostTag-RFP demonstrates dendritic targeting in granule cell layer (GCL). Blue: DAPI. Scale bars: left panels 100 μm, right panels 10 μm. (D) Synaptosome sorting strategy using native fluorescence. Left: Initial singlet gating based on scatter properties. Right: Identification of GFP+, RFP+, and double-positive populations. (E) Imaging validation of sorted synaptosomes. Representative confocal images of sorted double-positive synaptosomes showing membrane staining (CellBrite405, blue), RFP signal (red), and GFP signal (green). Scale bars: 2 μm. (F) Analysis of barcode abundance in cerebellar nuclei expressing fluorescent constructs. UMAP visualization of single-nucleus RNA-seq data showing PreRNA (left) and PostRNA (right) expression levels. Cerebellar nuclei were infected with PostRNA-expressing virus, and PreRNA signal came from contamination of ambient PreRNA from pons projections. Color intensity indicates barcode abundance in read counts.

**Extended Data Figure 3:**
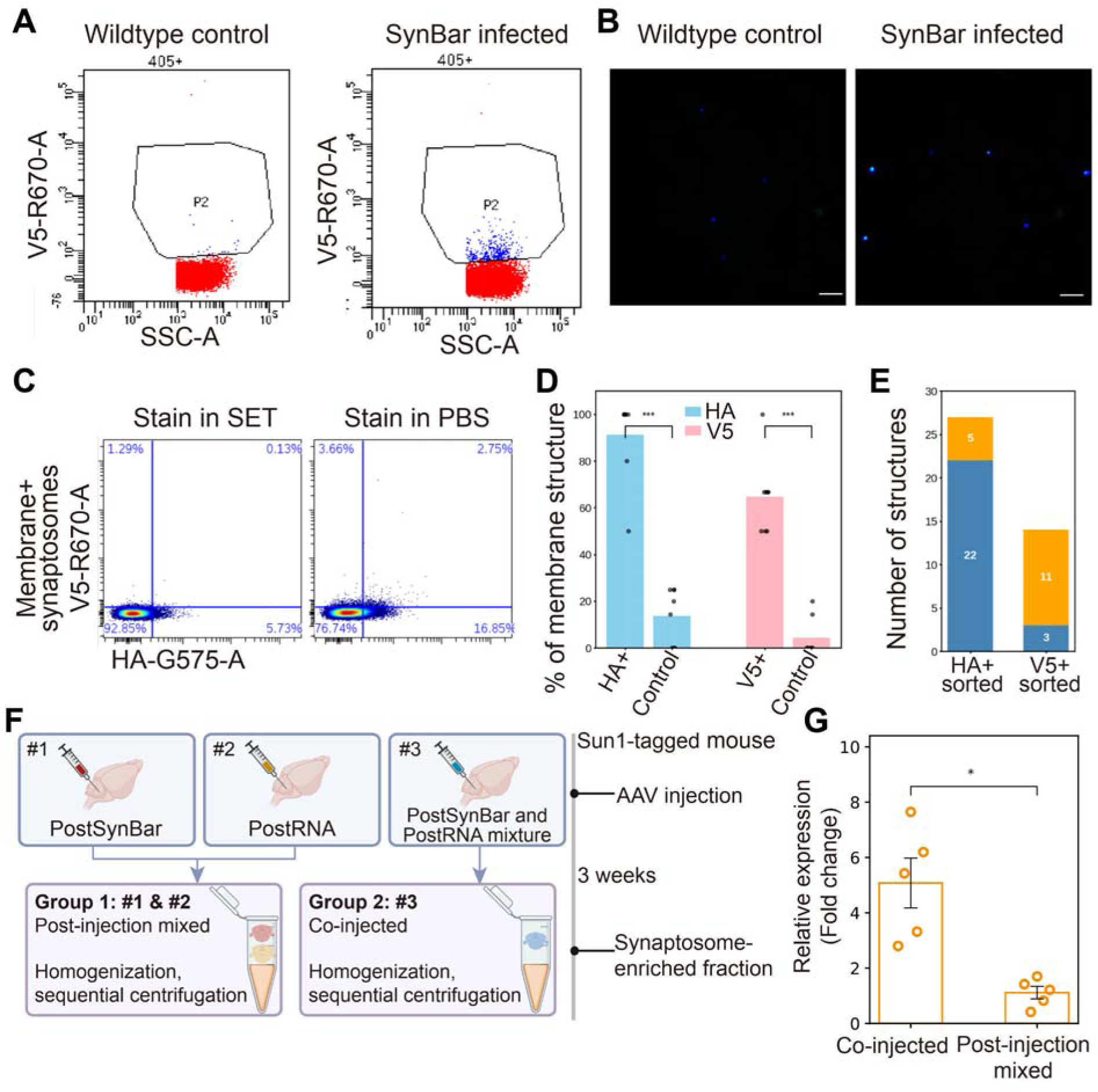
Optimization of Synaptosome Immunostaining. (A) Flow cytometry detection of reconstituted GFP using membrane+ gating. Wildtype control (left) versus SynBar-infected sample (right) shows specific detection of GFP+ synaptosomes (1.3% of membrane+ population) in infected tissue only. (B) Confocal imaging validation of sorted GFP+ synaptosomes from wildtype and SynBar-infected samples. Blue: CellBrite405 membrane dye; Green: anti-GFP signal. Scale bar: 10 μm. (C) Comparison of antibody staining efficiency in different buffers. Staining in sucrose-based SET buffer (left) shows reduced signal compared to PBS-based buffer (right), with higher detection of positive populations in PBS (2.75% vs 0.13% double-positive). (D) Quantification of marker presence in sorted populations. Bar graph showing the percentage of membrane-intact structures (based on CellBrite 405 signal) that are positive for HA or V5 in HA+ sorted populations versus control populations or V5+ sorted populations versus control populations. Error bars represent SEM; ***p < 0.001. (E) Analysis of single versus double-positive structures across sorted populations. Bar graph showing the number of structures that are either single positive (blue) or double positive (orange) for V5 and HA markers in HA+ sorted and V5+ sorted populations. (F) Design of the PostRNA contamination test. Three Sun1-tagged mice were used for this experiment: Mouse #1: Injected with only PostSynBar virus into the cerebellum; #2: Injected with only PostRNA virus into the cerebellum; Mouse #3: Co-injected with a mixture of PostSynBar and PostRNA viruses into the cerebellum. Three weeks post-injection, cerebella from these mice were harvested and divided into two experimental groups: Co-injected group: Cerebellum from mouse #3; Post-injection mixed group: Combined cerebellar tissues from mice #1 and #2. Synaptosome-enriched fractions were obtained through homogenization and sequential centrifugation and PostSynBar-positive population were collected based on HA staining by employing flow cytometry. PostRNA contamination was quantitatively assessed using qPCR from equal numbers (150,000) of HA+ synaptosomes collected from the two samples. (G) qPCR analysis of PostRNA barcode expression in sorted cerebellar synaptosomes. “Co-injected” indicates synaptosomes sorted from cerebellum injected with a mixture of PostSynBar and PostRNA viruses. “Post-injection mixed” refers to synaptosomes sorted from pooled cerebellar tissues injected separately with either PostSynBar or PostRNA viruses. Data are presented as mean ± SEM; *p < 0.05.

**Extended Data Figure 4.**
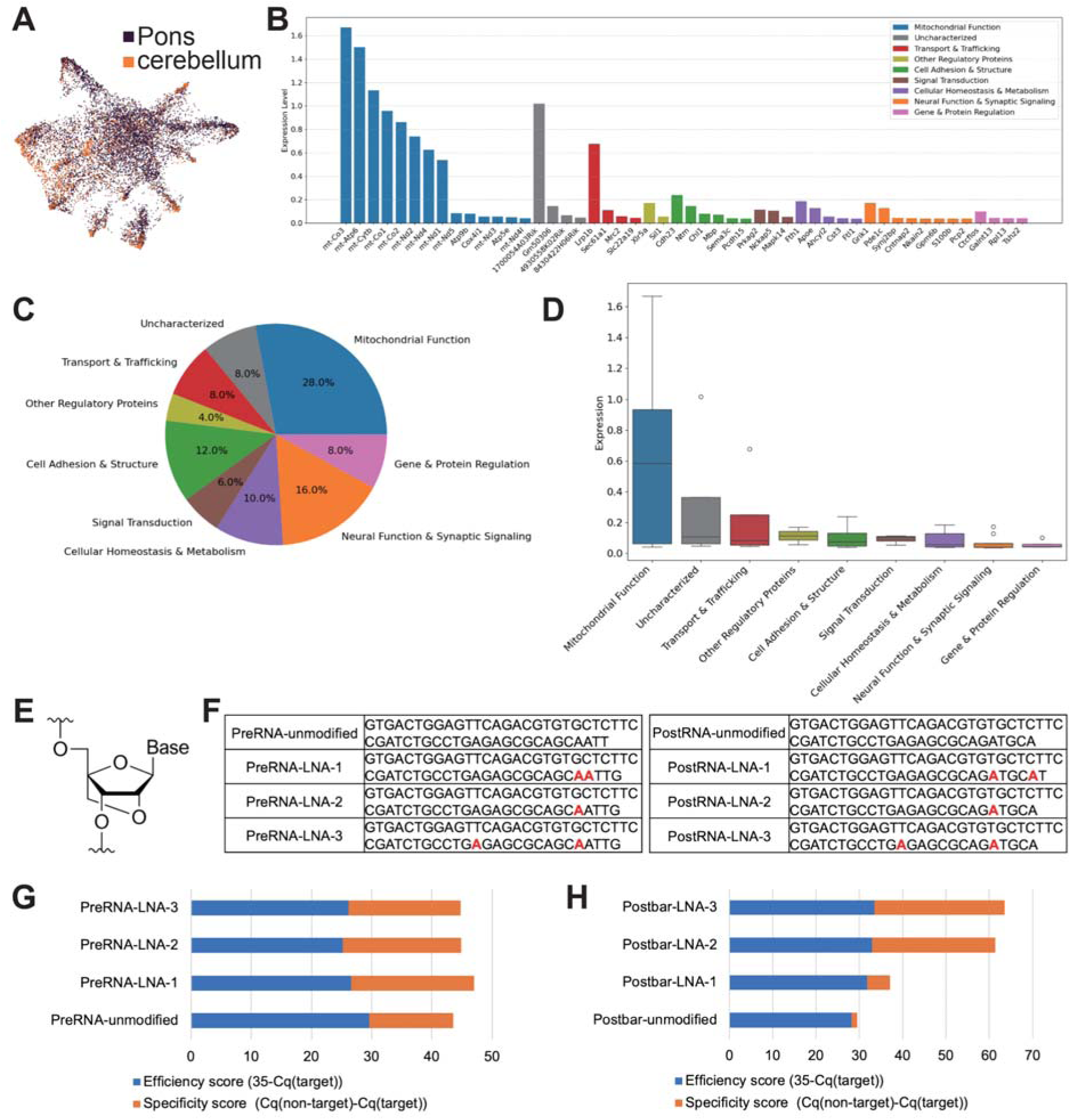
Analysis of single-synaptosome gene expression data and optimization of LNA-modified primer for balanced barcode amplification. (A) UMAP visualization of single-synaptosome RNA sequencing data from pons (purple) and cerebellum (orange). The extensive mixing of synaptosomes from different brain regions indicates insufficient gene detection sensitivity to distinguish synaptic origins based on transcriptome alone. (B) Expression levels of top 50 most abundant transcripts detected in synaptosomes, colored by gene category. Mitochondrial transcripts (blue) dominate the synaptosomal RNA pool, with limited representation of transport and synapse-related genes. (C) Proportion of different gene categories among top 50 expressed genes in synaptosomes. Mitochondrial genes comprise 28% of highly expressed transcripts. (D) Expression level distributions across gene categories, demonstrating higher expression of mitochondrial transcripts compared to other categories. Boxes show quartiles, whiskers extend to 1.5× interquartile range, and points indicate outliers. (E) Chemical structure of locked nucleic acid (LNA) nucleotide showing the methylene bridge between 2’ oxygen and 4’ carbon that “locks” the ribose in C3-endo conformation. (F) Sequence comparison of unmodified and LNA-modified primers for PreRNA (left) and PostRNA (right) amplification. LNA modifications (shown in red) were systematically introduced at different positions to optimize primer performance. Each version (LNA-1, LNA-2, LNA-3) represents different modification patterns. (G) Performance evaluation of PreRNA primers. Bar plots show efficiency score (35-Cq(target), blue) and specificity score (Cq(non-target)-Cq(target), orange) for unmodified and LNA-modified primers. Higher scores indicate better performance. PreRNA-LNA-1 was chosen as the optimized version. (H) Performance evaluation of PostRNA primers showing improved amplification efficiency and specificity with LNA modifications, particularly for LNA-2 and LNA-3 versions. Note the substantial improvement in specificity score compared to unmodified primers. PostRNA-LNA-3 was chosen as the optimized version.

**Extended Data Figure 5.**
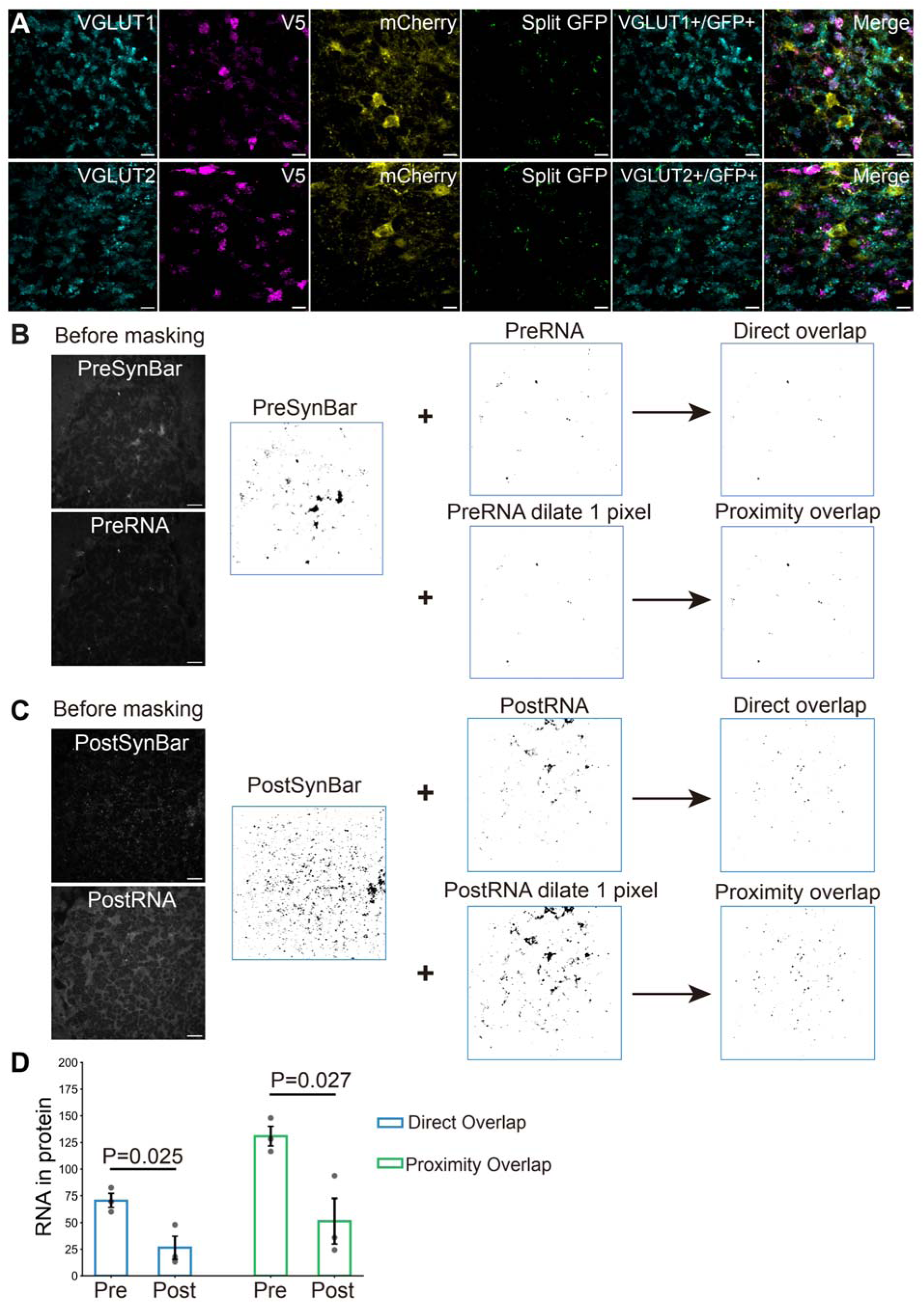
Validation of SynBar protein expression and RNA-protein colocalization. (A) Validation of PreSynBar expression with endogenous presynaptic markers, together with split GFP reconstitution. Upper panel, from left to right: VGLUT1 immunostaining (cyan), PreSynBar signal (anti-V5, magenta), PostSynBar signal (anti-mCherry, yellow), Split GFP signals marking connected synapses (green), overlay of VGLUT1 (cyan) and Split-GFP (green) to visualize spatial correspondence and merged image showing colocalization. Lower panel shows the same arrangement but with VGLUT2 instead of VGLUT1 in the leftmost channel. Representative max z projection field-of-view shown. Scale bar: 10 μm. (B-D) Quantitative analysis of RNA-protein colocalization. (B) Representative images showing the workflow for PreRNA-PreSynBar colocalization analysis. Binary masks were generated for RNA (FISH signal) and protein (immunofluorescence) channels from the raw images shown in the right panels. Direct overlap was calculated from the intersection of RNA and protein masks. Proximity overlap was determined after dilating the RNA mask by 1 pixel. (C) Similar analysis workflow for PostRNA-PostSynBar colocalization. (D) Quantification of RNA-protein colocalization (n=3 independent samples). Both direct overlap and proximity analysis showed significantly higher colocalization for PreRNA with PreSynBar compared to PostRNA with PostSynBar (direct overlap: p=0.025; proximity overlap: p=0.027, two-tailed t-test). Note the >100% value potentially came from the dilated signal from one RNA molecule being counted as associated with multiple nearby protein puncta.

**Extended Data Figure 6.**
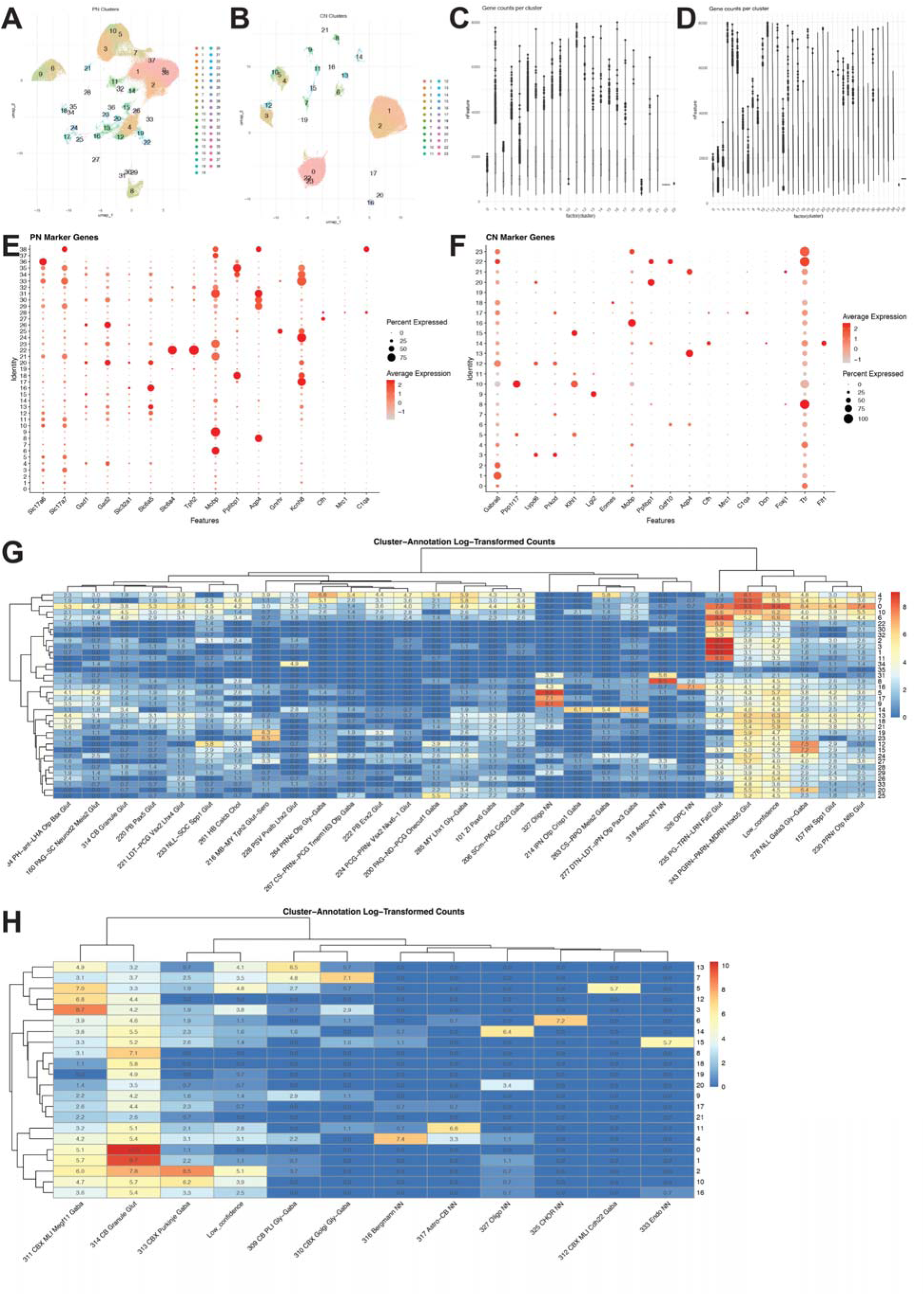
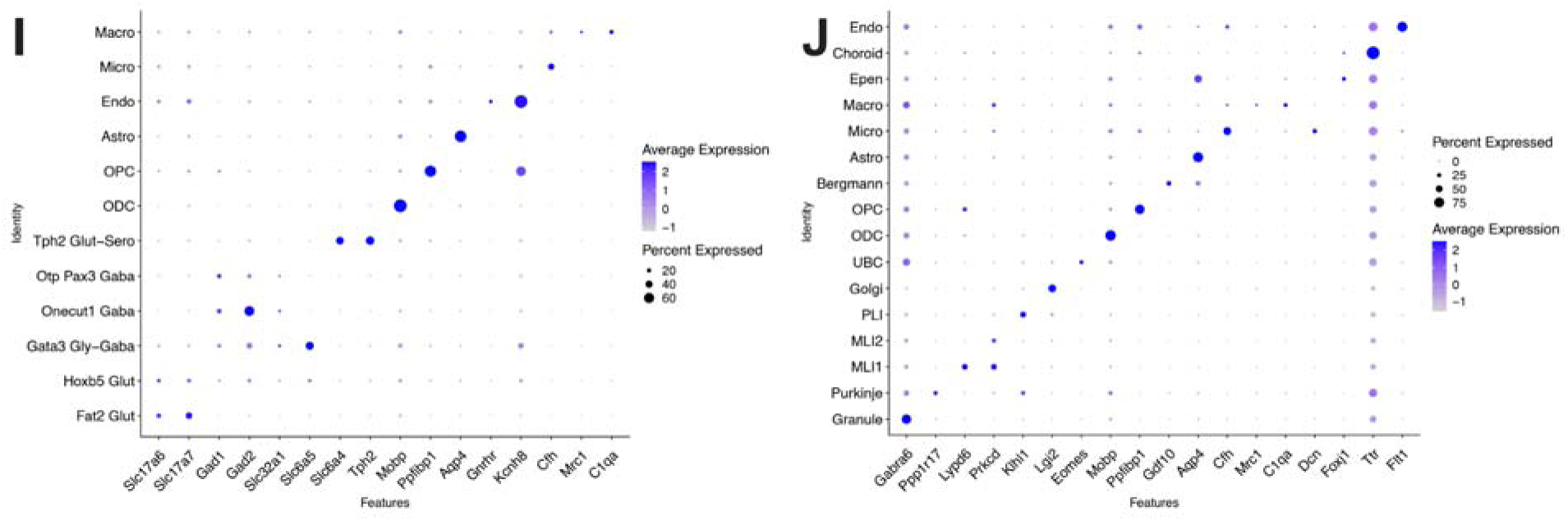
Cell type annotation of nuclei from pons and cerebellum. (A-B) Initial UMAP visualization of cell clusters in the pons (A) and cerebellum (B), with clusters numbered. (C-D) Violin plots showing gene counts per cluster, used to identify and remove low-quality clusters. Clusters 0, 2, 37, 38 were filtered out from pons (C), and clusters 0, 19, 22, 23 were removed from cerebellum (D) due to low gene counts (<1,000). (E-F) Dot plots for initial cell type annotation using canonical marker genes. For pons (E), these include neurotransmitter transporters (*Slc17a6, Slc17a7, Gad1, Gad2, Slc32a1, Slc6a5, Slc6a4*), synthesizing enzyme (*Tph2*), and non-neuronal cell markers. For cerebellum (F), markers include those for granule cells, interneurons (Golgi, UBC, MLI1/2, PLI), Purkinje cells, and non-neuronal cells. Dot size indicates percentage of expressing cells (20-80% for pons, 0-75% for cerebellum) and color intensity represents average expression level (−1 to 2). (G-H) Allen Institute’s MapMyCell annotation results shown as hierarchical clustering heatmaps for pons (G) and cerebellum (H), displaying the proportion of each annotated cell type within individual clusters to assign cluster identity. Values displayed in each entry of the box represent the log-transformed cell count with the predicted label. (I-J) Final dot plots showing expression of the same canonical marker genes across merged and annotated cell populations in pons (I) and cerebellum (J), confirming the consistency between marker gene expression and final cluster annotations. Dot size indicates percentage of expressing cells and color intensity shows average expression level.

**Extended Data Figure 7.**
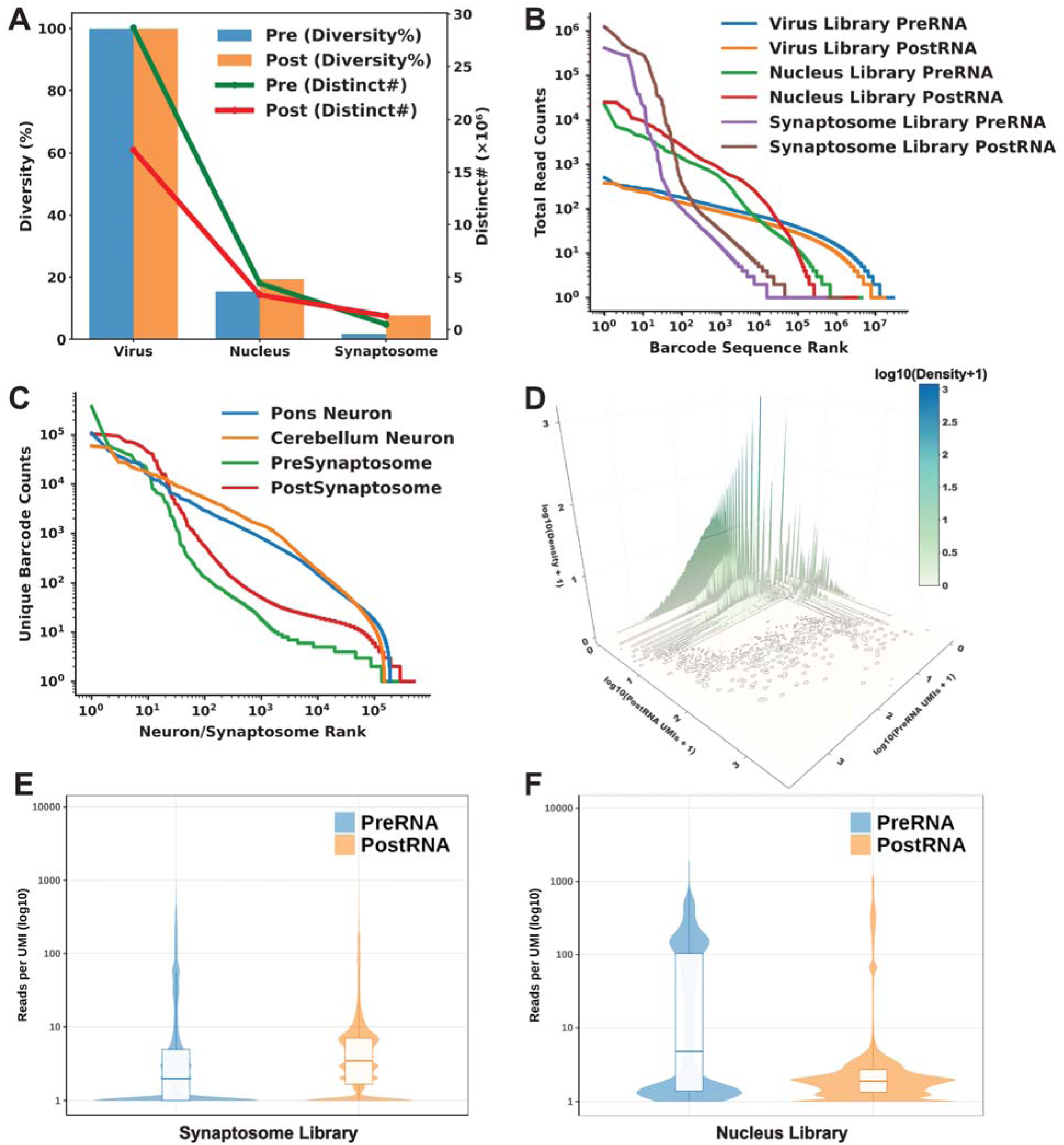
Analysis of Unbiased Features in Virus, Nuclei and Synaptosome Barcode Library. **(A)** Line and bar plots displaying Pre/PostRNA library diversity across virus, nuclei, and synaptosome barcode libraries, with PreRNA and PostRNA shown in blue and orange, respectively. The line plot shows the number of distinct Pre/PostRNA sequences determined for virus, nuclei, and synaptosome libraries (y-axis on the right). The bar plot shows relative diversity of the Pre/PostRNA libraries (virus library diversity set to 100%; nuclei and synaptosome diversity reported as the ratio of distinct barcode number relative to the virus library). **(B)** Rank–abundance plots of Pre/PostRNA total read counts (read count for virus and UMI count for nuclei and synaptosome) across virus, nuclei, and synaptosome barcode libraries. For each library, barcode sequences were ordered by descending abundance (rank 1 = most abundant) and plotted as sequence rank (x-axis) versus barcode total read count (y-axis). Both axes are on a log10 scale to highlight the heavy-tailed distribution. Curves correspond to Virus PreRNA Library (blue), Virus PostRNA Library (orange), Nucleus PreRNA Library (green), Nucleus PostRNA Library (red), Synaptosome PreRNA Library (purple), and Synaptosome PostRNA Library (brown). **(C)** Rank–abundance plots of distinct Pre/PostRNA sequences per neuron or per synaptosome in nuclei or synaptosome barcode libraries. Neurons or synaptosomes were ordered by descending count of distinct barcode sequences they contain (rank 1 = containing most distinct sequences) and plotted as neuron/synaptosome rank (x-axis) versus distinct barcode count (y-axis). Both axes are on a log10 scale to highlight the heavy-tailed distribution. Curves correspond to Pons Neuron (blue), Cerebellum Neuron (orange), PreSynaptosome (green), and PostSynaptosome (red). **(D)** 3D density surface plot comparing PreRNA and PostRNA UMI counts per synaptosome (log10 scales), where the x- and y-axes represent log10-transformed UMI counts for PreRNA and PostRNA, respectively, and the z-axis represents log10(density + 1). Here, density is the 2D kernel-density estimate of log10-transformed PreRNA and PostRNA UMI counts, representing the relative concentration of synaptosomes (barcode pairs) across the joint distribution of Pre/PostRNA. **(E–F)** Violin plots showing the distribution of reads per UMI for PreRNA (left) and PostRNA (right) in synaptosome (**E**) and nucleus (**F**) libraries. Box plots within the violins indicate median and quartile values, with extensions showing the full range excluding outliers.

**Extended Data Figure 8.**
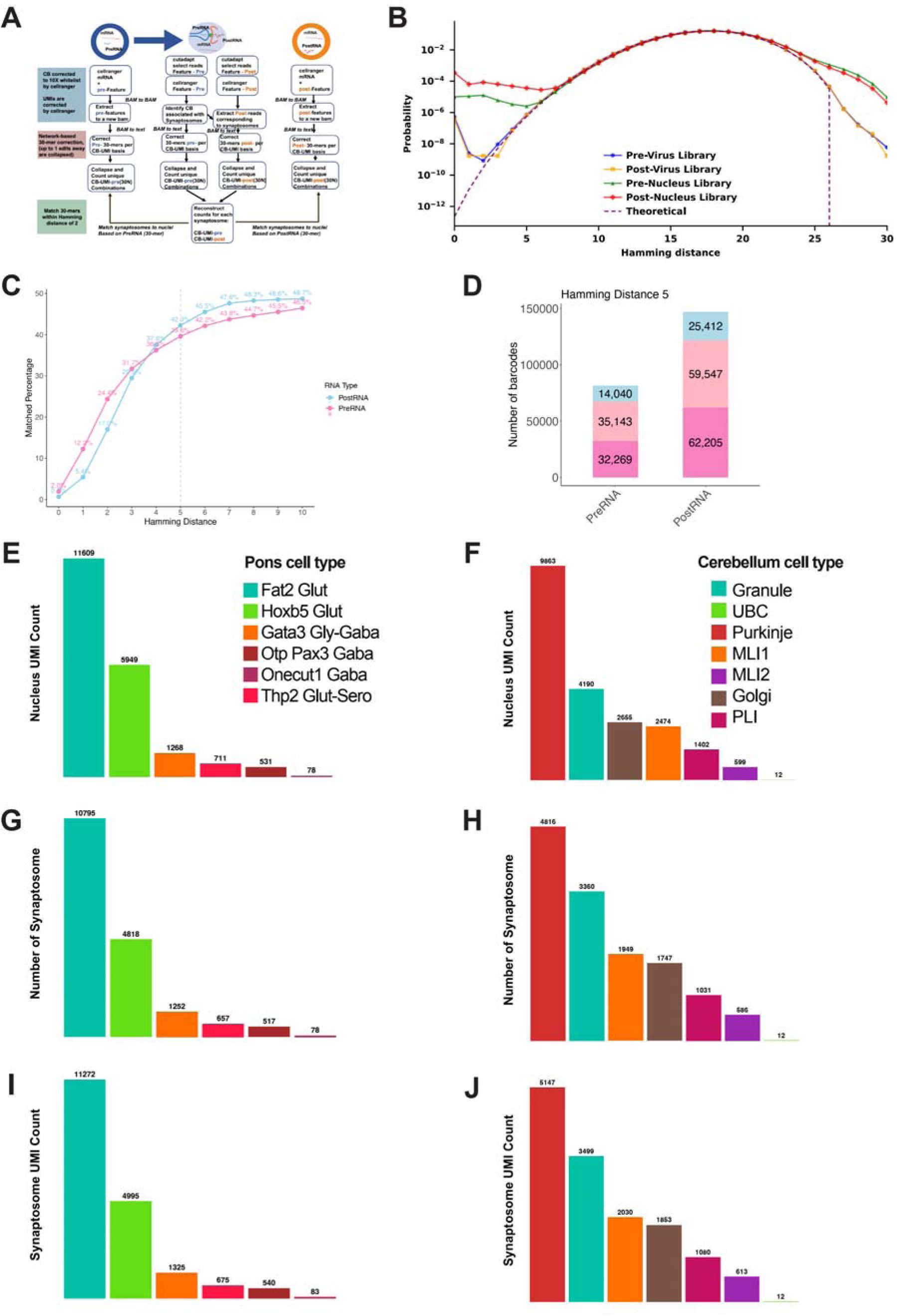
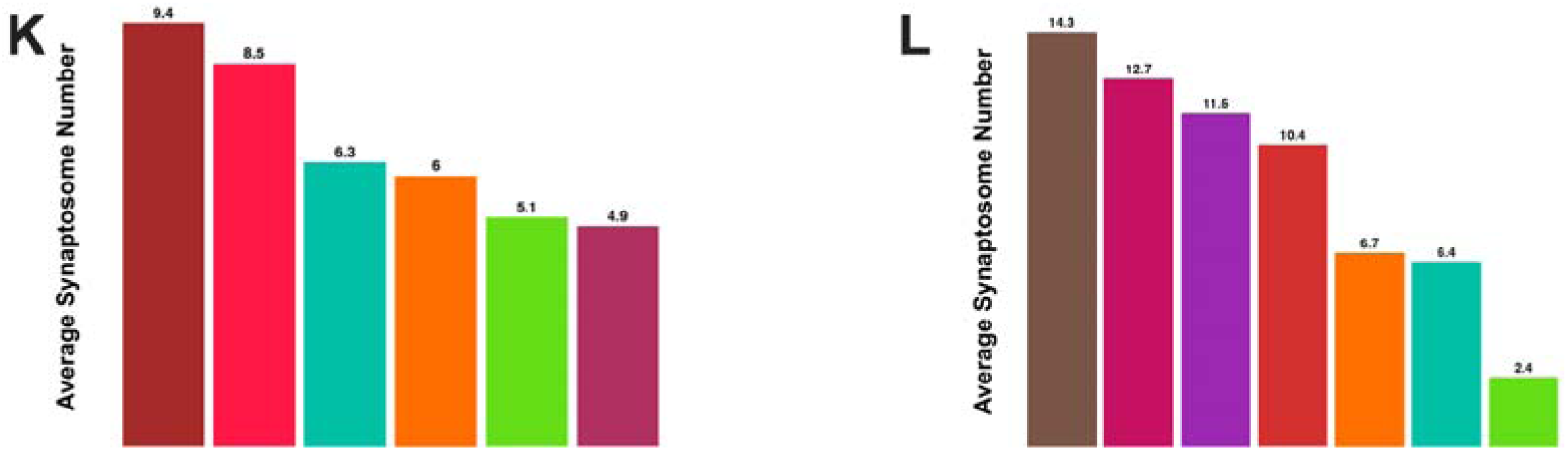
Computational pipeline and analysis for pontocerebellar connectome reconstruction. (A) Workflow of the computational pipeline showing parallel processing of nuclear mRNAs and barcodes. Key steps include CB-UMI-based error correction, feature extraction, network-based 30-mer correction, and barcode matching across compartments. (B) Barcode sequence pair editing distance distribution across virus and nucleus library. Curves show the probability distribution of pairwise Hamming distances for Virus PreRNA library (blue), Virus PostRNA library (orange), Nucleus PreRNA library (green), and Nucleus PostRNA library (red) libraries, compared to the expected theoretical distribution (purple dashed line). All values represent group means across replicates (n_boot=5). (C) Relationship between matching Hamming distance and sequence matching rate. Line plot showing the relationship between single matched percentage and Hamming distance for both PostRNA (blue) and PreRNA (pink). The dashed line indicates the chosen Hamming distance = 5 for subsequent analyses. (D) Quantification of matched and unmatched barcodes at chosen matching Hamming distances (5), shows the distribution between PostRNA and PreRNA, with barcodes matching with a unique nucleus (Single Match) in pink and unmatched in blue, while barcodes matching with multiple nuclear references (Multiple Matches) shown in light pink. Only Single Match barcodes were passed down to downstream analysis. (E-F) Distribution of total matched nucleus barcode UMI counts across pons (E) and cerebellum (F) cell types. Each barcode molecule from nucleus that matched to a synaptosome was counted, showing the total barcode coverage per cell type: in pons, Fat2 Glut (11,609 reads), Hoxb5 Glut (5,949 UMIs), Gata3 Gly-GABA (1,268 UMIs), Tph2 Glut-Sero (711 UMIs), Otp Pax3 Gaba (531 UMIs), and Onecut1 Gaba (78 UMIs); in cerebellum, Purkinje cells (9,863 UMIs), Granule cells (4,190 UMIs), Golgi cells (2,655 UMIs), MLI1 (2,474 UMIs), PLI (1,402 UMIs), MLI2 (599 UMIs), and UBC (12 UMIs). This represents the barcode UMI counts from the nuclei Pre/PostRNA that matched with synaptosome at Hamming distance 5. (G-H) Distribution of matched synaptosome number per cell type in pons (G) and cerebellum (H). Synaptosome number for each matched cell type: in pons, Fat2 Glut (n=10,795), Hoxb5 Glut (n=4,818), Gata3 Gly-GABA (n=1,252), Tph2 Glut-Sero (n=657), Otp Pax3 Gaba (n=517), and Onecut1 Gaba (n=78); in cerebellum, Purkinje cells (n=4,816), Granule cells (n=3,360), MLI1 (n=1949), Golgi cells (n=1,747), PLI (n=1,031), MLI2 (n=586), and UBC (n=12). (I-J) Distribution of total matched synaptosome barcode UMI counts across pons (I) and cerebellum (J) cell types. Each barcode molecule from synaptosomes that matched to a nucleus was counted, showing the total matched barcode presented in synaptosome: in pons, Fat2 Glut (11,272 UMIs), Hoxb5 Glut (4,995 UMIs), Gata3 Gly-GABA (1,325 UMIs), Tph2 Glut-Sero (675 UMIs), Otp Pax3 Gaba (540 UMIs), and Onecut1 Gaba (83 UMIs); in cerebellum, Purkinje cells (5,147 UMIs), Granule cells (3,499 UMIs), Golgi cells (2,030 UMIs), MLI1 (1,853 UMIs), PLI (1,080 UMIs), MLI2 (613 UMIs), and UBC (12 UMIs). This represents the barcode UMI counts from the synaptosomal Pre/PostRNA that matched with nuclei at Hamming distance 5. (K-L) Distribution of average synaptosome number per neuron across matched cell type in pons (K) and in cerebellum (L). In pons, Otp Pax3 Gaba (n=9.4), Tph2 Glut-Sero (n=8.5), Fat2 Glut (n=6.3), Gata3 Gly-GABA (n=6), Hoxb5 Glut (n=5.1), Onecut1 Gaba (n=4.9); in cerebellum, Golgi cells (n=14.3), PLI (n=12.7), MLI2 (n=11.5), Purkinje cells (n=10.4), MLI1 (n=6.7), Granule cells (n=6.4), UBC (n=2.4).

**Extended Data Figure 9.**
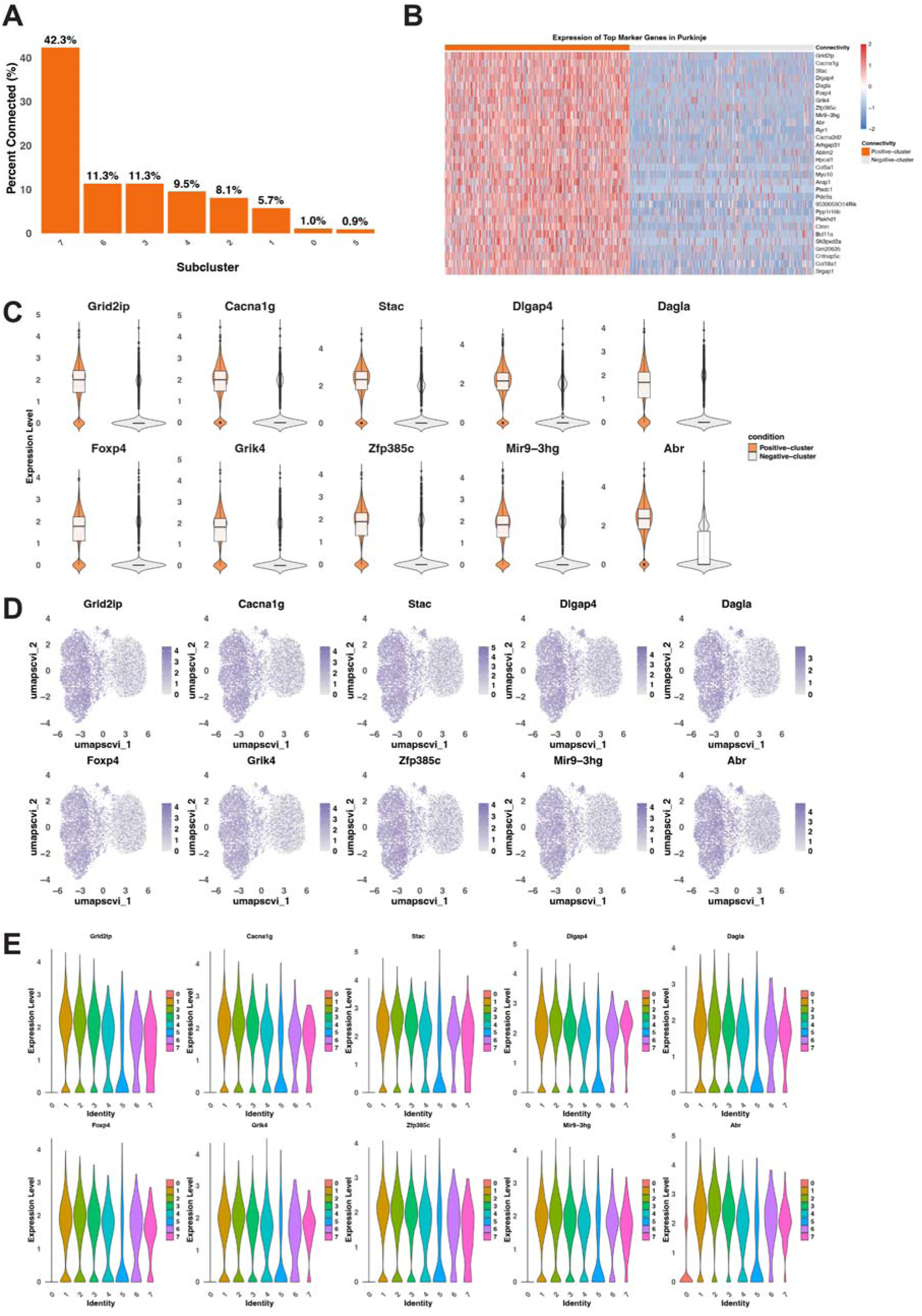
Detailed molecular analysis of pons-connected Purkinje cells. (**A**) Quantification of pons-connected Purkinje cells across different subclusters, showing highest enrichment in cluster 7 (42.3%), followed by cluster 6 (11.3%), with decreasing proportions in other clusters, demonstrating preferential connectivity in specific molecular subtypes. (B) Hierarchical clustering of expression patterns for top 30 marker genes in Positive-cluster (orange) versus Negative-cluster (gray) Purkinje cells. Color scale indicates normalized expression levels from −2 (blue) to 2 (red). (C) Two-condition violin plots showing expression distribution of top 10 marker genes in Positive-cluster (orange) versus Negative-cluster (gray) Purkinje cells, demonstrating consistent upregulation in connected populations. (D) UMAP feature plots showing expression patterns of top 10 marker genes across all Purkinje cells. Color intensity indicates expression level. (E) Detailed expression patterns of top 10 marker genes across all Purkinje cell subclusters (0-7). Each violin plot shows expression distribution within a subcluster, colored by cluster identity. Note the cluster-specific expression patterns correlating with connectivity.

**Extended Data Figure 10.**
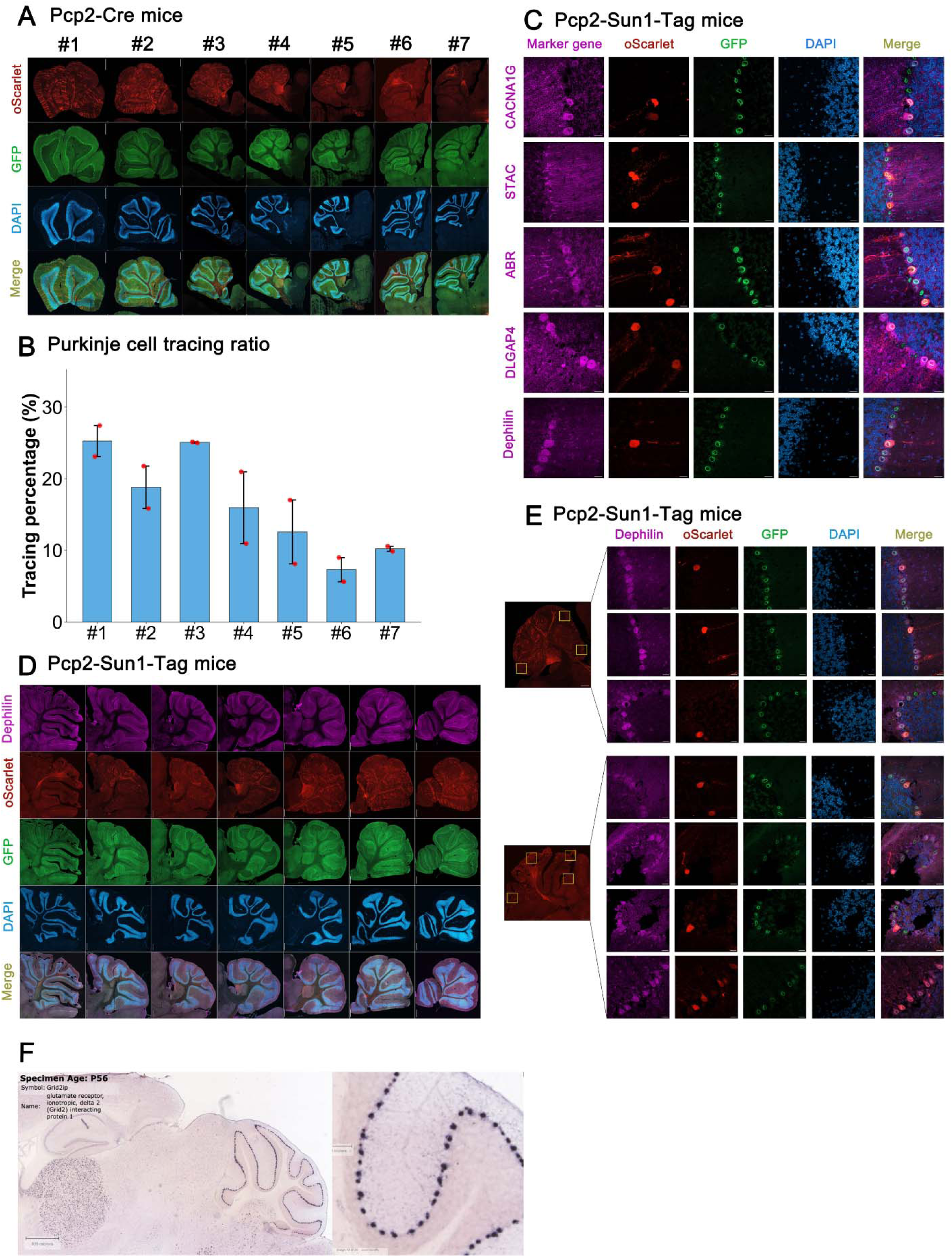
Detailed imaging analysis of pons-connected Purkinje cells. (A) Representative images from serial parasagittal mouse brain sections (from lateral to medial) showing consistent labeling of Purkinje cells by AAV1 anterograde tracing. For each panel from top to bottom: merged image showing colocalization, DAPI nuclear staining (blue), GFP-labeled Purkinje cell signal (green) and labeled pons-connected Purkinje cell signal (oScarlet, red). Scale bars: 400 μm. (B) Quantification of Purkinje cell tracing ratio by dividing oScarlet-positive cell numbers (pons-connected Purkinje cell) by GFP-positive cell numbers (total Purkinje cell). Each bar shows this ratio in corresponding parasagittal mouse brain sections from panel (A). (C) High-magnification images, ordered from top to bottom, display immunofluorescence validation of CACNA1G, STAC, ABR, DLGAP4, and *Grid2ip*/Delphilin. Marker gene expression immunostaining (magenta), labeled pontine-connected Purkinje cells (oScarlet, red), GFP-labeled Purkinje cell signal (green), DAPI nuclear staining (blue), and merged image showing colocalization. Scale bars: 400 μm (overview), 20 μm (high-magnification). (D-E) Further immunofluorescence validation of marker gene expression. Panels D (overview) and E (high magnification) show immunofluorescence validation of *Grid2ip*/Delphilin in serial parasagittal mouse brain sections, progressing from lateral to medial. (F) *Grid2ip* RNA in situ hybridization data from the Allen Brain Atlas (P56 mouse). Left: Low magnification image of sagittal cerebellar section. Right: High magnification of boxed region showing *Grid2ip* expression in a subset of Purkinje cells, providing independent evidence of restricted expression pattern.

**Table S1.** Related to Methods. Plasmids used or generated in this study.

| Name | Vector | Promoter | Description |
| --- | --- | --- | --- |
| hSyn-PreSynBar-hSyn-PreRNA barcode landing pad | pAAV | hSyn | SynBar protein and RNA construct used in HEK and neuron culture experiment, related to Fig. 1 |
| hSyn-PostSynBar-hSyn-PostRNA barcode landing pad | pAAV | hSyn | SynBar protein and RNA construct used in HEK and neuron culture experiment, related to Fig. 1 |
| hU6-presynaptic-barcode-3BoxB | pAAV | hU6 | hU6 promoter driven barcode RNA with no polyA, related to Extended Data Fig. 1C |
| hU6-postsynaptic-barcode-3BoxB | pAAV | hU6 | hU6 promoter driven barcode RNA with no polyA, related to Extended Data Fig. 1C |
| CAG- PreRNA-PPIB-original-hSyn-Cre | pAAV | CAG and hSyn | PreRNA with different cis-element to potentially improve its synaptic localization, related to Extended Data Fig. 1D |
| CAG- PreRNA-PPIB-5' RPL7-hSyn-Cre | pAAV | CAG and hSyn | PreRNA with different cis-element to potentially improve its synaptic localization, related to Extended Data Fig. 1D |
| CAG- PreRNA-PPIB-5' RPL28-hSyn-Cre | pAAV | CAG and hSyn | PreRNA with different cis-element to potentially improve its synaptic localization, related to Extended Data Fig. 1D |
| CAG- PreRNA-PPIB-3' Trak2-hSyn-Cre | pAAV | CAG and hSyn | PreRNA with different cis-element to potentially improve its synaptic localization, related to Extended Data Fig. 1D |
| CAG- PreRNA-PPIB-3' Net1-hSyn-Cre | pAAV | CAG and hSyn | PreRNA with different cis-element to potentially improve its synaptic localization, related to Extended Data Fig. 1D |
| CAG- PreRNA-PPIB-5' RPL28-3' Trak2-hSyn-Cre | pAAV | CAG and hSyn | PreRNA with different cis-element to potentially improve its synaptic localization, related to Extended Data Fig. 1D |
| pre-SynView | pAAV | hSyn | Original pre-SynView construct cloned into pAAV vector, related to Extended Data Fig. 1E |
| post-SynView | pAAV | hSyn | Original post-SynView construct cloned into pAAV vector, related to Extended Data Fig. 1E |
| hSyn-Cre-hSyn- | pAAV | hSyn | The set of RNA constructs used in the optimized |
| PreRNA<br>barcode landing<br>pad (PPIB) |  |  | connectome-seq experiments, design shown in Fig. 1F. Also used for Basescope experiment shown in Fig. 4F |
| hSyn-PreSynBar-hSyn-PreRNA<br>barcode landing<br>pad | pAAV | hSyn | The set of SynBar constructs used in the optimized connectome-seq experiments, design shown in Fig. 1F |
| hSyn-Cre-hSyn-PostRNA<br>barcode landing<br>pad (dapB) | pAAV | hSyn | The set of RNA constructs used in the optimized connectome-seq experiments, design shown in Fig. 1F. Also used for Basescope experiment shown in Fig. 4F |
| hSyn-DIO-PostSynBar-P2A-mCherry | pAAV | hSyn | The set of SynBar constructs used in the optimized connectome-seq experiments, design shown in Fig. 1F |
| hSyn-DIO-axonTag-mGreenLantern<br>-P22N21 | pAAV | hSyn | Axon-targeted GFP with RNA binding domain, design shown in Extended Data Fig. 3A |
| DIO-dendriteTag-oScarlet3-P22N21 | pAAV | hSyn | Dendrite-targeted RFP with RNA binding domain, design shown in Extended Data Fig. 3A |
| sCAG-DIO-oScarlet3 | pAAV | sCAG | Used for AAV1 tracing experiment, design shown in Fig. 6F |
| AAV1 capsid | N/A | N/A | Used for AAV generation, Addgene #112862 |
| AAV2 capsid | N/A | N/A | Used for AAV generation, Addgene #104963 |
| pAdDeltaF6 | N/A | N/A | Used for AAV generation, Addgene #112867 |

**Table S2.**
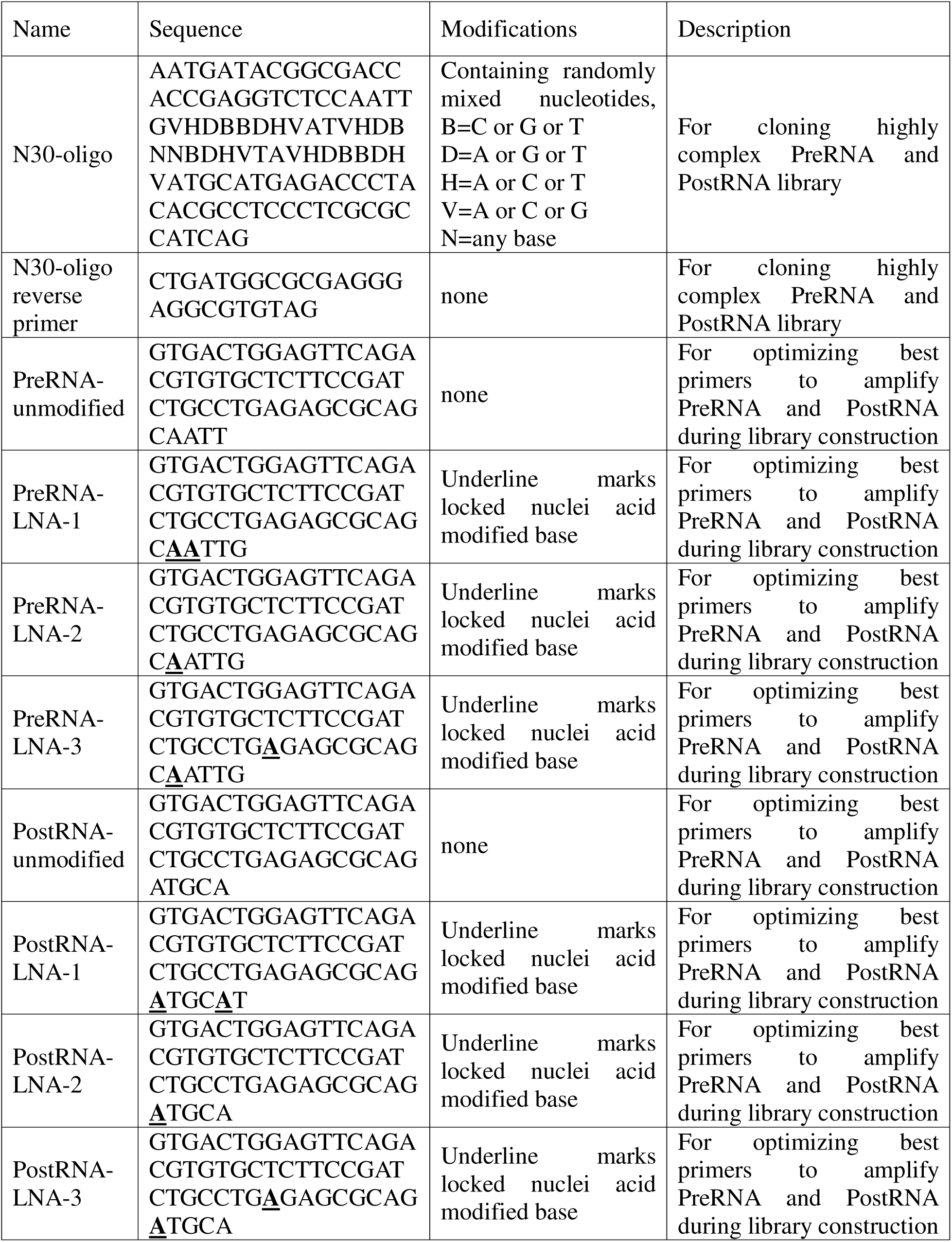
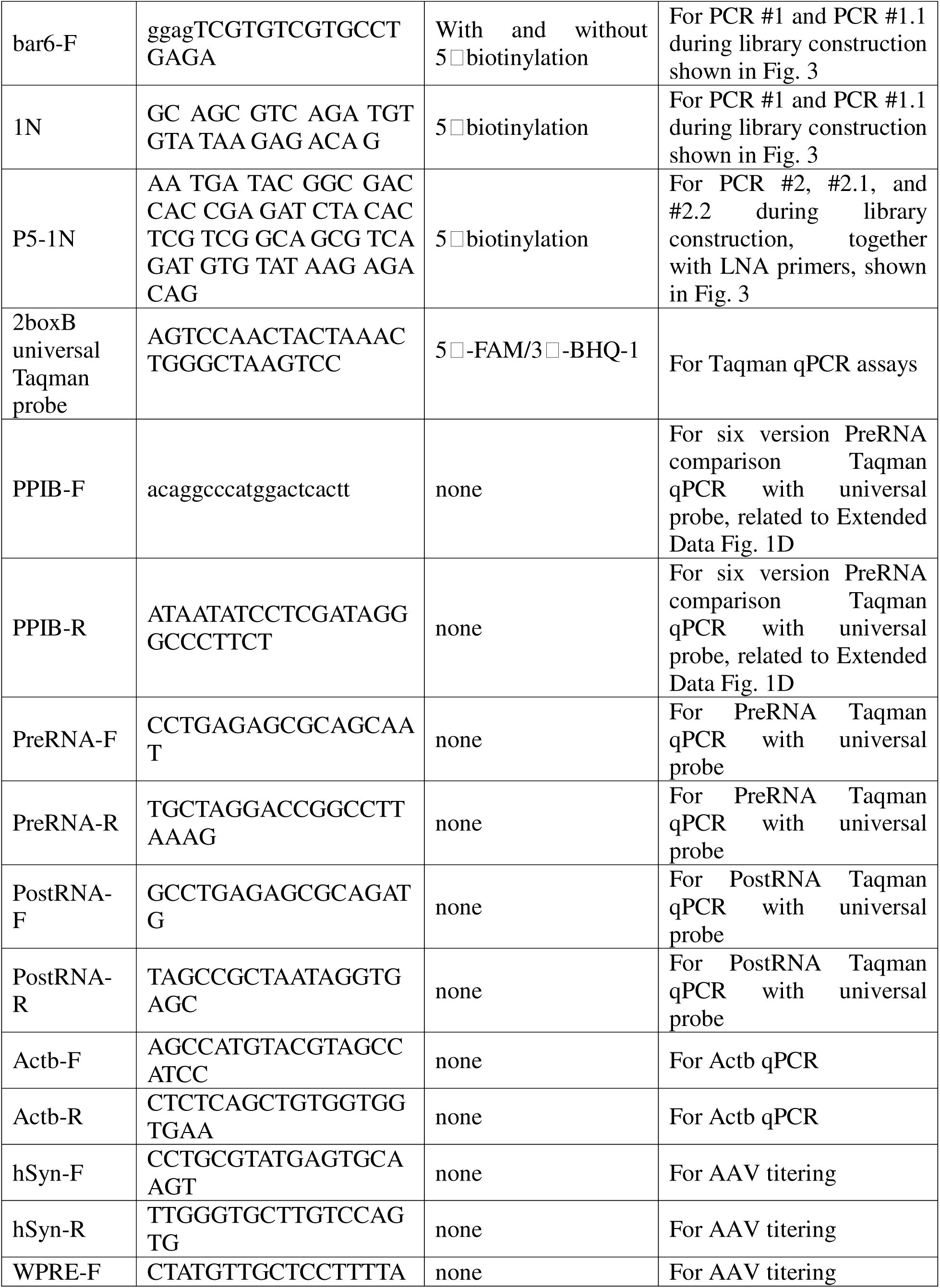
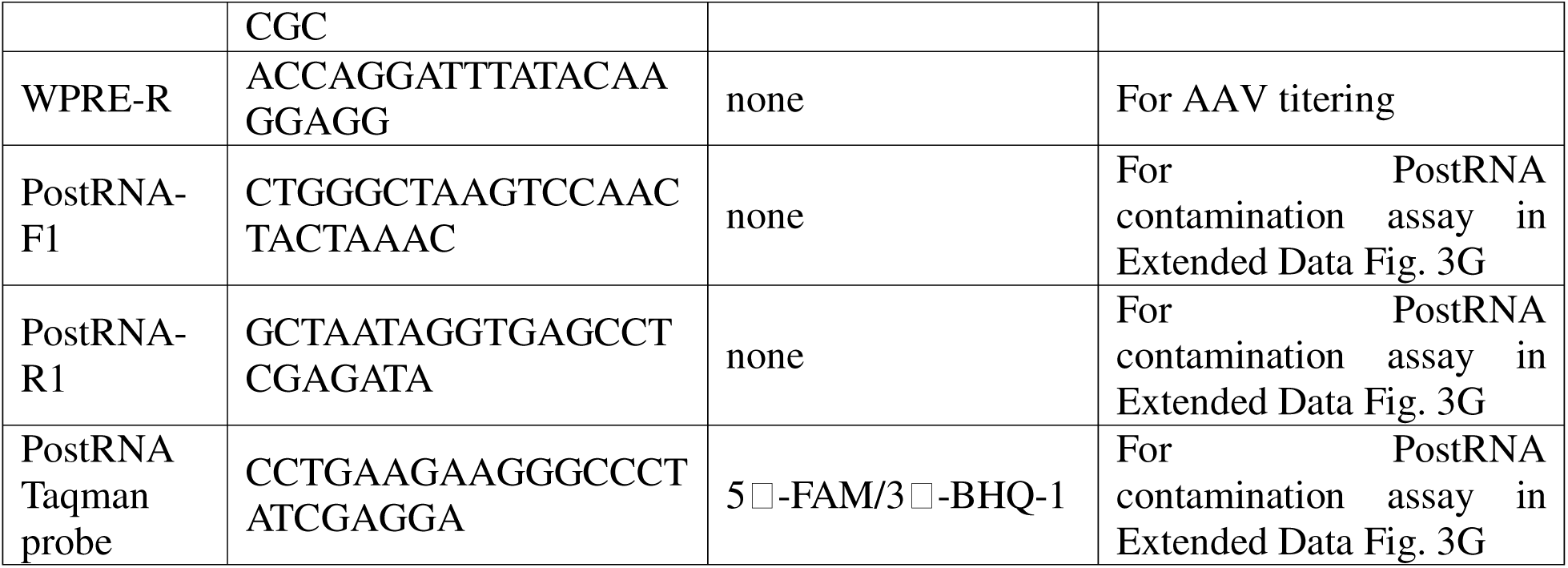
Related to Methods. Oligos used in the paper.

**Table S3.** Related to Methods. AAV used for different experiments.

| Name | Capsid | Titer (vg/ul) | Description |
| --- | --- | --- | --- |
| hSyn-PreSynBar-hSyn-PreRNA barcode landing pad | AAV1/2 mix | 3.98E+09 | For neuron culture infection, related to Fig. 1 |
| hSyn-PostSynBar-hSyn-PostRNA barcode landing pad | AAV1/2 mix | 9.13E+08 | For neuron culture infection, related to Fig. 1 |
| CAG-PreRNA-PPIB-original-hSyn-Cre | AAV1/2 mix | 4.55E+11 | For mouse brain injection to optimize PreRNA, related to Extended Data Fig. 1D |
| CAG-PreRNA-PPIB-5'RPL7-hSyn-Cre | AAV1/2 mix | 1.30E+11 | For mouse brain injection to optimize PreRNA, related to Extended Data Fig. 1D |
| CAG-PreRNA-PPIB-5'RPL28-hSyn-Cre | AAV1/2 mix | 3.66E+11 | For mouse brain injection to optimize PreRNA, related to Extended Data Fig. 1D |
| CAG-PreRNA-PPIB-3'Trak2-hSyn-Cre | AAV1/2 mix | 1.92E+11 | For mouse brain injection to optimize PreRNA, related to Extended Data Fig. 1D |
| CAG-PreRNA-PPIB-3'Net1-hSyn-Cre | AAV1/2 mix | 2.33E+11 | For mouse brain injection to optimize PreRNA, related to Extended Data Fig. 1D |
| CAG-PreRNA-PPIB-5'RPL28-3'Trak2-hSyn-Cre | AAV1/2 mix | 1.27E+11 | For mouse brain injection to optimize PreRNA, related to Extended Data Fig. 1D |
| hSyn-Cre-hSyn-PreRNA-library | AAV1/2 mix | 2.20E+9 | The set of SynBar viruses used for sequencing experiments, design shown in Fig. 1F |
| hSyn-PreSynBar-hSyn-PreRNA-library | AAV1/2 mix | 7.88E+9 | The set of SynBar viruses used for sequencing experiments, design shown in Fig. 1F |
| hSyn-Cre-hSyn-PostRNA barcode | AAV1/2 mix | 2.79E+9 | The set of SynBar viruses used for sequencing experiments, design shown in Fig. 1F |
| hSyn-DIO-PostSynBar-P2A-mCherry | AAV1/2 mix | 4.32E+10 | The set of SynBar viruses used for sequencing experiments, design shown in Fig. 1F |
| hSyn-DIO-axonTag-mGreenLantern-P22N21 | AAV1/2 mix | 1.74E+10 | Axon-targeted GFP with RNA binding domain, design shown in Fig. 1F |
| DIO-dendriteTag-oScarlet3-P22N21 | AAV1/2 mix | 1.93E+10 | Dendrite-targeted RFP with RNA binding domain, design shown in Fig. 1F |
| hSyn-PreSynBar-hSyn-PreRNA-PPIB | AAV1/2 mix | 4.51E+09 | For mouse brain injection for Basescope experiment shown in Fig. 4F |
| hSyn-Cre-W3SL-hSyn-PostRNA-dapB | AAV1/2 mix | 1.97E+10 | For mouse brain injection for Basescope experiment shown in Fig. 4F |
| sCAG-DIO-oScarlet3 | AAV1 | 1.42E+11 | Used for AAV1 tracing experiment, design shown in Fig. 6F |
| hSyn-Cre | AAV1 | 1.00E+10 | Used for AAV1 tracing experiment |

